# Enhanced genetic analysis of type 1 diabetes by selecting variants on both effect size and significance, and by integration with autoimmune thyroid disease

**DOI:** 10.1101/2021.02.05.429962

**Authors:** Daniel J. M. Crouch, Jamie R.J. Inshaw, Catherine C. Robertson, Jia-Yuan Zhang, Wei-Min Chen, Suna Onengut-Gumuscu, Antony J. Cutler, Carlo Sidore, Francesco Cucca, Flemming Pociot, Patrick Concannon, Stephen S. Rich, John A. Todd

**Affiliations:** JDRF/Wellcome Diabetes and Inflammation Laboratory, Wellcome Centre for Human Genetics, Nuffield Department of Medicine, NIHR Oxford Biomedical Research Centre, University of Oxford, Oxford, United Kingdom; Center for Public Health Genomics, University of Virginia, Charlottesville, Virginia, USA; Department of Public Health Sciences, University of Virginia, Charlottesville, Virginia, USA; Institute for Research in Genetics and Biomedicine (IRGB), Sardinia, Italy; Department of Pediatrics, Herlev University Hospital, Copenhagen, Denmark; Institute of Clinical Medicine, Faculty of Health and Medical Sciences, University of Copenhagen, Copenhagen, Denmark; Type 1 Diabetes Biology, Department of Clinical Research, Steno Diabetes Center Copenhagen, Gentofte, Denmark; Department of Pathology, Immunology, and Laboratory Medicine, University of Florida, Gainesville, Florida, USA; Genetics Institute, University of Florida, Gainesville, Florida, USA

## Abstract

For polygenic traits, associations with genetic variants can be detected over many chromosome regions, owing to the availability of large sample sizes. Most variants, however, have small effects on disease risk and, therefore, unravelling the causal variants, target genes, and biology of these variants is challenging. Here, we define the Bigger or False Discovery Rate (BFDR) as the probability that either a variant is a false-positive or a randomly drawn, true-positive association exceeds it in effect size. Using the BFDR, we identified 302 previously unreported signals with larger effect associations with type 1 diabetes and autoimmune thyroid disease. Out of 239 genome-wide significant signals in both diseases, only 66 (28%) show evidence for having a large effect using the BFDR, further demonstrating how using a combination of effect size and significance, rather than significance alone, is important in identifying SNPs and candidate genes for further investigation.

## Introduction

Most genome-wide association study (GWAS) associations have small effects on phenotypes, with the genetic risk for common diseases distributed across hundreds of loci, mostly over common alleles^1^. Interpretation of the biological effects of these variants, and establishment of which are causal, either statistically (fine-mapping) or experimentally, requires considerable effort and the results may provide limited or non-actionable mechanistic insight. In type 1 diabetes (T1D), for example, HLA, *PTPN22, INS* and *IL2RA* have variants with alleles with relatively large effects on risk (Odds Ratios (ORs)>1.3), and it is these loci that have yielded most biological insights so far^2–8^, subsequently taken forward to translation and clinical trials^9^. As sample sizes increase even further, for example from the availability of data from large biobanks, loci with P values that pass the established threshold for significance, P < 5 × 10^−8^, have either lower effect sizes or lower MAFs, the latter of which are arguably preferable for follow-up. An alternative to this typical genome-wide significance threshold is the false discovery rate (FDR)^10–12^, an estimate of the probability that associations beyond a specified P value threshold are false (e.g., at FDR=0.01, 1% of associations are likely to be false).

Analyses using FDR may yield many more small-effect variants associated with a trait than P<5×10^−8^, providing motivation to develop a method that priorities variants based on both significance and effect size. We define the Bigger or False Discovery Rate (BFDR) as the probability that either a variant is a false positive or that a randomly chosen true association exceeds it in effect size. Variables with BFDR of, for example, 1% therefore warrant greater interest than those with FDR of 1%, as they are likely to be both true positives and to exceed most other true positives in effect size. Whereas FDR=5% may be deemed too lenient for declaring an association in many settings, BFDR=5% implies both FDR<5% and a larger than expected effect, and so a BFDR<5% threshold could be used to select large effect variants that would be normally missed by a more stringent FDR threshold (e.g. 1%). This is based on the understanding that the overall cost of following up an association is a combination of a) its probability of being false and b) its probability of having an unremarkable effect size compared to other associations. In addition to this strategy, we use a more stringent 1% BFDR threshold to obtain a set of associations that are very likely to be both true positives and to have large effects. Our novel summary-stats method for BFDR estimation based around a technique we call ‘prior splitting’ (SI appendix) is available for download as an R package, priorsplitteR (see code availability).

Genetic variation underlying susceptibility to T1D was revealed initially in candidate gene studies, then GWAS followed by fine-mapping using the custom SNP array, ImmunoChip^13–19^, identifying 98 genome-wide chromosome regions associated with disease risk (P<5×10^−8^). Here, we apply the BFDR to discover new regions affecting T1D risk with larger effects. We performed the largest GWAS meta-analysis to date (15,573 cases), combining two UK case-control datasets, Sardinian and Finnish case-control datasets, and families with affected offspring from the Type 1 Diabetes Genetics Consortium (T1DGC). Using SNPs outside of the HLA region, we identified 89 independently-associated susceptibility regions satisfying BFDR<5% (40 not previously reported), of which 25 would have been missed by a more conventional FDR threshold of 1%. We found further associations for 130 regions satisfying FDR<1% but with BFDR≥5%, which we suggest are likely to be true associations but with risk effect sizes that are difficult to follow up in downstream experiments. We found only 16 regions with BFDR<1% (which by definition also have FDR<1%) indicating that even in GWAS that are well-powered to detect significant associations, finding the loci with the conclusively largest effects remains challenging.

Common autoimmune diseases share many loci across the genome, in addition to HLA, and this overlap in genetic risk and immune pathology offers the opportunity to conduct joint analyses that may increase statistical power^20^. Since autoimmune thyroid disease (ATD) is very common in the UK Biobank (UKBB), with approximately 30,000 cases out of 500,000 participants, we used the BFDR and the UKBB GWAS data to map ATD associations with larger-than-average effects. We found 338 regions with BFDR<5%, of which 158 had FDR≥1%. We also found 692 associated with FDR<1% but with smaller effects (BFDR≥5%). Using colocalisation analysis, we found several T1D-associated regions that showed no evidence of effects on ATD nor with other immune diseases, suggesting that they may have roles outside of the immune system, indicative of the distinct organ-specific targets of the two diseases, i.e. the pancreatic islet beta cells (T1D) and thyroid gland (ATD). We drew on the higher power of the ATD dataset, together with pleiotropic effects of ATD variants on T1D, to identify 105 additional T1D regions (76 not previously reported) with either BFDR<5% or FDR<1% that were also ATD-associated.

## Results

### Bigger or False Discovery Rate (BFDR)

We searched for variants which, as is the case for *PTPN22, INS* and the HLA region, have large effects on T1D risk, but which might have been previously overlooked due to their low frequencies (e.g. below 5%), leading to lower statistical power for detection. For this purpose we define the BFDR, for a given variant, as the overall probability that either a) the SNP is a false positive association or b) the SNP is a true positive but randomly choosing from the distribution of true associations produces a larger effect size. Informally, the BFDR for SNP *i* is:

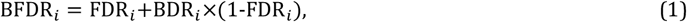

where,

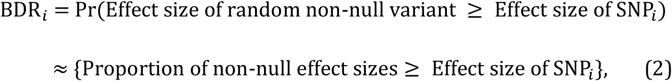

where BDR stands for ‘Bigger Discovery Rate’. The BFDR is upper bounded by the FDR, so that all BFDRs below e.g. 1%will also have FDR<1%, and therefore applies further stringency (in terms of sizes of effects) to a set of associations passing the same FDR threshold. Here, we explore associations passing either BFDR<1% or FDR<1%. It may be admissible to accept a higher error rate in order to reject the two separate hypotheses tested by the BFDR, so we also explore associations with a more lenient BFDR<5%.

We developed an empirical Bayesian method for estimating the BFDR without the need to explicitly model prior distributions of SNP effects (SI appendix). Our method does not account for dependence between SNPs, but we apply it to GWAS data containing LD structure as standard methods of FDR control remain accurate under positive dependency between test statistics^21^. The method was implemented in R and requires several hours computing time for GWAS-scale data.

To test the method, we randomly simulated 100 effect-size distributions representing different genetic architectures, before simulating true and estimated effect sizes for 100,000 SNPs, representing a GWAS, from each architecture. In a given simulated architecture, the extent to which effect sizes varied with allele frequency was randomly decided, so that rare SNPs might have a similar or much more variable true effect size distribution compared to common SNPs. Half of the true effect sizes were zero, i.e. true null effects, in all architectures. We ran BFDR estimation separately on each simulated GWAS, returning estimates of BFDR, BDR and FDR for each SNP. We thresholded SNPs based on these three quantities, at several levels of error control (α), and then computed the empirical error rates for SNPs passing each threshold, using the known true effects (Figure 1a). SNPs with BFDRs passing a given a threshold had, on average, empirical error rates lower than a, indicating good error control, and we found the equivalent was true for the BDR and FDR components of the BFDR (Figure 1b-c).

**Figure 1.**
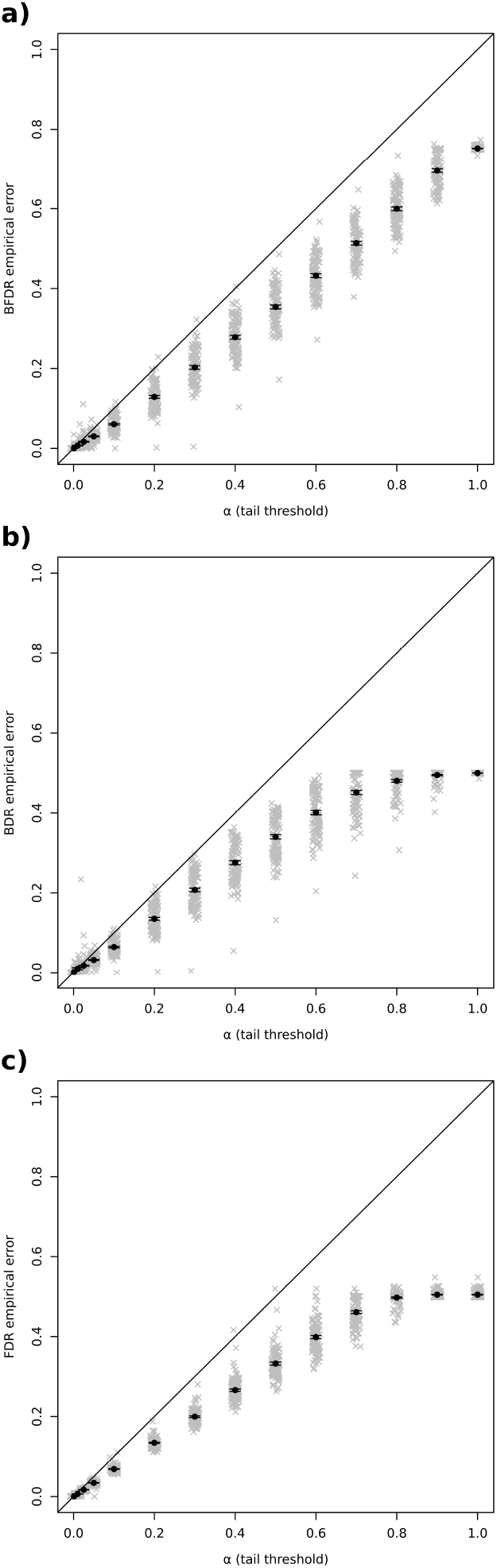
Average error rates of a) BFDR, b) BDR and c) FDR estimates for 100 GWAS simulations at 14 α thresholds (10^−3^, 10^−2^, 2.5×10^−2^, 5’10^−2^, and 0.1 to 1 in increments of 0.1) in grey, with means and standard errors of the grey points shown in black. Means and upper error bars fell below the 1:1 line, indicating good error control. FDR empirical error (i.e. the proportion of null SNPs passing the threshold) approached 0.5 as α was increased to 1, as half of the SNPs in our simulations had null effects of zero. BDR empirical error was defined as the proportion of non-null SNPs with larger true effects than each threshold-passing SNP, averaged over the threshold-passing SNPs (see online methods). This approached 0.5 as α was increased to 1, as this is the maximum value it can take. As the BFDR is a function of these two values (see Equation 1), its error rate approached 0.75 as α was increased.

### T1D GWAS meta-analysis

Imputation, QC and GWAS analysis was performed on Illumina (Infinium 550K, 3983 cases) and Affymetrix (GeneChip 500K, 1926 cases) genotyped UK samples, and on samples from Sardinia (Affymetrix 6.0 and Illumina Omni Express, 1558 cases). Affected-offspring trios from T1DGC were genotyped on the Illumina Human Core Exome beadchip and analysed with the TDT test (3173 trios) after imputation and QC. Results from the four cohorts were meta-analysed under the additive model, together with FinnGen (4933 cases).

LD-based filtering found 219 independent T1D-associated signals (Figures 2a, 3a and S3a, Table S1a), 194 of which satisfied FDR<1% for the lead variant in the signal, with the remaining 25 having BFDR<5% in spite of having FDR≥1% (we use ‘independent signals’ to refer to a disease-associated variant or set of variants in LD, related to, but distinct, from a physical region). These 25 variants would have been missed by using an FDR threshold of 1%, despite having a median OR for the risk variant (OR_risk_) of 1.23, (median MAF=2.9%, all T1D MAFs for controls only) and by definition of the BFDR have FDR<5%. The 89 total variants satisfying BFDR<5% had median OR_risk_=1.18 (median MAF=7.7%) versus 1.10 for the 194 with FDR<1% (median MAF=26.7%), indicating the shift in average effect sizes and MAF using BFDR. Sixteen SNPs satisfying a stringent BFDR of 1% had median OR_risk_=1.27 and median MAF= 5.9%, shown in Table 1a.

**Figure 2.**
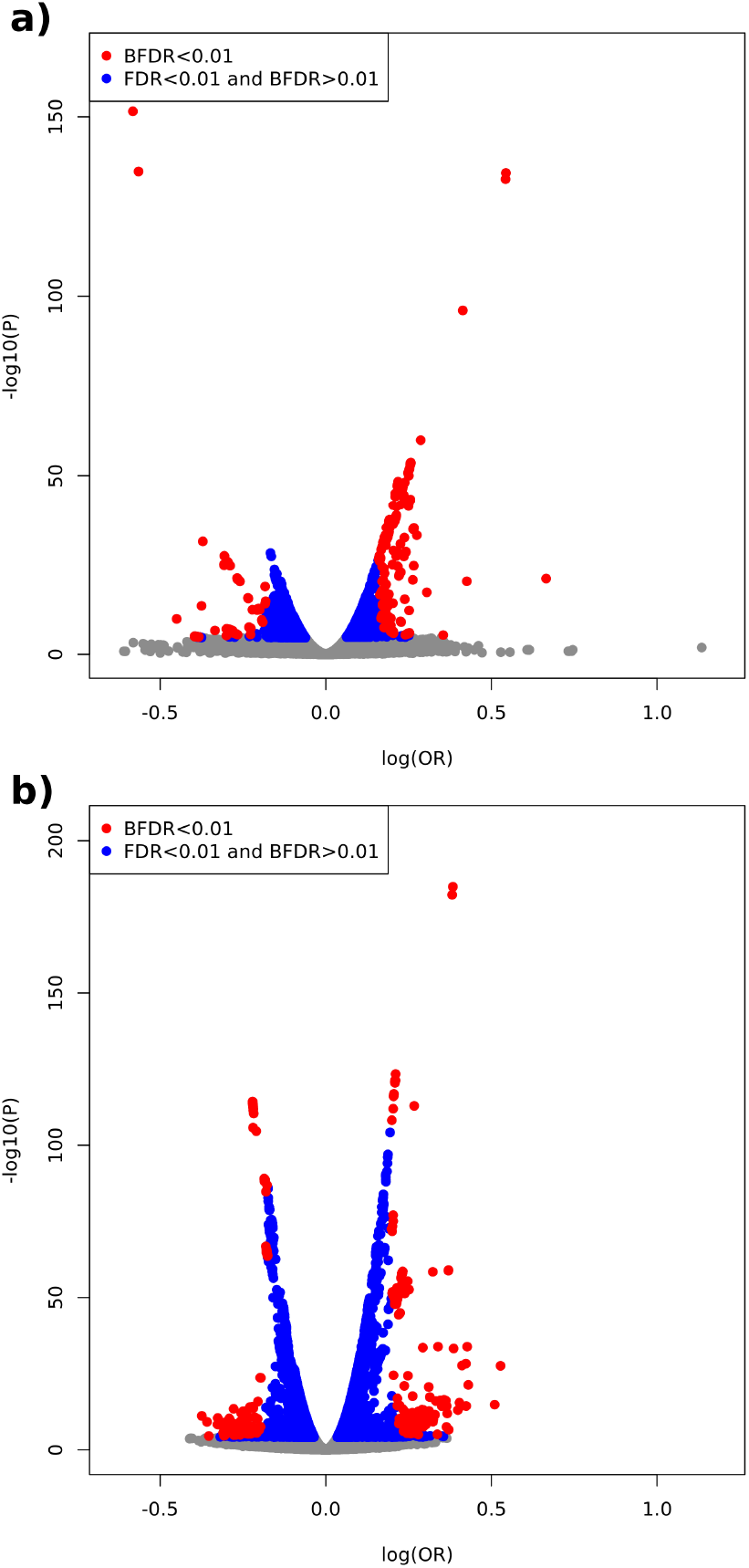
Volcano plots of minor allele effect size (log odds ratio) versus significance (-log10 P values) for a) type 1 diabetes and b) autoimmune thyroid disease. All analysed variants are shown. Variants with either i) BFDR<1% or ii) FDR<1% are highlighted. See Supplementary Figure S3 for the same volcano plots highlighting variants with BFDR<5% or FDR<1%.

**Figure 3.**
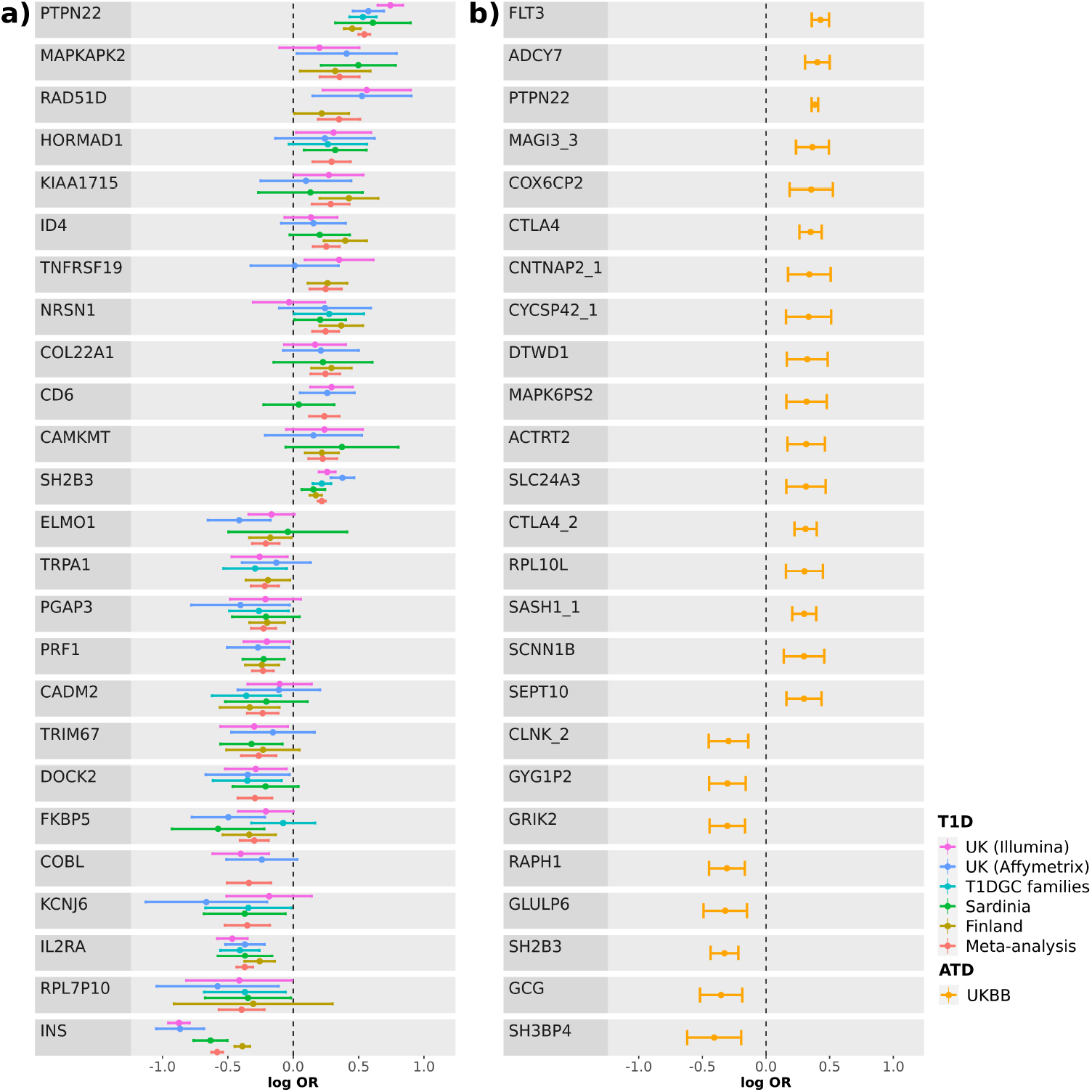
Forest plots of lead variants in 25 largest-effect signals for a) T1D, all five cohorts plus their meta-analysis, and b) ATD. Bars represent 95% confidence intervals.

**Table 1.** Signals with low BFDR, for a) T1D (BFDR<l%) and b) ATD (BFDR<0.5%). See Tables S1-S2 for details of all associated signals. All ORs are for minor alleles. IDs are for each signal’s lead variant. Bolded signals are previously unreported.

| *Nearest gene* | *Band* | *ID* | *MAJ* | *MIN*<br>*(effect)* | *MAF\** | *Imputation*<br>*info\*\** | *P* | *OR (95% CI)* | *FDR* | *BFDR* | *Annotation* |
| --- | --- | --- | --- | --- | --- | --- | --- | --- | --- | --- | --- |
| *PTPN22* | 1p13.2 | rs2476601 | G | A | 9.43% | 99.91% | 4.54E-135 | 1.72 (1.65-1.8) | 9.92E-43 | 8.23E-05 | missense; 3 prime UTR; NMD transcript;<br>non coding transcript exon |
| *INS* | 11p15.5 | rs689 | T | A | 29.94% | 96.24% | 3.02E-152 | 0.56 (0.53-0.58) | 3.06E-27 | 9.88E-05 | NMD transcript; splice region; upstream<br>gene; 5 prime UTR; downstream gene |
| *IL2RA* | 10p15.1 | rs61839660 | C | T | 10.16% | 98.69% | 3.49E-32 | 0.69 (0.65-0.73) | 3.06E-27 | 2.46E-04 | non coding transcript |
| *FKBP5* | 6p21.31 | rs2092427 | G | A | 2.45% | 99.76% | 8.10E-08 | 0.74 (0.67-0.83) | 1.47E-04 | 3.28E-03 | downstream gene |
| *SH2B3* | 12q24.12 | rs3184504 | C | T | 48.66% | 99.96% | 4.62E-49 | 1.24 (1.21-1.28) | 1.47E-42 | 5.07E-03 | missense; downstream gene; NMD<br>transcript |
| *MAPKAPK2* | 1q32.1 | rs75320486 | G | A | 0.97% | 97.14% | 4.31E-06 | 1.43 (1.23-1.66) | 3.95E-03 | 5.24E-03 | non coding transcript; regulatory region |
| ***RPL7P10*** | 1p31.1 | rs111551178 | C | T | 0.77% | 97.06% | 8.36E-06 | 0.67 (0.57-0.8) | 4.45E-03 | 6.14E-03 | intergenic |
| *RPS26* | 12q13.2 | rs705704 | G | A | 33.94% | 98.92% | 1.89E-37 | 1.22 (1.18-1.25) | 1.48E-33 | 6.33E-03 | upstream gene; downstream gene;<br>regulatory region |
| ***ID4*** | 6p22.3 | rs75356149 | G | T | 2.33% | 98.69% | 1.04E-06 | 1.29 (1.16-1.42) | 1.27E-03 | 6.35E-03 | intergenic |
| *PRF1* | 10q22.1 | rs78325861 | C | G | 3.93% | 90.51% | 2.70E-08 | 0.79 (0.73-0.86) | 6.43E-05 | 6.71E-03 | regulatory region |
| ***NRSN1*** | 6p22.3 | rs11965813 | T | A | 1.30% | 99.09% | 1.58E-06 | 1.28 (1.16-1.42) | 1.87E-03 | 6.86E-03 | intergenic |
| *CTRB2* | 16q23.1 | rs55993634 | C | G | 7.88% | 97.07% | 4.76E-14 | 1.2 (1.15-1.26) | 9.00E-10 | 7.22E-03 | downstream gene |
| *PTPN11* | 12q24.13 | rs11066320 | G | A | 42.74% | 99.94% | 1.88E-33 | 1.2 (1.16-1.23) | 6.16E-30 | 7.53E-03 | NMD transcript; regulatory region |
| ***RPL35AP21*** | 9q21.13 | rs17619081 | A | G | 2.98% | 95.54% | 4.01E-07 | 1.23 (1.13-1.33) | 6.33E-04 | 8.00E-03 | non coding transcript |
| *PTPN2* | 18p11.21 | rs7237497 | C | T | 16.49% | 96.88% | 1.17E-18 | 1.19 (1.15-1.24) | 4.48E-14 | 8.06E-03 | intergenic |
| *PGAP3* | 17q12 | rs76096054 | C | A | 1.48% | 99.27% | 2.91E-06 | 0.8 (0.72-0.88) | 2.00E-03 | 9.85E-03 | non coding transcript; NMD transcript |

\*UK Illumina controls \*\*UK Illumina all samples

b)
| *Nearest gene* | *Band* | *ID* | *MAJ* | *MIN*<br>*(effect)* | *MAF\** | *Imputation*<br>*info* | *P* | *OR (95% CI)* | *FDR* | *BFDR* | *Annotation* |
| --- | --- | --- | --- | --- | --- | --- | --- | --- | --- | --- | --- |
| *FLT3* | 13q12.2 | rs76428106 | T | C | 1.29% | 91.97% | 1.39E-34 | 1.53 (1.43-1.64) | 7.16E-31 | 1.26E-04 | NMD transcript |
| *PTPN22* | 1p13.2 | rs2476601 | G | A | 10.25% | 100% | 1.33E-185 | 1.47 (1.43-1.51) | 3.03E-41 | 2.61E-04 | missense; 3 prime UTR; NMD transcript; non coding transcript exon |
| *ADCY7* | 16q12.1 | rs78534766 | C | A | 0.61% | 100% | 3.40E-16 | 1.5 (1.36-1.65) | 2.44E-12 | 3.66E-04 | downstream gene; missense; NMD transcript; non coding transcript exon; upstream gene |
| *CTLA4* | 2q33.2 | rs79877750 | C | T | 0.90% | 88.51% | 4.29E-15 | 1.42 (1.3-1.55) | 1.22E-11 | 7.13E-04 | regulatory region |
| ***MAGI3*** | 1p13.2 | rs547473095 | G | C | 0.57% | 64.56% | 3.19E-08 | 1.44 (1.27-1.64) | 3.32E-05 | 8.89E-04 | non coding transcript |
| *SH2B3* | 12q24.12 | rs117202552 | C | T | 0.86% | 96.54% | 4.76E-09 | 0.72 (0.65-0.8) | 6.43E-06 | 9.43E-04 | upstream gene |
| *CTLA4* | 2q33.2 | rs78847176 | C | T | 0.87% | 89.35% | 2.93E-12 | 1.36 (1.25-1.49) | 8.75E-09 | 1.59E-03 | regulatory region |
| *ATXN2* | 12q24.12 | rs193097290 | A | G | 1.15% | 94.44% | 7.70E-09 | 0.76 (0.69-0.83) | 9.65E-06 | 2.09E-03 | NMD transcript; non coding transcript |
| *SASH1* | 6q24.3 | rs146201355 | A | G | 0.72% | 92.79% | 4.38E-10 | 1.35 (1.23-1.48) | 8.12E-07 | 2.34E-03 | intergenic |
| *PTPN11* | 12q24.13 | rs201479994 | T | TA | 0.95% | 98.10% | 8.71E-08 | 0.76 (0.69-0.84) | 7.81E-05 | 3.00E-03 | intergenic |
| *IFIH1* | 2q24.2 | rs72871627 | A | G | 1.50% | 99.71% | 4.23E-10 | 0.77 (0.71-0.84) | 8.08E-07 | 3.13E-03 | non coding transcript |
| *KRT18P39* | 2q33.2 | rs114937187 | G | A | 0.87% | 90.81% | 7.66E-07 | 0.76 (0.68-0.84) | 4.00E-04 | 3.50E-03 | intergenic |
| *FOXE1* | 9q22.33 | rs925489 | T | C | 33.16% | 100% | 4.56E-115 | 0.8 (0.79-0.82) | 5.04E-41 | 3.72E-03 | intergenic |

\*White Europeans

Of the 219 signals, 103 can be considered ‘new’, being independent (r^2^<0.01) from any lead variants in neither the previously most highly powered GWAS/ImmunoChip studies^13,17,19^ nor in a significantly larger recent ImmunoChip study^18^ (Table S1b). New signals consisted of 40 associations with BFDR<5% (median OR_risk_=1.21, median MAF=4.1%), plus 63 with FDR<1% but BFDR≥5% (median OR_risk_=1.08, median MAF=29.9%), indicating smaller effects on risk. Four of the new associations, *RPL7P10, ID4, NRSN1* and *RPL35AP21* had BFDR<1% (Table 1a). Note that signals are named after the gene closest to the lead variant, although in many cases this will not be the causal gene. These signals have the potential to provide important insights into T1D biology, but due to their low frequencies fine-mapping was unable to pinpoint likely candidate variants, requiring larger samples and more dense genotyping coverage (Figure S5a-d, Table S8). In addition to the 40 new larger effect signals, 49 out of 116 previously reported T1D signals (42%) had BFDR<5% and 12 (10%) had BFDR<1% (*INS, PTPN2, IL2RA, MAPKAPK2* (a.k.a. *FCMR*, a another nearby gene), *FKBP5* (a.k.a. *DEF6*), *PRF1, PGAP3* (a.k.a. *IKZF3*), *SH2B3* and *RPS26* (a.k.a. *IKZF4*), *CTRB2* (a.k.a. *BCAR1*), *PTPN11* and *PTPN2* (OR_risk_ ranging 1.19-1.79), highlighting the importance of their biological effects on T1D risk.

We detected 67 previously reported independent signals that were significant at FDR<1% but had BFDR≥5%, implying smaller effects on risk relative to other true associations despite high confidence that they are disease-associated. Significance was at the genome-wide level (P<5×10^−8^) for 25 of these smaller effect associations with OR_risk_ ranging 1.08-1.12). Genome-wide significance was observed for 58 signals in total (Manhattan and QQ plots in Figures S1a and S2a), with four signals (near *RLIMP2, SLC25A37, MAGI3*, and *LHFPL5*) not previously reported. Numbers of signals satisfying various significance criteria are summarised in Figures 4a and S4a. In total we found eight signals containing multiple conditional signals after stepwise model selection (*INS, PTPN22, IL2RA, CHD9, PTPN2, RLIMP2, PRKCQ* and *AKAP11*, Table S1d).

**Figure 4.**
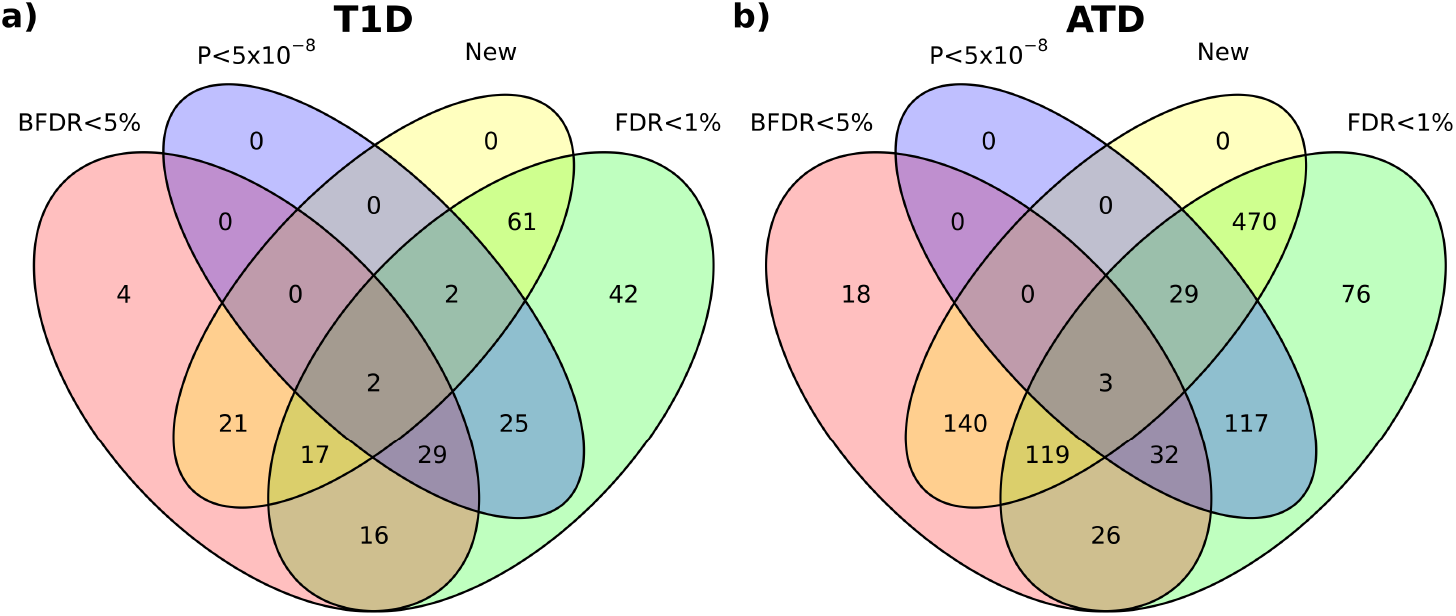
Numbers of new and previously reported signals satisfying BFDR<5%, FDR<1% and genome-wide (P<5×10^−8^) significance criteria for a) type 1 diabetes and b) autoimmune thyroid disease, quantified using the lead variant in each signal. Supplementary Figure S4 shows the equivalent Venn diagrams for results using a 1% BFDR threshold.

Among the new T1D signals, *RAD51D* was notable due to having the third largest observed effect together with a relatively low BFDR, though not passing our 1% threshold (lead variant rs28670687, P=1.74×10^−5^, BFDR=1.36%, OR=1.42 (1.21-1.67), MAF=0.72%), and its lead variant being in LD with three transcribed 3’UTR variants displaying some evidence for causality (fine-mapping log10 Bayes Factors 1.53) (Table S8, Figure S5e), though they could not definitively separated from a larger credible set (log 10 Bayes factor 2.73). The chromosome region encoding its interaction partner, RAD51B, contained a T1D-associated signal with a much smaller effect on risk (OR=0.92 (0.9-0.95), P=6.11×10^−7^, FDR=5.56×10^−4^, BFDR=12.7%). Fine-mapping suggested two separate signals, but we found no obvious causal candidates near *RAD51B* (Figure S5f).

Using the two cohorts with individual genotypes available (UK Illumina and Affymetrix), we re-tested all additive associations for dominant and recessive effects at each lead variant (Table S1c). Of the 219 lead SNPs, four were most significant when the minor allele was dominant (Table 2a), including *RAD51D*. A lower P-value implies a better quality model than additivity due to the number of parameters being the same in both cases. No lead T1D variants were most significant under a recessive model. We also repeated the full GWAS analysis under both dominant and recessive models, finding 16 dominant and three recessive signals that were missed by the additive-model GWAS (independent from additive lead variants at *r*^2^<0.05, Tables S1e-f).

**Table 2.** Signals for which dominant or recessive inheritance models provided the best fits (most significant P value): a) T1D signals and b) Top five largest-effect ATD signals. IDs are for each signal’s lead variant.

| *Nearest gene* | *Band* | *ID* | *MAJ* | *MIN* | *MAF\** | *Imputation info\*\** | *P (additive)* | *OR (additive) (95% CI)* | *Annotation* | *P (best model)* | *OR (best model) (95% CI)* | *Best Model* |
| --- | --- | --- | --- | --- | --- | --- | --- | --- | --- | --- | --- | --- |
| *C11orf21* | 11p15.5 | rs756919 | T | G | 5.10% | 95.71% | 6.21E-06 | 1.13 (1.07-1.19) | intergenic | 3.74E-06 | 2.06 (1.84-2.3) | DOM |
| *RAD51D* | 17q12 | rs28670687 | T | A | 0.72% | 95.28% | 1.74E-05 | 1.42 (1.21-1.67) | NMD transcript; non coding transcript | 1.50E-05 | 1.75 (1.36-2.25) | DOM |
| *CD6* | 11q12.2 | rs117883663 | T | G | 3.53% | 97.59% | 5.20E-05 | 1.27 (1.13-1.42) | downstream gene; NMD transcript; non coding transcript; regulatory region | 2.04E-05 | 1.33 (1.17-1.51) | DOM |
| *TBC1D4* | 13q22.2 | rs554648 | T | G | 41.02% | 97.61% | 1.03E-05 | 1.07 (1.04-1.1) | downstream gene | 6.06E-08 | 1.24 (1.15-1.35) | DOM |

\*UK Illumina controls \*\*UK Illumina all samples

b)
| *Nearest gene* | *Band* | *ID* | *MAJ* | *MIN* | *MAF\** | *Imputation Info* | *P (additive)* | *OR (additive) (95% CI)* | *Annotation* | *P (best model)* | *OR (best model) (95% CI)* | *Best model* |
| --- | --- | --- | --- | --- | --- | --- | --- | --- | --- | --- | --- | --- |
| *TGFB2* | 1q41 | rs767491614 | CAATA<br>AATA | C | 4.75% | 97.30% | 6.25E-07 | 0.9 (0.87-0.94) | intergenic | 1.82E-07 | 0.5 (0.38-0.65) | REC |
| *FLT3* | 13q12.2 | rs76428106 | T | C | 1.29% | 91.97% | 1.39E-34 | 1.53 (1.43-1.64) | NMD transcript | 4.71E-35 | 1.54 (1.44-1.65) | DOM |
| *ADCY7* | 16q12.1 | rs78534766 | C | A | 0.61% | 100% | 3.40E-16 | 1.5 (1.36-1.65) | downstream gene; missense; NMD transcript; non coding transcript exon; upstream gene | 2.75E-16 | 1.5 (1.36-1.65) | DOM |
| *MAGI3* | 1p13.2 | rs547473095 | G | C | 0.57% | 64.56% | 3.19E-08 | 1.44 (1.27-1.64) | non coding transcript | 2.08E-08 | 1.45 (1.27-1.65) | DOM |
| *CNTNAP2* | 7q35 | 7:146077700_<br>TA_T | TA | T | 0.53% | 41.77% | 6.78E-05 | 1.41 (1.19-1.66) | non coding transcript | 4.88E-05 | 1.42 (1.2-1.67) | DOM |

\*White Europeans

Taking an alternate approach to discovery, we performed GWAS meta-analysis on four of the five cohorts (UK Illumina, UK Affymetrix, Sardinians and T1DGC), and replication analysis in the left-out Finnish cohort (FinnGen, 4933 cases and 148,190 controls), providing further evidence that variants with BFDR<5% but FDR≥1% in the discovery data represent replicable statistical associations. Ten signals met these criteria, of which two replicated at P<5%, in line with what would be expected given plausible assumptions about the power of the replication data (P=0.26, binomial test assuming 35% power, see online methods).

### ATD GWAS in UKBB

Using 28,742 unrelated cases and 427,388 unrelated controls, we found a total of 1030 ATD-associated signals (Figures 2b and S3b, Table S2a): 872 variants with FDR<1% and 338 with BFDR<5%, with median OR_risk_ of 1.07 and 1.23, respectively (median MAFs 16.6% and 0.90%). Among the 158 lead variants that failed to pass FDR<1% but passed BFDR<5%, the median OR_risk_ was 1.24 (median MAF 0.75%). We found 34 lead variants with BFDR<1% (median OR_risk_=1.30, median MAF=0.88%) (Table 1b).

Previously established associations in *FLT3, ADCY7, PTPN22* and *CTLA4*^22–24^, among others, have BFDR<1% (Table 1b, Figure 3b), verifying their biological importance. The largest-effect association, *FLT3*, has a lead SNP minor allele (rs76428106 C: OR=1.53, MAF=1.29%) associated with increased monocyte count in UKBB^25^. *FLT3*’s status as the largest known risk effect outside the HLA region has recently been highlighted^26^, and our BFDR estimate of 1.26×10^−4^ confirms its importance. We found nine previously unreported signals including *IL6, FAM117B* and a new signal near *MAGI3*, with BFDR<1% (Tables 1b and S2b, Figure S4b), several of which appeared potentially causal after fine-mapping, while others tagged similarly large-effect loci (Figure S5g-o, Table S9). A number of interesting low frequency signals with large estimated ORs were detected at BFDR<5%, but with FDR≥1%, for example *SH3BP4* (OR=0.67, MAF=0.05%, FDR=2.19%, BFDR=2.80%). Among these, 65/158 (41%) had lead variants that were located in either non-coding transcripts or regulatory regions. Fine-mapping low frequency associations is challenging, but for *SH3BP4* we found suggestive evidence that either the lead variant, which is a 9 bp insertion situated on the boundary of an enhancer element 2.8 Mb upstream of *AGAP1* (log10 Bayes Factor = 2.47), or a second 5 bp insertion in close LD (log10 Bayes Factor = 2.39), is causal (Figure S5p, Table S9).

Despite a considerable overlap in data, by including further samples not of white-European ancestry we detected 32 new signals with genome-wide significance (P<5×10^−8^) that were not found in a recent UKBB ATD GWAS^24^. We found 589 further unreported signals by using an FDR<1% threshold, and 140 further still using BFDR<5% (Figures S3b and 4b, Table S2b). Only three associations from the previous GWAS were further than 500 Kb from one of our lead variants, two of which lay within our definition of the the HLA region and were thus were not part of our GWAS analysis.

Median risk OR for 872 variants with FDR<1% was 1.07 (IQR=0.07), demonstrating the high polygenicity of ATD, with many small effect associations that are detectable with large sample sizes. Genome-wide significance (P<5×10^−8^) was obtained for 181 signals in total (Figures S1b and S2b). Of these, 146 (81%) had BFDR≥5% (risk ORs ranging 1.05-1.13), demonstrating that most genome wide hits are obtained for variants that we are >95% sure are either false associations or, more likely, true associations with true effect sizes below the top 5% (Figure 4b). Only 164/181 (91%) genome-wide significant signals had BFDR≥1%. However, BFDR analysis finds that 76 (28%) of all 269 previously reported associations, including those below genome-wide significance, are likely to have risk effects in the top 5%. Applying stepwise model selection, 35 of the 1030 signals contained multiple conditional signals (Table S2d).

Testing all 1030 lead SNPs under dominant and recessive inheritance models revealed 280 (27%) that were most significantly associated with dominantly acting minor alleles (Table S2c), including 13 BFDR<1% signals such as *FLT3, MAGI3, FABP3P2* and *FAM117B*. Recessive models were most significant for 14 lead SNPs (1%), including *TGFB2* (rs767491614, deletion of AATAAATA), which showed a large protective recessive effect of the minor deletion allele (OR_rec_=0.50 (0.38-0.65), P_rec_=1.82×10^−7^, OR_ADD_=0.90 (0.87-0.94), P_add_=6.25×10^−7^). Details of the five largest-effect non-additive signals are shown in Table 2b. Genome-wide testing of dominant and recessive models detected 39 and 200 signals, respectively, that were independent (*r*^2^<0.05) from any additive lead variants (Tables S2e-f).

### Shared and non-shared genetic causes of ATD/T1D

We define a total of 836 independent signals (173 T1D and 663 ATD) from the full set of 1249 (219 T1D plus 1030 ATD) by removing those with lead SNPs closer than 100 Kb to a more significant lead variant in the other disease, after excluding ATD signals with MAF<3%, as in many cases the most significant SNPs were not present in the T1D data. Colocalisation analysis^27^ of the summary statistics in both diseases found a number of signals showing strong evidence for one of four hypotheses, quantified by the corresponding posterior probabilities (PPs): a) a causal variant exists for T1D only (PP_T1D_), b) a causal variant exists for ATD only (PP_ATD_), c) there are separate causal variants in each disease (PP_separate_) or d) the diseases share a causal variant (PP_shared_). We found 19 signals with PP_T1D_>0.9, and 86 with PP_ATD_>0.9 (Figure 5a-b limited to the 25 with highest PP). Of these, only 3/19 PP_T1D_>0.9 signals and 24/86 PP_ATD_>0.9 signals were found to associate with other immune diseases in the Open Targets Genetics Portal^1^ (Table S10). Strong evidence for separate causal variants was found in 16 signals (PP_separate_>0.9, Figure 5c) and shared causal variants in 31 signals (PP_shared_>0.9, Figure 5d, Tables S5 and S6 for full results). Several established autoimmune loci show high evidence of shared causal variants, e.g. *PTPN22, CTLA4, UBASH3A* and *BACH2*. The colocalisation method assumes a single causal variant per disease within each signal, so SNPs in LD with multiple signals’ lead variants (*r*^2^>0.01) were assigned to the most significant signal and excluded from others.

**Figure 5.**
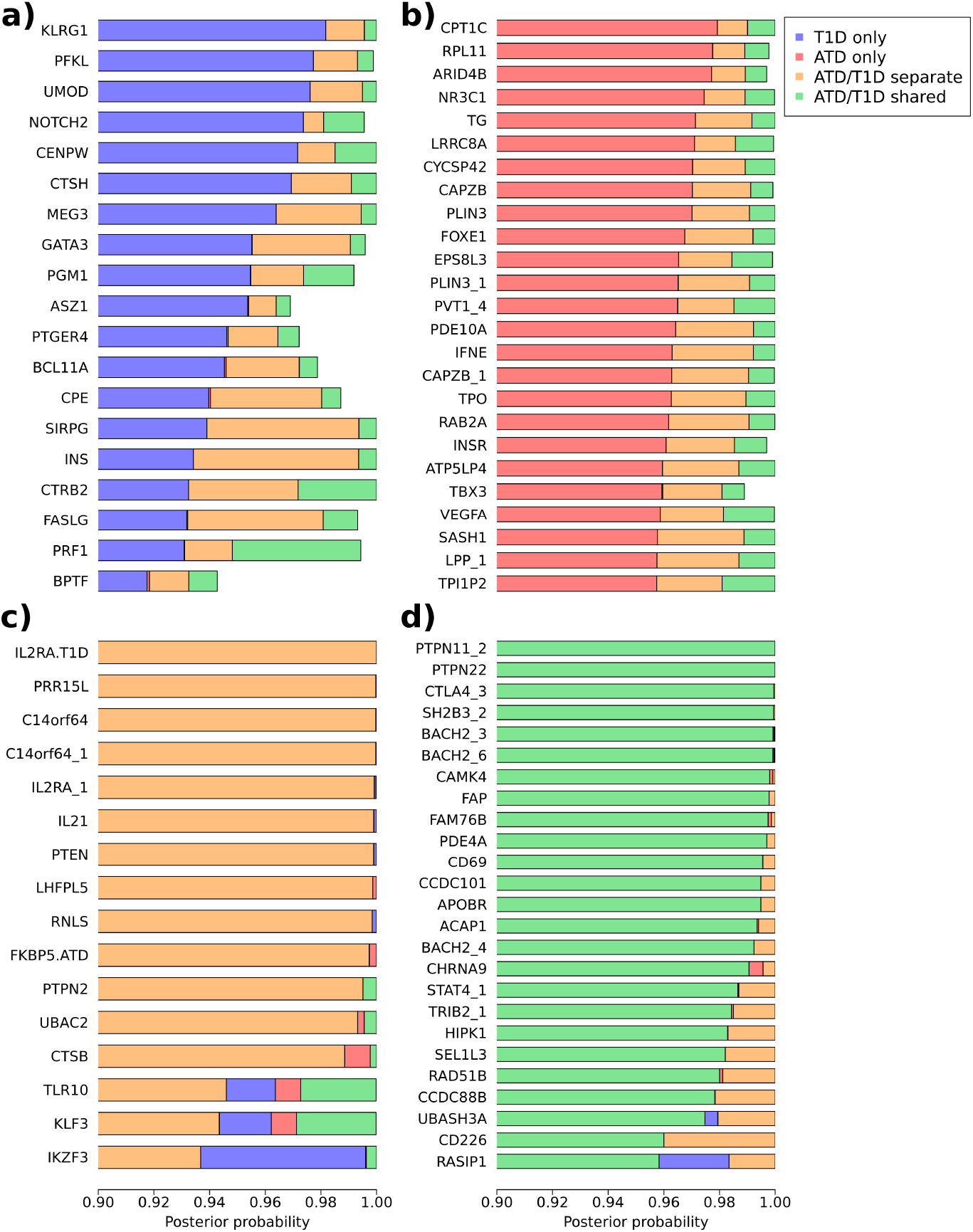
Posterior probabilities of colocalisation for signals with a) PP_T1D_>0.9, b) PP_ATD_>0.9, c) PP_separate_>0.9 d) PP_shared_>0.9. Up to the top 25 signals, when ranked by maximum posterior probability, are shown in each case (full results in Tables S5 and S6). Multiple independent signals near the same gene have numerical suffixes separated by an underscore. Posterior probabilities are truncated at 0.9 to aid visualisation.

When both diseases share a causal variant, it is possible that the risk allele for one disease is protective towards the other. However, we found that risk and protective alleles for the lead variants (or largest available association present in both the ATD and T1D datasets) were the same in both diseases for all 31 signals with PP_shared_>0.9.

To determine the extent of shared genetic architecture between T1D and ATD, we examined the proportions of signals with high evidence for causal effects on only one of the diseases (PP_T1D_ or PP_ATD_>0.9). Only 86/663 (13%) ATD signals showed some degree of ATD-only effect by this definition (PP_ATD_>0.9), and 19/173 (11%) for T1D (PP_T1D_>0.9). However, many signals have quite high PP for the null hypothesis of neither variant being associated (PP_null_), with many more ATD signals showing PP_null_>0.33 proportionally (N_ATD_=357/663, N_T1D_=66/173). This is most likely due to the conservative prior probability (99.99%) assigned to the null hypothesis that a signal is not disease-associated. As local FDRs (fdrs) are empirical Bayesian estimates of the probability of zero effect for a given P value^28^, we computed rescaled posterior probabilities (rsPPs) in order to maintain approximate consistency with our GWAS fdrs. Taking ATD as an example, PP_ATD_,PP_shared_ and PP_separate_ can be scaled to sum to 1-fdr_ATD_, and PP_null_ and PP_T1D_ scaled to sum to fdr_ATD_ (see online methods). After rescaling, 80/173 T1D signals (46%) and 357/663 ATD signals (54%) showed rsPP_T1D/ATD_ > 0.9 (full results in Table S7), indicating that many regions may have signals specific to one of the diseases.

Evidence for the four novel T1D signals with BFDR<1% having effects on T1D only was limited before rescaling (PP_T1D_ ranging 0.73-0.85) but stronger after (rsPPτiD ranging 0.91-0.94), suggesting that these signals are not present in ATD (Figure S6a-d). Of the four, only *ID4* associated with another autoimmune disease in the Open Targets Genetics Portal (ulcerative colitis). Despite weak evidence for T1D specificity using unscaled colocalisation PPs (PP_T1D_=0.15, PP_null_=0.83), *RAD51D* had signals that appeared to be different between ATD and T1D (Figure S6e), and fdr-rescaling gave rsPP_T1D_=0.92, while *RAD51B* strongly colocalised (PP_shared_=0.98, Figure S6f). Only five ATD signals with MAF≥3% had BFDR<1%. Two had PP_ATD_>0.9 (*VAV3* and *FOXE1*, Figure S6g-h) and three had PP_shared_>0.9 (*PTPN22, SH2B3* and *CTLA4*, Figure S6i-k). Consistent with having ATD-specific causal effects, *VAV3* and *FOXE1* did not associate with other autoimmune diseases in the Open Targets Genetics Portal, while *PTPN22, SH2B3* and *CTLA4* had several other associations consistent with general autoimmune roles (Table S10).

Colocalisation analysis suggested that power might be increased for discovering T1D associations with pleiotropic ATD effects by utilising information from the larger ATD study. There are a number of methods for jointly analysing pleiotropic phenotypes^29–32^, but we took a straightforward approach by calculating BFDRs and FDRs for T1D association using only those SNPs satisfying our significance criteria in ATD (BFDR<5% or FDR<1%), excluding the HLA region. These SNPs were enriched for T1D association, so the resulting BFDRs and FDRs were lower. Of the 45,633 SNPs satisfying the significance criteria in ATD, 34,381 were present in the T1D data. An additional 105 T1D signals were identified (r^2^<0.01 with any lead variant in the primary T1D analysis), 76 of which have not been previously reported as T1D-associated (Tables S3a and S3b). Most had small effect estimates (median OR_risk_ 1.06, IQR=0.03), but one signal (*CD200R1*) had OR_risk_=1.20 (FDR=7.97×10^−3^).

## Discussion

Using highly powered GWAS analyses of T1D and ATD, we detected a large number of independent genetic signals, many of which have not been associated with either disease before. It is has become widely acknowledged that many, even most, variants are likely to have non-zero effects on complex phenotypes^33–35^. A large proportion of small-effect ‘polygenic’ risk variants are likely to be discovered in a large GWAS, especially applying thresholds on FDRs rather than conservative Bonferroni-corrected P values, exemplified by our discovery of 691 ATD signals below genome-wide significance but with FDR<1%, mostly with small effects. Increasing availability of even larger genome-wide SNP and sequencing datasets is likely to exacerbate this issue. Introducing the concept of the BFDR, and developing a method for its estimation, we were able to narrow down FDR<1% signals to those we were confident had larger than average risk effects, by selecting SNPs with BFDR<1%. We found only 34 signals with BFDR<1% for ATD, and 16 for T1D, most of which have been previously reported, indicating that although current GWAS datasets are well-powered to reject the null hypothesis for many SNPs, there is less power to differentiate between variants’ effect sizes. Signals with large ORs that have been missed by previous GWAS studies are expected to have lower MAFs, as we found for 13 previously unreported BFDR<1% signals (four in T1D, nine in ATD). Fine-mapping low frequency signals is challenging and in most cases evidence regarding which specific variants are causal is lacking.

A more lenient BFDR<5% threshold allowed us to detect signals that did not meet FDR<1% (25 for T1D and 158 for ATD) but were nevertheless likely to have larger effects, at the cost of an increased error rate (with a variants’ FDR always being lower than its BFDR). Of the 239 genome-wide significant signals in T1D and ATD, only 66 (28%) had BFDR<5%, showing that high significance may be accompanied by evidence for a stand-out effect only in a minority of cases.

One example of a more lenient BFDR association lies near *RAD51D* (BFDR=1.36%, FDR=1.08%, OR=1.42), which encodes a DNA repair protein belonging to the *BRCA1* and *BRCA2* interacting BCDX2 complex^36^. Our finding that *RAD51B,* another BCDX2 component, is T1D- and ATD-associated, albeit with much smaller effects on risk, further supports this association. Three potential causal variants are in the *RAD51D* 3’UTR, suggested by fine-mapping and in high LD with the lead variant (rs28670687). Mutations in *RAD51D* are associated with ovarian cancer^37^, and cells lacking *RAD51D* frequently undergo deletions of large chromosome segments, demonstrating its role in genome stability^38^. Pancreatic beta cells are long lived and non-duplicating, thus requiring high levels of genome-stability for normal functioning, and could therefore be where *RAD51D* variants lead to T1D risk. Colocalisation analysis showed no effect of *RAD51D* on ATD.

We found that 25 previously reported ATD signals have significantly larger risk effects than the polygenic risk background due to having BFDR<1%, including well known signals near *ADCY7, PTPN22, CTLA4, SH2B3, IFIH1* and the recently reported *FLT3^26^*, which appears to act dominantly (OR_dom_=1.54). We also found nine new independent signals with BFDR<1%, including those near *MAGI3, IL6* and *FAM117B*. However, we found that 81% of genome-wide significant ATD associations had risk effects that could not be said to be larger than a randomly chosen ATD effect (BFDR≥5%), casting doubt over the extent of biological insight provided by thresholding on genome-wide significance alone. High statistical significance in these cases is a consequence of the risk variants being very common in the population, rather than by them having stand-out effect sizes.

We found many further signals using the more lenient BFDR<5% threshold, including a low frequency insertion situated 32bp from an enhancer element near *SH3BP4*, showing a large protective effect on ATD (OR=0.67). The enhancer element lies in a region of few genes, but is downstream of *SH3BP4* and upstream of *AGAP1*. Despite having low MAF in most populations, the protective minor allele is found in roughly 25% of West-African individuals, emphasising the requirement for more diverse GWAS datasets. Signals passing BFDR<5% but not FDR<1% have low frequencies (below 5%), and are mostly imputed, thus should be considered tentative findings in need of further follow up.

The BFDR is designed for selecting genetic associations for follow-up in the ‘post-GWAS’ era, in which the prior expectation is for there to be a very large number of small-risk associations, many with unknown biological connections to the phenotype under study. A cruder approach would be to follow-up variants that are both significant at a satisfactory FDR (e.g. <1%) and have a satisfactory observed effect estimate. However, thresholds on observed effect sizes are fairly arbitrary when there is no consideration of how much these are affected by noise (sampling error), and what the distribution of true effects is. Our main statistical innovation for estimating the BFDR is the ‘prior splitting’ method, which allows the distribution of pairwise SNP effect estimate differences to modelled as a mixture of negative, null (zero) and positive differences, without explicit modelling of the prior distribution of true effect differences. By constraining the relationship of the negative and positive mixtures to the null distribution, one can guarantee that the negative and positive mixtures contain only negative and positive true effect differences respectively (see SI appendix).

Many genetic risk variants are likely to have pleiotropic effects on T1D and ATD, as both are autoimmune disorders. As our ATD analysis in UKBB was highly powered, this provided an opportunity to leverage additional statistical power for T1D association analysis. Restricting computation of T1D FDRs and BFDRs to SNPs associated with ATD, we found 76 associations that were not associated in our primary T1D GWAS nor in previous studies, though these had mostly small risk effects.

Among our GWAS signals, 19 signals had causal effects on T1D but not ATD, of which 16 did not associate with 12 additional autoimmune diseases, suggesting causal mechanisms outside of the immune system, e.g. in pancreatic islets. Similarly, 86 signals had causal effects on ATD but not T1D, of which 62 had no other autoimmune associations, suggesting causal roles unrelated to autoimmunity. Integrating FDR and colocalisation data, we found that 46% of T1D signals and 54% of ATD signals show some evidence for having causal effects on one disease but not the other. All four previously unreported BFDR<1% T1D signals showed evidence for no-effect on T1D, only one of which associated with other autoimmune diseases (ulcerative colitis). Of the five BFDR<1% ATD associations available for colocalisation, two had ATD-specific causal variants and three had shared causal effects with T1D plus additional autoimmune associations. Altogether, these results suggest that new or established large effect GWAS associations are roughly as likely to have disease-specific effects as to have general effects predisposing to several diseases, e.g. via autoimmunity.

By using a conventional FDR<1% threshold in addition to BFDR<5%, we find further common (high MAF) associations for both diseases, but with smaller risk effects. The median effect size of lead SNPs with FDR<1%, i.e. those selected using significance alone, was 1.10 for T1D and 1.07 for ATD (median MAF 26.7% and 16.6%). However, this missed 25 T1D and 158 ATD variants passing BFDR<5%, with median effect sizes of 1.23 and 1.24 (median MAF 2.93% and 0.75%). Associations with low FDR but small effects may be of limited importance for exploring biological hypotheses, and we recommend using a combination of effect size and significance i.e. using the BFDR. Nevertheless, common variants with smaller effects may be useful for gene-category enrichment analysis, fitting polygenic risk scores, Mendelian Randomisation, or for providing supporting evidence for the involvement of any larger effect variants with which they biologically interact.

## Online Methods

### BFDR simulation experiments

For 100 genetic architectures, we simulated a mixture of 19 normal distributions with mean zero and SDs randomly drawn from an exponential distribution, to which we added a point-null distribution as the 20th mixture component. Exponential rate parameters were chosen separately for each architecture by drawing 100 random uniform variables between 0.1 and 1, allowing variability in signal/noise ratios between architectures. Mixing proportions were simulated from a Dirichlet distribution parameterised by the vector 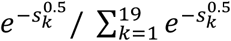 divided by its median value, where *k* identifies a mixture distribution and *s_k_* is its SD, such that smaller mixing proportions were favoured for distributions with large SDs, consistent with realistic genetic architectures. The 19 resulting mixture proportions were standardised to sum to 0.5, and we included the 20th mixture proportion of 0.5 corresponding to the null mixture. We sampled 100,000 multinomial variables from the vector of 20 mixture proportions, providing the mixture memberships for each SNP, and for each SNP then drew a random variable from the corresponding normal distribution, representing its true effect size on the Z-sore scale, *θ*. Allele frequencies for each SNP were simulated from a beta distribution with shape parameters equal to 0.8, after which they were scaled to lie between 0.5% and 99.5% (reflecting how SNPs with MAF<0.05% are typically not analysed in GWAS) and approximate standard errors (SEs) for the log OR estimates calculated as:

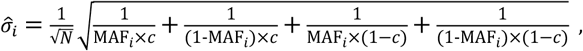

where *N* is the sample size, *c* the proportion of cases, which we set to 15,000 and 0.5 respectively, and *i* a SNP identifier. Rather than simply taking true effect sizes (logORs) to be 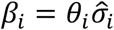, following the definition of *θ_i_* as a Z-score, we allowed for the relationship between MAF and effect size to differ between genetic architectures by setting:

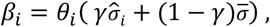

where *γ* was sampled separately for each genetic architecture from a uniform distribution bounded by 0 and 1, and 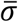 was the mean SE across all SNPs within the simulation. High *γ* therefore induces a strong positive relationship between effect size and SE, as larger SEs amplify *θ_i_* further from zero. Estimated logORs were then simulated as 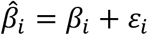 where *ε_i_* was a normal random variable with mean zero and SD 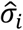. BFDR estimation, returning tail-area estimates of BFDRs, FDRs and BDRs (see SI appendix) for each SNP, was run using priorsplitteR on the 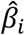 and 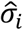 data (for all SNPs *i* in 1,2…100,000) within each simulation. Outlier variants for exclusion were defined as those with z-scores greater than 12 (default 15) and Expectation Maximisation convergence thresholds were relaxed from 10^−6^ to 10^−4^ to save computation time, but otherwise we used the default settings given in the SI appendix.

After BFDR estimation, SNPs were thresholded according to BFDR, BFDR and FDR tail-area probabilities, using a thresholds of 0.001, 0.01, 0.025, 0.05, 0.1, 0.2, 0.3 … 0.9 and 1. At each value of α, we recorded the true error for each SNP *i* with estimated BFDR ≤ α, to quantify how well this error was controlled by the estimation method, based on true FDR and BDR errors:

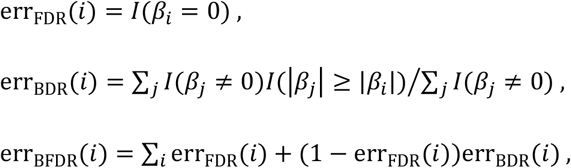

where *I*(·) is an indicator function taking the value one if the condition in parentheses is met or zero otherwise. The true FDR error is therefore one if the SNP is null and zero if it is non-null, whereas the true BDR error is the proportion of non-null SNPs with true effect sizes at least as large as SNP *i*. The true BFDR error is an empirical version of Equation 1 using the known true effects from the simulation. Averages of each of the three errors were taken over *i* for SNPs passing the different a thresholds, within each architecture (Figure 1).

### T1DGC data QC and GWAS analysis

Samples were obtained through the Type 1 Diabetes Genetics Consortium (T1DGC). T1DGC samples were genotyped on Illumina Human Core Exome beadchip following manufacturer protocols and genotype clusters were generated using the Illumina GeneTrain2 algorithm at University of Virginia. Since multiple array versions were used, we harmonized variants across array versions in the following way: for those SNPs that are available on the 1000 Genomes Project SNP panel, we align them to the 1000 Genomes Project SNPs separately for each array version; for those SNPs that are not available on the 1000 Genomes Project SNP panel, we harmonize them according to their positions/names, allele labels and allele frequencies that are specific to each of the four array versions.

Sample identity was confirmed by comparing genotypes from the same samples generated with an alternative array^18^ (ImmunoChip). Variants were removed for the following reasons: a) more than 5% of genotypes were missing, b) genotypes were inconsistent across duplicates (discordant in >1% of duplicate samples or MZ twins), c) genotype frequencies deviated from Hardy Weinberg Equilibrium (P < 10^−6^), d) Mendelian inconsistencies in more than 1% of trios or parent-offspring pairs or e) more than 10% of homozygous parent-offspring pairs or trios with heterozygous offspring. Samples were removed if more than 5% of genotypes were missing or genotypes were inconsistent with reported sex. Sample pedigree information was confirmed or corrected using genotype-inferred relationships, as determined using the software KING^39^. After QC, there were 3173 affected-offspring trios.

Subjects with European ancestry were identified for analysis using KING. Specifically, genetic principal components (PCs) were generated for 1000 Genomes Phase 3 subjects and T1DGC subjects were projected onto this PC space. Then a Support Vector Machine was used to classify T1DGC subjects into one of five ancestral super-populations, as described here https://www.kingrelatedness.com/manual.shtml.

European individuals (N=10,406) were aligned to the Haplotype Reference Consortium (HRC) reference panel using available tools (https://www.well.ox.ac.uk/~wrayner/tools/index.html#Checking) and imputed to the HRC using the Michigan Imputation Server. Imputed variants were filtered for imputation quality (removed variants with imputation R-squared <0.3) and Mendelian errors (removed variants with errors in > 1% of homozygous parent-offspring pairs or trios with heterozygous offspring). ORs were derived as OR=T/U, where T and U are the numbers of transmitted and non-transmitted alleles, and SEs of log ORs for inverse-variance weighting were obtained following Kazeem and Farrall^40^. To prevent extreme ORs (e.g. zero or infinity), we added 0.5 to both T and U for a given SNP whenever either value was 5 or lower.

### UK T1D cohorts QC and GWAS analysis

Type 1 diabetes summary statistics were generated using GWAS data from 13,245 UK individuals in two cohorts: 7977 genotyped using the Illumina Infinium 550K platform (3,983 cases and 3,994 controls) and 5,268 using the Affymetrix GeneChip 500K platform (1926 cases and 3,342 controls), analysed in previous publications^41,42^. We refer to these collections as ‘UK Illumina’ and ‘UK Affymetrix’. Genotypes were imputed using the haplotype reference consortium (HRC) haplotypes and the Michigan Imputation server, pre-phasing using SHAPEIT2 and imputing using Minimac3^43^. Variants failing either of two imputation quality criteria in either UK cohort were removed: a) imputation information score of <60% in either cases or controls, or b) difference in imputation information score between cases and controls > 1% together with MAF < 5%. Exceptions were made for two well-established T1D variants in the *INS-IGF2* region (rs689 and rs3842753), which were poorly imputed in the Affymetrix cohort but well imputed in the Illumina cohort. GWAS summary statistics were produced using the ‘newml’ method from SNPTEST, including the three largest PC covariates. Variants were LD pruned (*r*^2^<0.3) and low (<1%) MAF SNPs removed during calculation of PCs. PCs were calculated within UK Affymetrix and Illumina collections separately.

### GWAS meta-analysis of five type 1 diabetes cohorts

Summary statistics for the two UK cohorts and T1DGC were meta-analysed, together with a Sardinian cohort^42^ (1,558 cases and 2,882 controls, genotyped on Affymetrix 6.0 and Illumina Omni), imputed from a custom reference panel of 3,514 Sardinians, and samples from the FinnGen biobank resource (data freeze 4, phenotype code E4_DM1, n=4933 cases and 148,190 controls). Finngen test statistics correlated well with the test statistics obtained using strictly defined T1D cases (Figure S7). Variants with MAF<0.5% in either UK cohort or in the Sardinian cohort were removed. SNPs not present in at least one of the UK cohorts were removed, as statistical power was likely to be low, and it is possible BFDR is unreliable for effect estimates with low precision. As the HLA region is already well established as being T1D-associated, and has extensive LD which may interfere with downstream analysis, we removed this as standard (40,656 SNPs with build 37 positions 25-35 Mb on chromosome 6), leaving 6,254,180 SNPs.

Combined estimates of effect size from the five cohorts were obtained using inverse-variance weighting, in R:

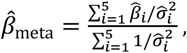

where 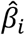 is the estimate of the log OR for the *i*th cohort and 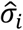 its estimated SE, which are set to zero and infinity respectively when the SNP is missing in cohort *i*. P values were computed from the meta-analysis Chi-square statistics 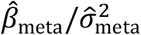 (1 degree of freedom), where 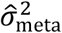 is the variance of 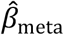 :

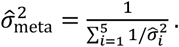

Meta-analysis of dominant and recessive models was restricted to available SNPTEST results from the two UK cohorts, using the same SNPs as in the additive meta-analysis. Recessive association results for low frequency SNPs had deflated significance relative to the null distribution, which creates problems for the distribution-fitting steps of BFDR estimation. We therefore excluded SNPs with MAF < 5% (in either UK cohort, combined cases and controls) from these steps in the T1D recessive BFDR analysis, but reintroduced them for estimating individual SNP BFDRs based on the fitted distributions. As the additive results derive from the larger five-cohort meta-analysis, the number of signals we determined to be dominant or recessive by comparing the P values of the three inheritance models is likely to be conservative.

We saw an inflation of meta-analysed Chi-square statistics (ratio of the observed median statistic over the null median of 0.456, *λ*_GC_ = 1.12, quantile-quantile plot in Figure S2a). Analysis of the T1DGC cohort, using TDT, is immune to population stratification. Inflation was comparable within this cohort alone (*λ_GC_* = 1.09), consistent with polygenicity of T1D, rather than population stratification, being the major cause of test statistic inflation observed in the meta-analysis. Using LD-Score regression gave an estimate of 0.99 for the inflation factor due to population stratification, unbiased by the effects of polygenic association signals^44^.

To assess the replicability of variants, discovery meta-analysis was performed without the inclusion of the FinnGen cohort, and replication of lead variants satisfying BFDR<5%, or FDR<1% significance criteria was performed within FinnGen. Definition of signals was performed as for the full five-cohort meta-analysis. When lead variants were not available in the replication sample, the closest LD proxy was selected using white-European LD data from UKBB. We found 149 signals in the discovery phase, of which 63 passed P<5% in the replication phase (Table S4). Replication power at P<5% was 35% for either common variants with OR=1.05 (MAF=20%) or low frequency variants with OR=1.2 (MAF=1%)^45^, assuming *D’*=0.8 and 1% T1D prevalence in Finland. We found 10 variants satisfying the lenient BFDR<5% threshold with FDR≥1%, in the discovery phase, but as expected their MAFs were mostly low (median 4.2% in FinnGen, IQR=4.8%), implying low power for replication. We found that 2/10 of these variants had P<5% in the FinnGen replication set, which does not depart significantly from a model in which each SNP has 35% power, consistent with them having true effect sizes and MAFs similar to those described above, as a one-tailed binomial test with probability 0.35 gave P=0.26.

### Autoimmune thyroid disease GWAS in UKBB

Imputed post-QC genotype data was available for 487,409 individuals and 93,095,623 autosomal variants from the UKBB. Phenotype data was available for 487,320 individuals for whom genotype was also available, and used to designate 29,045 individuals (5.96%) as ATD cases. The majority of these possessed hypothyroidism/myxedema as a non-cancer illness code (n=24,403), and 4642 additional cases were found using ‘other hypothyroidism’ (as distinguished from ‘subclinical iodine-deficiency hypothyroidism’) ICD10 main (n=67) and secondary (n=4575) codes. The various forms of hyperthyroidism were excluded from our definition of ATD as it is likely that many genetic variants have heterogeneous, possibly opposite, effects to those on hypothyroidism.

We restricted analysis to 77,675,727 autosomal SNPs with imputation information scores > 0.3. Although the UKBB genotype data has already been subjected to QC^46^, we removed 153,773 SNPs showing significant departure from Hardy-Weinberg Equilibrium at P<10^−12^, as these probably result from poor genotype calling, and were not previously removed by the original within-batch filtering approach^47^.

We detected first and second degree relatives via IBD sharing > 25% using PLINK 2’s King-robust estimator^39,48,49^ and removed one random individual (n=31,190) from each pair, leaving 456,130 (28,742 cases and 427,388 controls). Relatedness was calculated using 12,788 independent autosomal variants with MAF>5% (pairwise *r*^2^<0.01, pruned using PLINK^48^).

Logistic regression models for additive risk of allele dosage on ATD were fitted for all remaining 11,261,140 autosomal SNPs outside of the HLA region with MAF>0.5%, using PLINK2. Firth regression was used instead of logistic regression for SNPs with empty contingency table cells, or when logistic regression otherwise failed to converge, as implemented in PLINK2’s firth-fallback option. We controlled for population stratification using the 20 largest genetic PCs, available from UKBB, as covariates. These have been shown to reflect the broad range of ethnic backgrounds from which the participants are drawn^46^, though the majority of participants are white Europeans. We also controlled for age (at initial assessment) and genotypic sex.

Chi-square statistics were inflated with *λ_GC_* = 1.19 (Figure S2b), but this did not differ when controlling for the 40 largest PCs when we tested on chromosome 22 alone. The analysis was repeated for 381,380 white European individuals (24,332 cases and 357,048 controls), identified by UKBB using a combination of self identified ethnicity and genetic ancestry PCs, giving *λ_GC_* = 1.18, suggesting that inflation was primarily due to polygenic association signal rather than population stratification. LD-score regression suggested minor inflation due to population stratification, with the intercept of 1.06 implying that a large majority of inflation is due to polygenic signals of association, and this was similar (1.05) when using GWAS results derived from Europeans only.

Analysis of dominant and recessive effects was performed with the same covariates. To overcome computational limitations, we used PLINK2’s firth-residualize approximation to firth regression^50^ for analysing dominant and recessive effects across all SNPs. As for T1D, we found that recessive associations for low-MAF SNPs showed some deflation of significance compared to the null distribution, though less extreme than we observed for T1D, so SNPs with MAF<2% were withheld from recessive BFDR model-fitting.

### Definition of signals

BFDR analysis was run on each GWAS analysis separately (including separately under additive, dominant and recessive models) under the default settings, using non-HLA variants, providing tail-area estimates of both FDR and BFDR (see SI appendix for details on BFDR estimation). Although it would be possible to use the popular Benjamini-Hochberg procedure^51^ to calculate FDRs, we use our own method for both FDR and BDR components of BFDR in order to maintain a consistent SNP effects model. Our criteria for a variants’ significance were FDR<1% or BFDR<5%, though we were also particularly interested in the subset of signals with BFDR<1% (see main text). To define independent signals among the significant variants, we selected the most significant SNP (lowest P value) on each chromosome, designating this as the lead SNP for the signal, before removing any significant variant in LD (*r*^2^>0.01, up to maximum 1 MB distance), then choosing the next most significant SNP remaining as the lead variant for the next signal. This process (‘LD clumping’) was repeated until no SNPs passing our significance criteria remained. Due to the ubiquity of LD, single-SNP associations, i.e. those that have no disease-associated LD partners, are likely to be spurious due to problems with genotyping or imputation, especially when MAF is low. At each step we therefore excluded the most significant SNP from the process if it had no LD partners (*r*^2^>0.1) with log10 P values lower than log10(P)/3-1, where *P* is the lead SNP P value. LD calculations were performed in plink^52^ using 381,380 individuals from the UKBB, after restricting to white-Europeans and removing first and second-degree relatives (using UKBB data field 22006). LD comparisons were restricted to a sliding window of 1 MB. The signal definition procedure was performed for each set of GWAS results separately (e.g. ATD additive, T1D additive, ATD dominant, T1D additive within ATD signals, etc.). We designated T1D signals as ‘new’ if they had *r*^2^<0.05 (using the UKBB LD data) with, and were physically located at least 250kb from, the lead variants from any established regions^13,17–19^. We used the same criteria to identify new signals for ATD, establishing lead variants’ independence from 175 found by Kichaev et al. using integration of functional enrichment information and GWAS data from UKBB hypothyroidism cases (white Europeans only), plus additional data sources^24^.

### Annotation of lead variants

Functional annotations for each lead variant were obtained using the biomaRt R package (Ensembl build 38 human SNP database). We also wrote command-line GraphQL and R scripts to automatically download and filter lists of immune disease pheWAS associations (P<5×10^−5^) from the Open Targets Genetics Portal^1^ (https://genetics.opentargets.org), for each lead variant. The diseases we filtered for were Addison’s disease, asthma, celiac disease, Crohn’s disease, eczema, hayfever, lupus, multiple sclerosis, psoriasis, rheumatoid arthritis, ulcerative colitis and vitiligo.

### Colocalisation of genetic signals between type 1 diabetes and autoimmune thyroiditis

Colocalisation between ATD and T1D GWAS signals was performed for 500 Kb physical windows around each lead variant using the coloc package^27^, after excluding variants not present in both ATD and T1D datasets. ATD signals with MAF<3%were not included, as a large proportion of their most significant variants were not present in the T1D data. Variants having *r*^2^>0.01 with a lead variant from a more significant (lower P value) signal were removed to ensure analyses pertained to the signals in question, and single-SNP associations were removed as previously described. LD data for coloc plots, and MAFs for coloc analysis, were computed using UKBB (white Europeans only). When lead variants in T1D- and ATD-discovered signals were within 100 Kb of each other, the signal for the least significant disease (largest P value) was removed to avoid repeating analysis of the same signals. For comparing effect directions between diseases, we used the lead variant when this was present in both T1D and ATD datasets. Otherwise, effects were compared at the variant with the lowest P value in either disease, among all SNPs in the physical region that were present in both datasets.

Rescaling of posterior probabilities to be consistent with FDRs was performed using the local FDR^28^ (fdr) of each signal’s lead variant, for the disease that it was originally associated with at the GWAS stage. Then, taking a T1D signal as an example, the rescaled posterior probabilities (rsPPs) are:

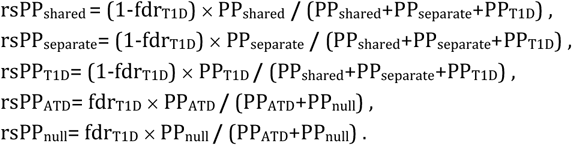

This makes use of the fact that the fdr can be interpreted as an estimate of the posterior probability of no association for a given P value, and so 1-fdr is the equivalent estimate of probability of association (i.e. either a shared causal variant, two separate causal variants or a causal variant only for the disease in question). Local FDRs were computed as an essential part of BFDR analysis (see SI appendix).

### Stepwise regression and Fine-mapping

Stepwise model selection was performed on GWAS summary statistics and LD data from UKBB white Europeans (n=381,380), before joint regression analysis of the selected SNPs, using COJO^53^. SNPs were incorporated into the model if they had a stepwise P value either a) lower than genome-wide significance (P<5×10^−8^) for signals where the lead variant was genome-wide significant or b) lower than the GWASP value for the lead variant for signals where this was greater than 5×10^−8^. Signals were defined as SNPs within 250 Kb of each lead variant, but not in LD with any more significant lead variants (r^2^>0.01). Variants previously determined to be single-SNP associations were omitted, as described above. The meta-analysis sample size, for each variant, was calculated as the summed sample sizes of the four case-control cohorts plus half the number of informative allele transmissions in the T1DGC TDT test. Fine-mapping for ATD and T1D was performed using FINEMAP version 1.4^54^, using the GWAS summary statistics and LD data from UKBB (381,380 white Europeans). For T1D, fine-mapping was restricted to the largest non-Finnish meta-analysis cohort only (UK Illumina 550K, 3983 cases and 3994 controls), to ensure homogeneity of genotyping coverage. Fine-mapping for each signal was performed using all SNPs within 500 Kb windows around the lead SNP. Due to limited statistical power, fine-mapping was not performed on the additional T1D signals detected via pleiotropic effects on ATD, which have mostly small effects on T1D risk.

## Supporting information

Supplementary Note: T1DGC Membership

## Data availability

Type 1 diabetes GWAS meta-analysis summary statistics generated during the study are available from GWAS catalog (study accession GCST90013791). Autoimmune thyroid disease GWAS data are available from UK Biobank.

## Code availability

An R package for BFDR estimation, priorsplitteR, is available from https://github.com/djmcrouch/priorsplitteR. Analysis code is available from the authors on request.

## Acknowledgements

This work was funded by JDRF grants 9-2011-253, 5-SRA-2015-130-A-N and 4-SRA-2017-473-A-N, and Wellcome grants 091157/Z/10/Z and 107212/Z/15/Z, to the Diabetes and Inflammation Laboratory, Oxford.

Computational work was performed using the Oxford Biomedical Research Computing (BMRC) facility, a joint development between the Wellcome Centre for Human Genetics and the Big Data Institute supported by Health Data Research UK and the NIHR Oxford Biomedical Research Centre. Financial support was provided by the Wellcome Trust Core Award grant 203141/Z/16/Z. The views expressed are those of the author(s) and not necessarily those of the NHS, the NIHR or the Department of Health

Work was supported from grant U1301.2015/AI.1157.BE from Fondazione di Sardegna to Francesco Cucca.

We acknowledge the participants and investigators of the FinnGen study.

This study makes use of data generated by the Wellcome Trust Case Control Consortium, funded by Wellcome Trust award 076113; a full list of the investigators who contributed to the generation of the data is available from http://www.wtccc.org.uk/.

This research uses resources provided by the Type 1 Diabetes Genetics Consortium, a collaborative clinical study sponsored by the National Institute of Diabetes and Digestive and Kidney Diseases (NIDDK), the National Institute of Allergy and Infectious Diseases (NIAID), the National Human Genome Research Institute (NHGRI), the National Institute of Child Health and Human Development (NICHD) and JDRF and supported by US NIH grant U01 DK062418, NIDDK grant DP3 DK111906 and US National Library of Medicine grant T32 LM012416. We acknowledge the participants and members of the T1DGC (see Supplementary note for a list of members), and T1DGC steering group members Beena Akolkar, Henry A Erlich, Cécile Julier, Grant Morahan, Jørn Nerup and Concepcion Nierras.

We thank Walter Bodmer for discussions related to the BFDR method.

## Author contributions

The project was conceived by J.A.T. and S.S.R.

Genotype data processing, quality control, imputation, and statistical analyses were performed by D.J.M.C., J.R.J.I, C.C.R., J-Y.Y., W-M.C., S.O.G., and C.S.

New statistical methods were designed and tested by D.J.M.C.

T1DGC DNA samples for genotyping were managed by S.O.G.

C.S. and F.C contributed genotype data for the Sardinian cohort

F.P. and P.C. provided samples for genotyping through their affiliated institutions and research programs.

Biological interpretation of results was provided by A.J.C and J.A.T.

The manuscript was written by D.J.M.C. and J.A.T. with additional material from J.R.J.I. and C.C.R.

