## Supplementary Figures for "Enhanced genetic analysis of type 1 diabetes by selecting variants on both effect size and significance, and by integration with autoimmune thyroid disease"

**Figure S1**

Manhattan plots of GWAS associations for a) T1D and b) ATD. Blue lines indicate P values corresponding to the 1% FDR threshold in each disease, and red lines the genome-wide significant level of $5{\times}10^{-8}$. Lead SNPs for each independent region are highlighted in green. Lead SNPs lying below the blue line have been selected due to having large effect sizes (BFDR<5%, see text). Figure produced using the qqman R package (Turner, 2018).

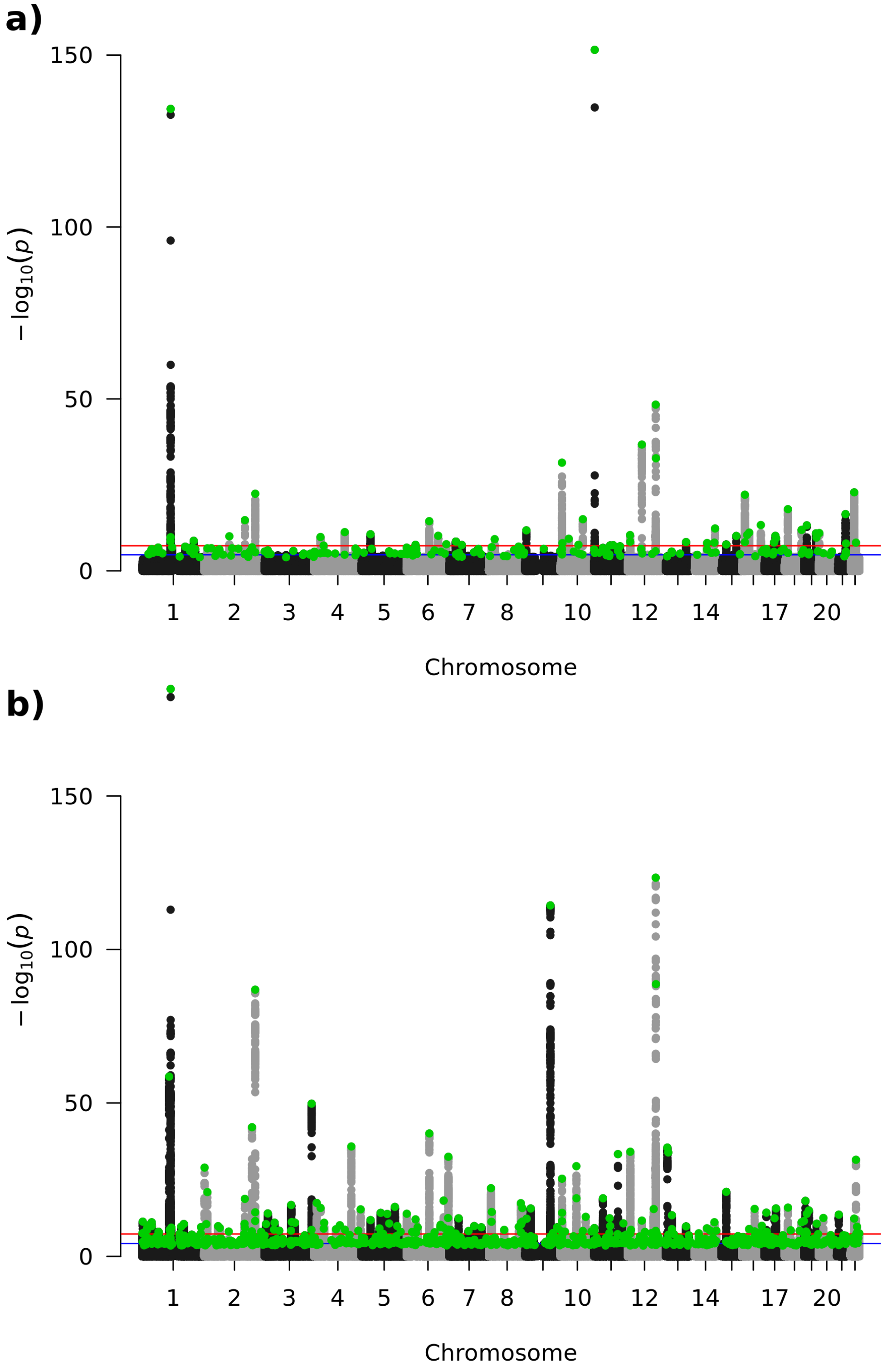

**Figure S2**

Quantile-quantile plots of observed versus expected P values under the null hypothesis, excluding HLA region, for a) T1D ($\lambda_{GC}$ = 1.12) and b) ATD ($\lambda_{GC}$ = 1.19). Genomic control inflation factors ($\lambda_{GC}$) were produced by dividing the median Chi-square by the median under the null hypothesis (0.456).

**a)**

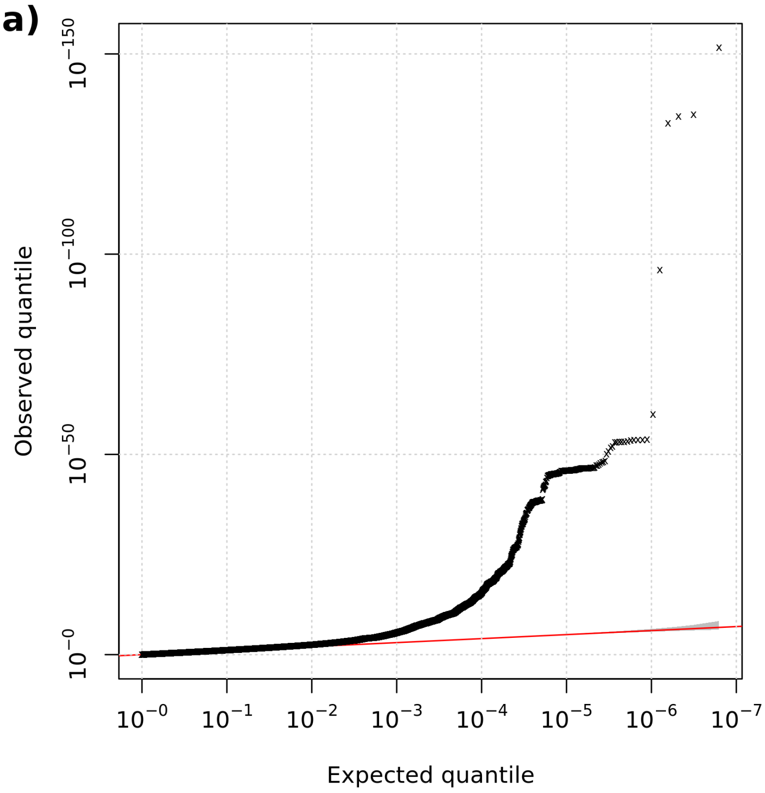


**b)**

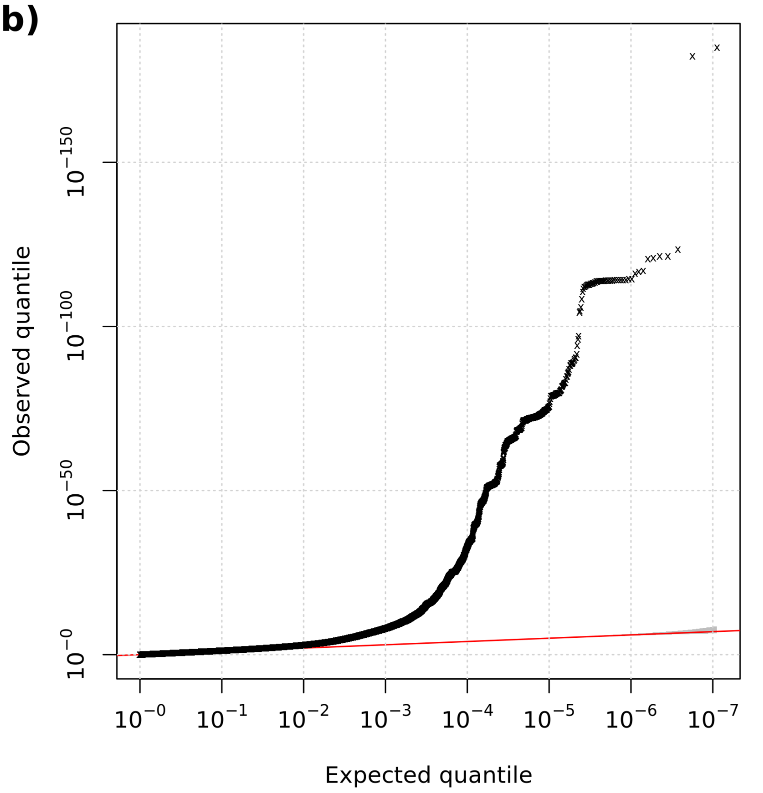

**Figure S3**

Volcano plots of minor allele effect size (log odds ratio) versus significance (-log10 P values) for a) type 1 diabetes and b) autoimmune thyroid disease. All analysed variants are shown. Variants with either i) Bigger or False Discovery Rate (BFDR)<1% or ii) FDR<1% were designated as associations. See Figure 3 (main text) for the same volcano plots highlighting variants with BFDR<1% or FDR<1%.

**a)**

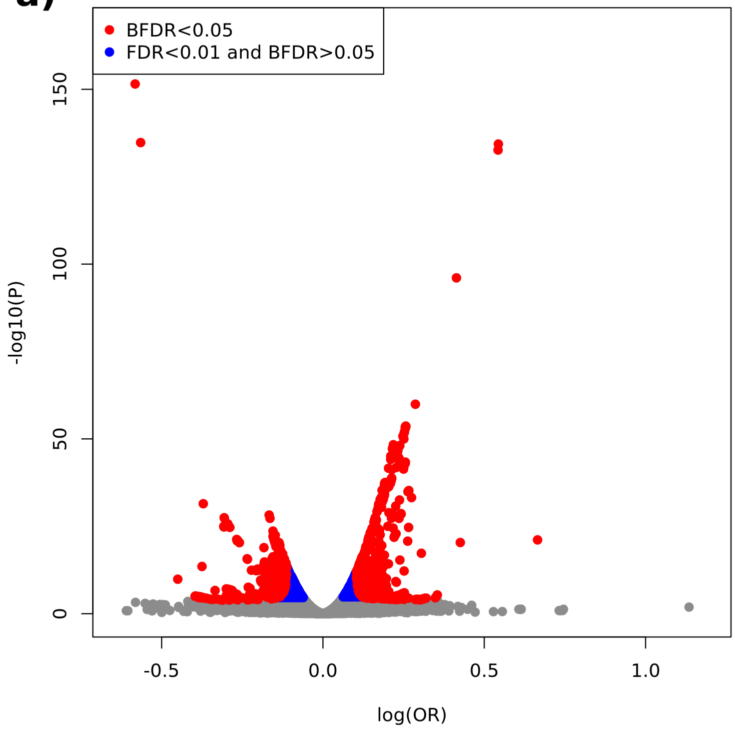


**b)**

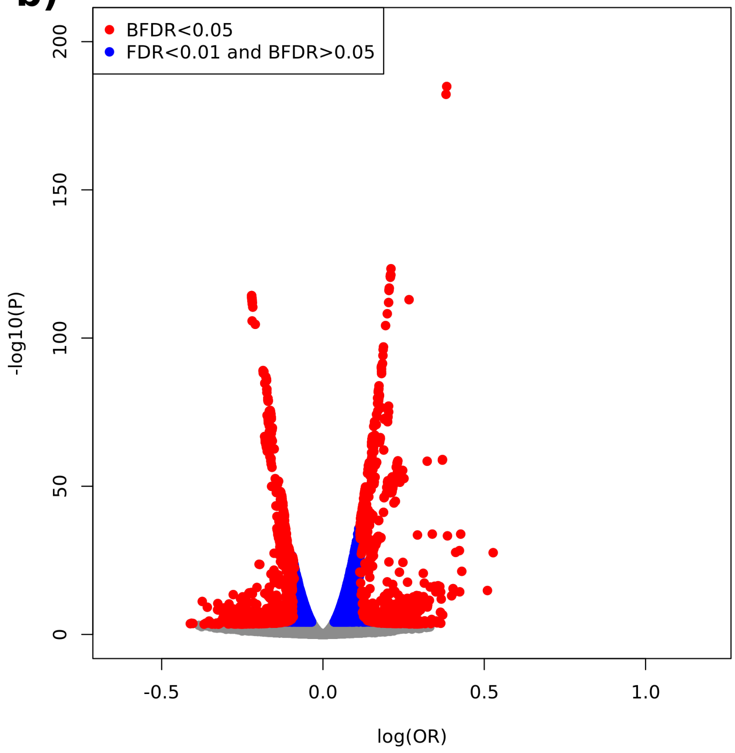

**Figure S4**

Numbers of new and previously reported signals satisfying BFDR<1%, FDR<1% and genome-wide (P<5×10$^{-8}$) significance criteria for a) type 1 diabetes and b) autoimmune thyroid disease, quantified using the lead variant in each signal. Figure 4 (main text) shows the equivalent Venn diagrams for results using a 5% BFDR threshold.

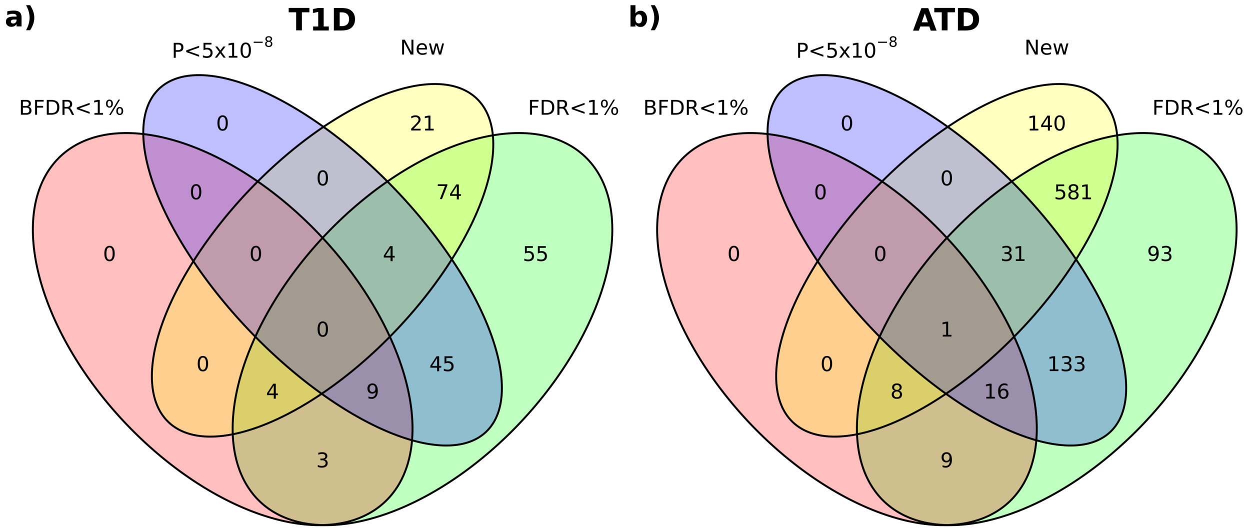

**Figure S5**

Fine-mapping results for previously unreported signals with BFDR<1% (plus *RAD51D,* BFDR=1.36% and SH3BP4, BFDR=2.80%). The disease associated with each signal is given in parentheses. Credible sets are shown by coloured blocks, with the height of the blog representing the cumulative log10 Bayes Factor of SNPs in the set. Asterisks denote the location of the lead GWAS variant (coloured by credible set). Full fine-mapping data is provided in Tables S8 and S9.

a) *RPL7P10* (T1D)

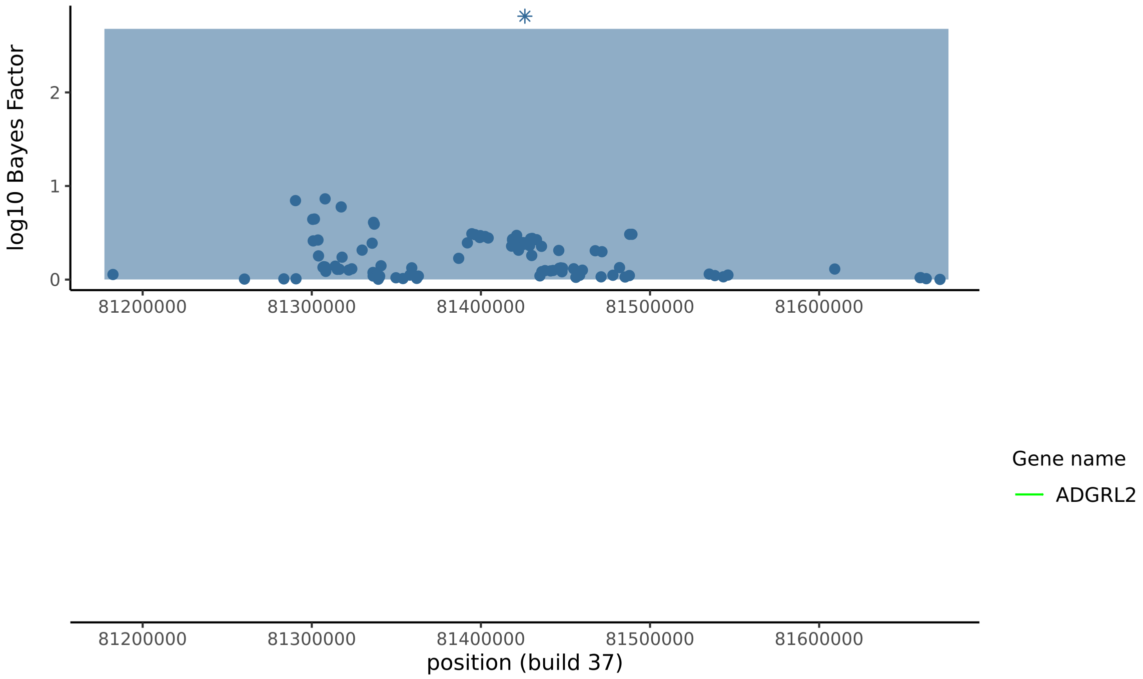


b) *ID4* (T1D)

Lead variant not present in any credible set.

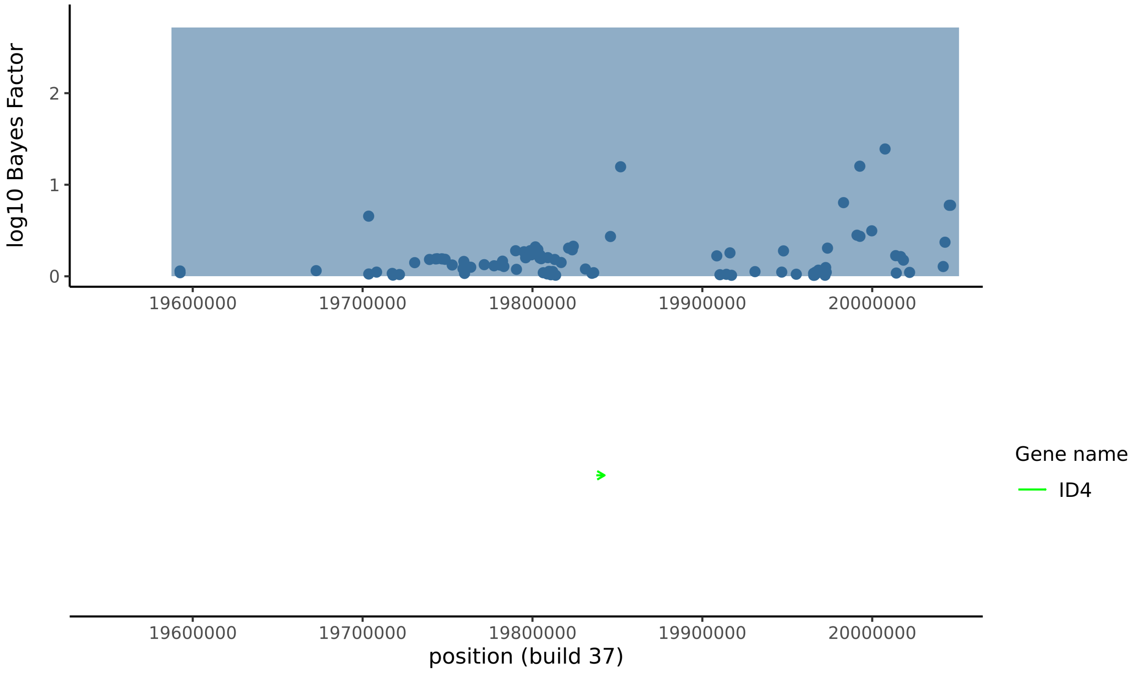

c) *NRSN1* (T1D)

Lead variant not present in any credible set.

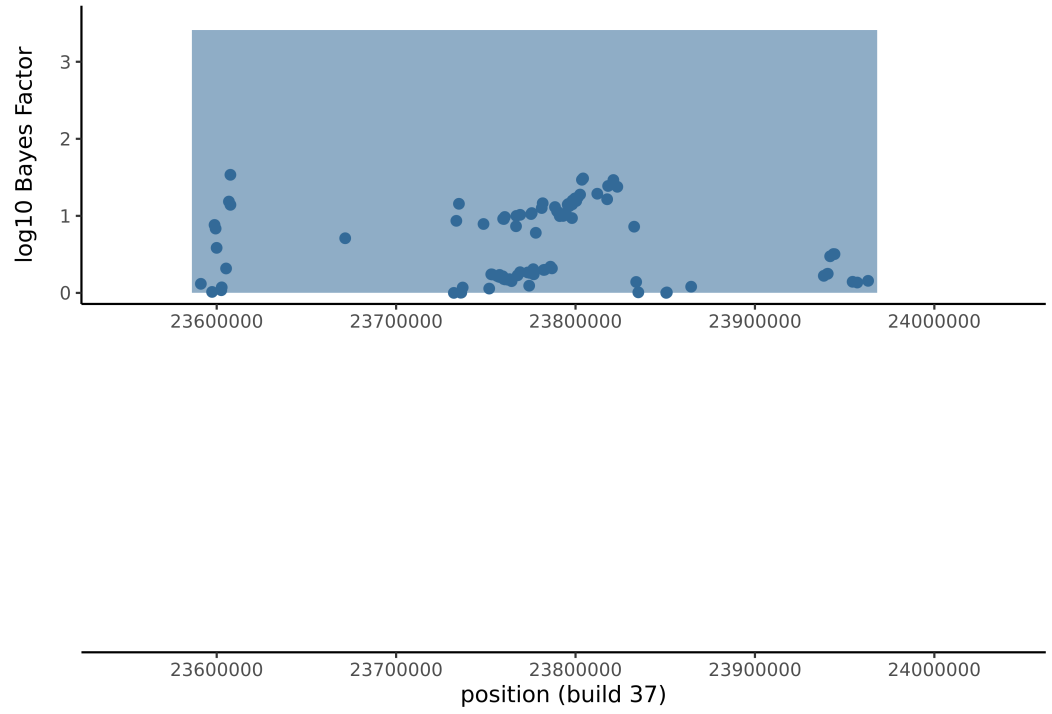


d) *RPL35AP21* (T1D)

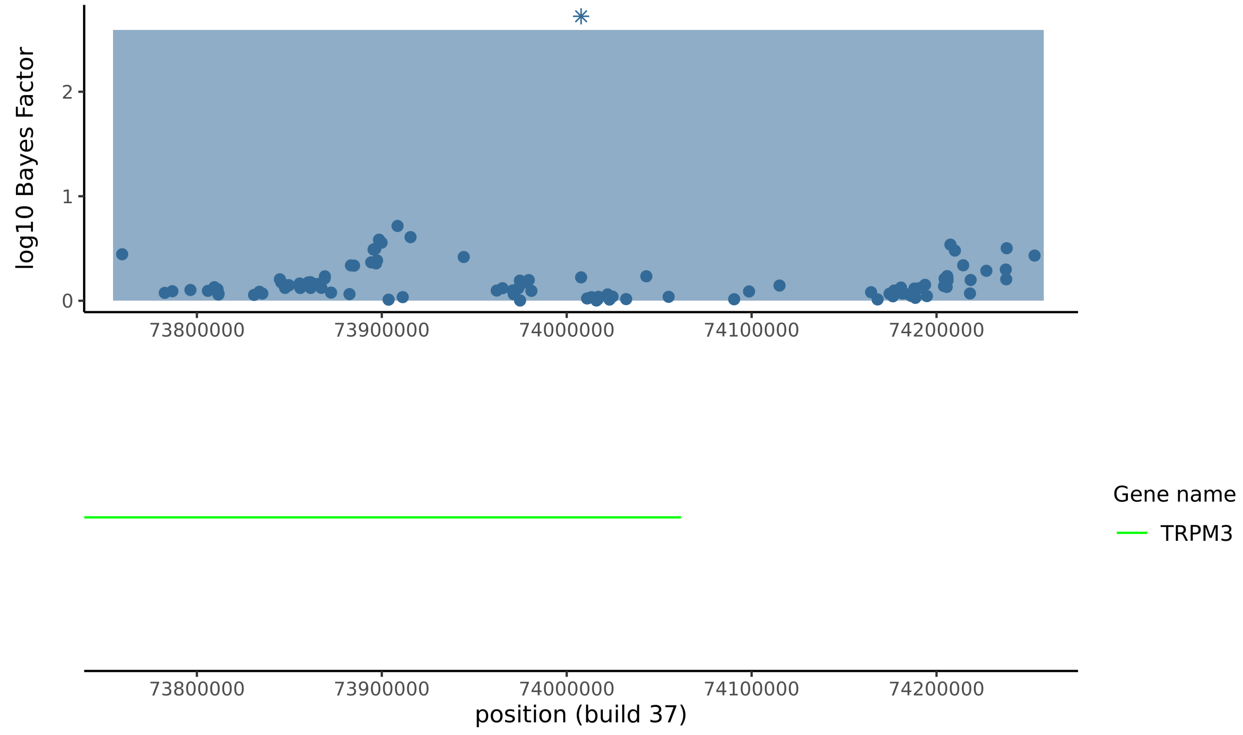

e) *RAD51D* (T1D)

FINEMAP could not distinguish between SNPs in the credible set, but three 3'UTR variants had log10 Bayes factors that were somewhat higher than other variants in the region (see also Table S8).

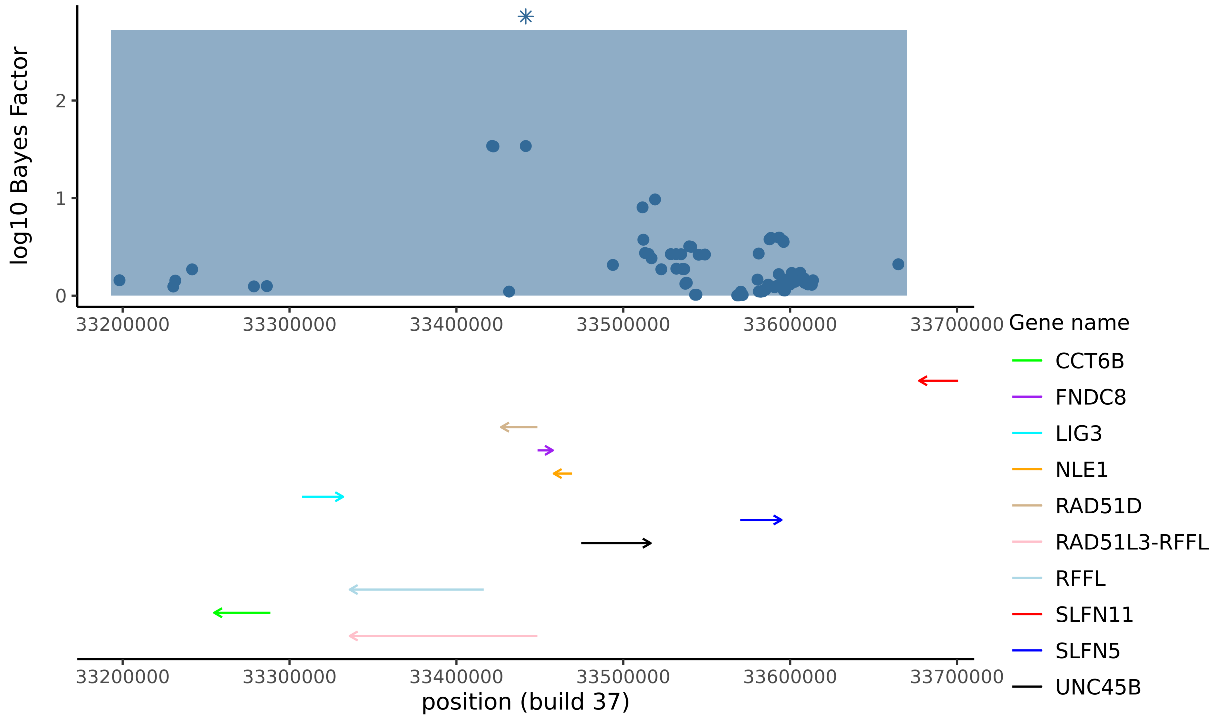


f) *RAD51B* (T1D)

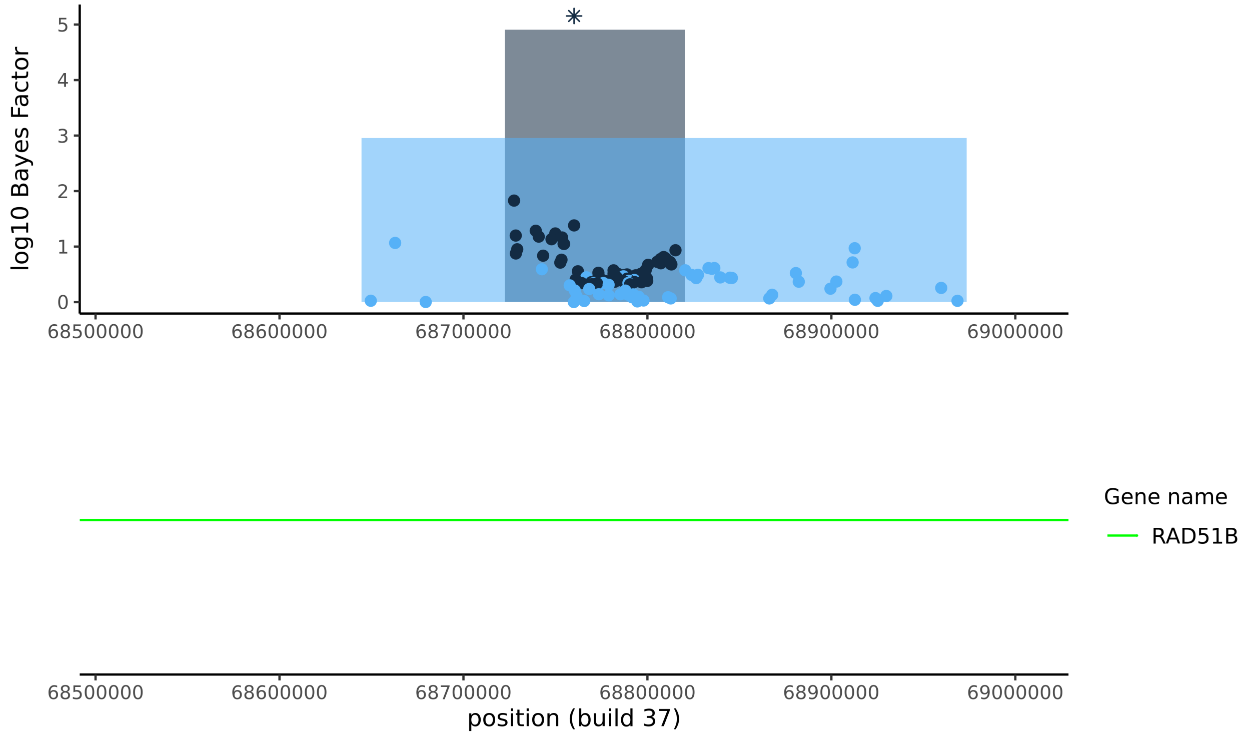

g) *MAGI3* (ATD)

The group with high posterior log odds on the right hand side is associated with the well know *PTPN22* lead variant, rs2476601. Our novel MAGI3 association is not represented in any of these three credible sets.

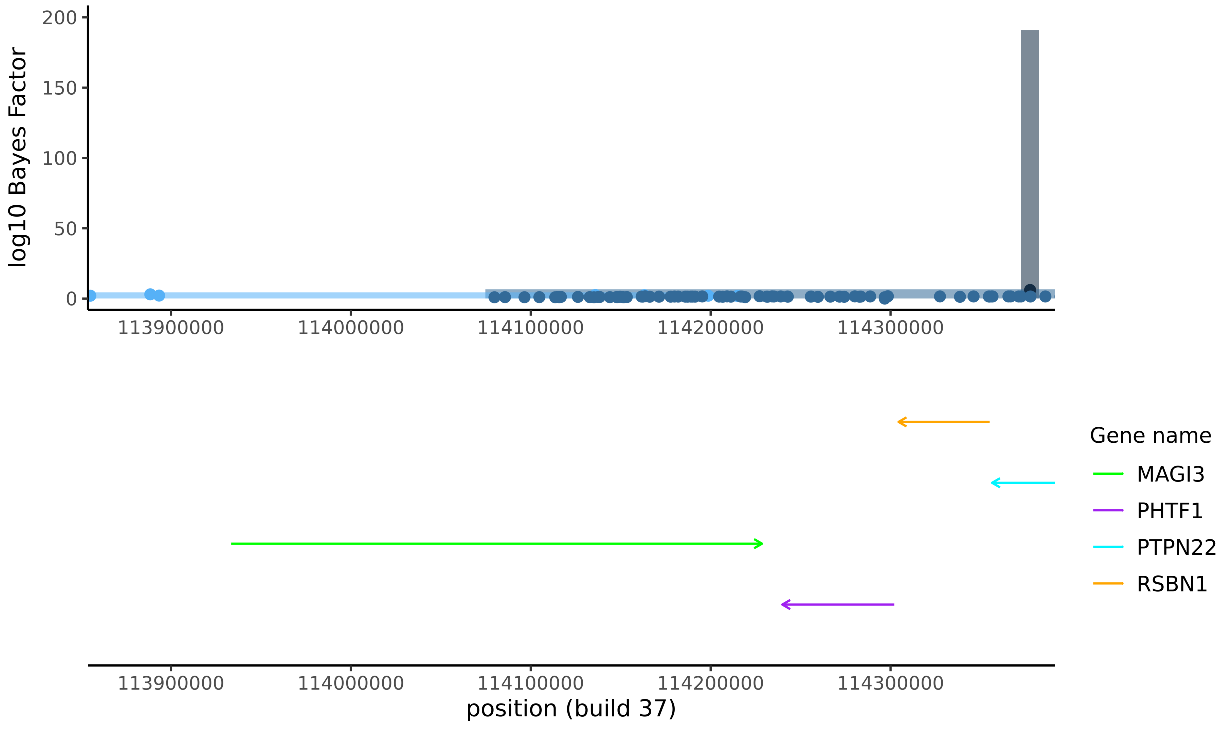


h) *RAPH1* (ATD)

Lead variant not present in any credible set.

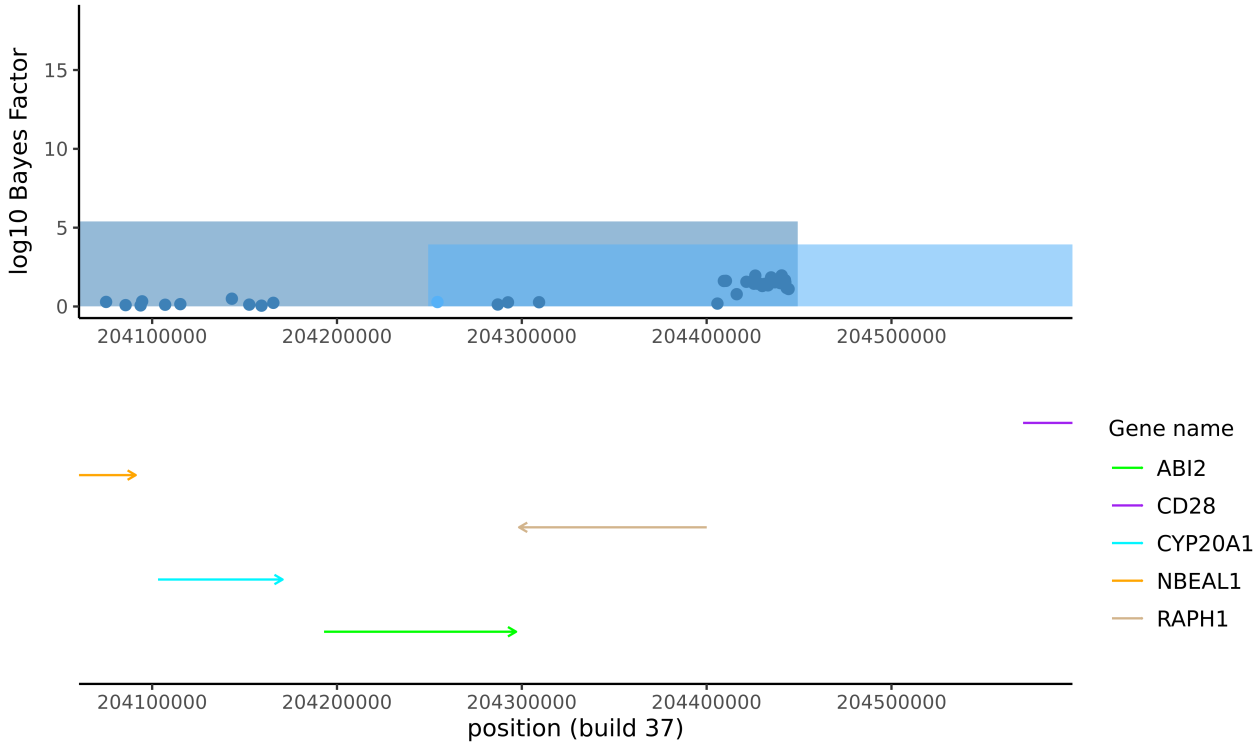

i) *GRIK2* (ATD)

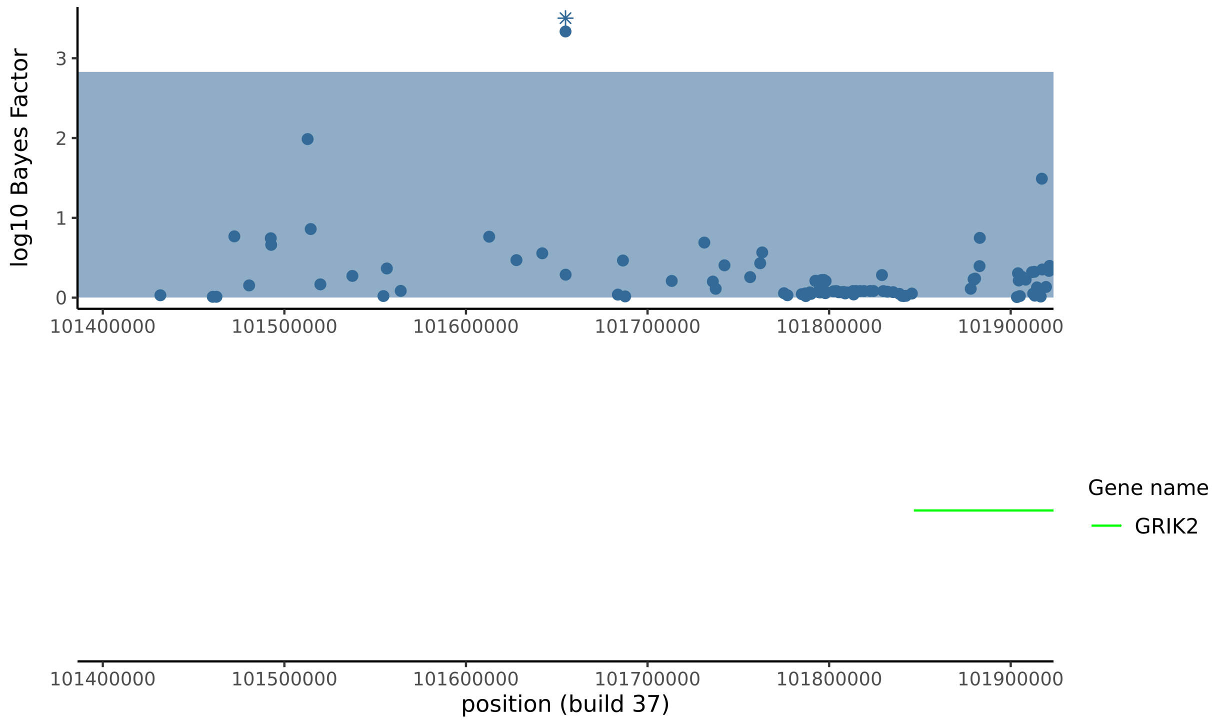


j) *IL6* (ATD)

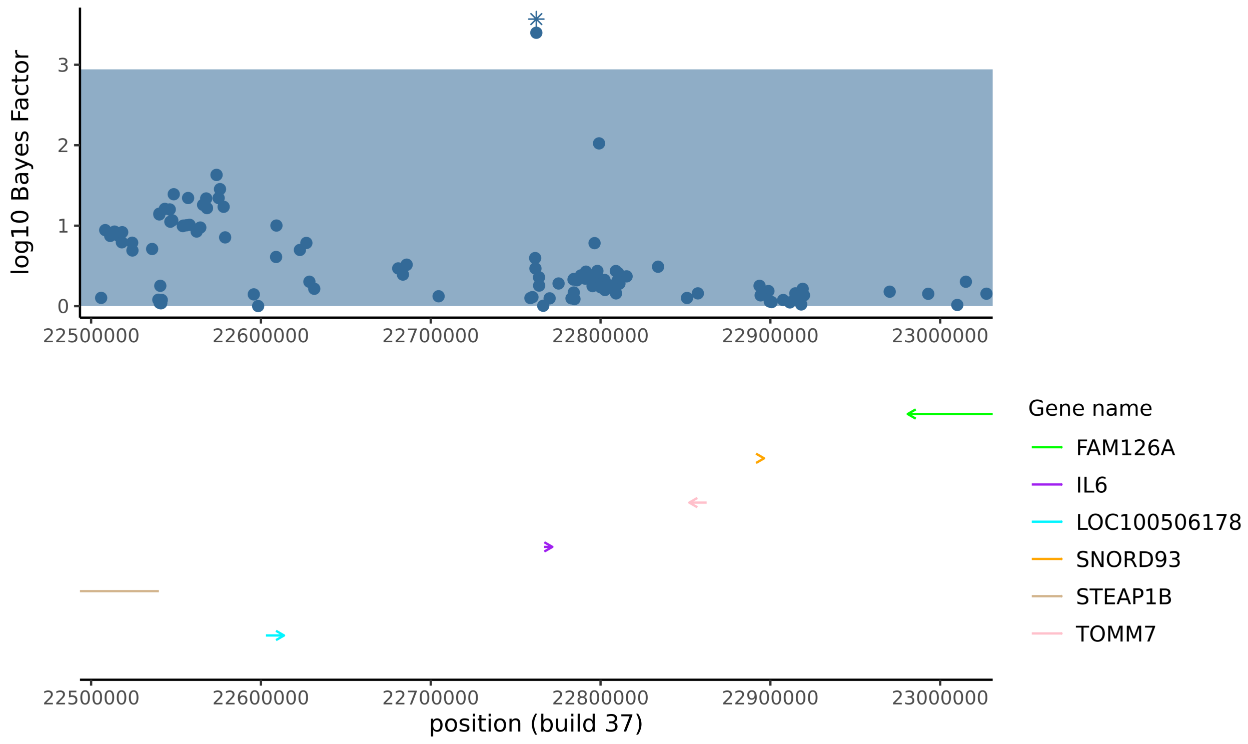

k) *BMPR2* (ATD)

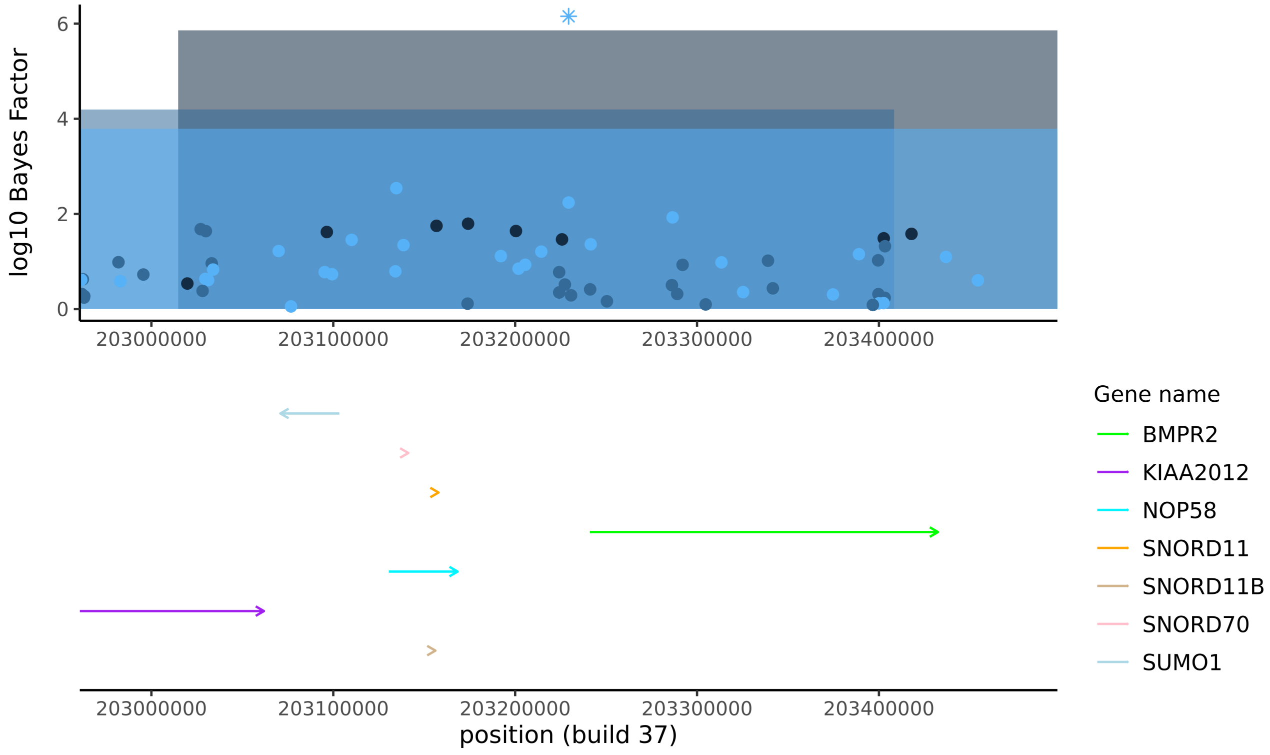


l) *PTPRF* (ATD)

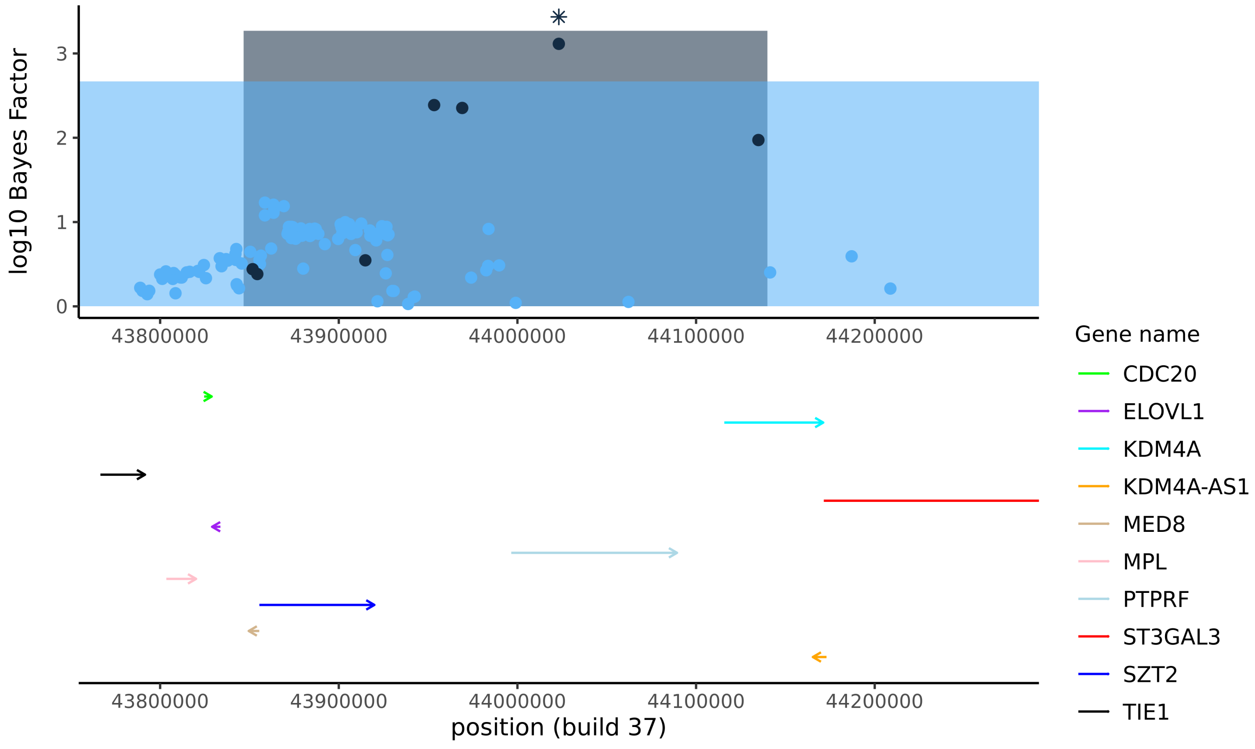

m) *FAM117B* (ATD)

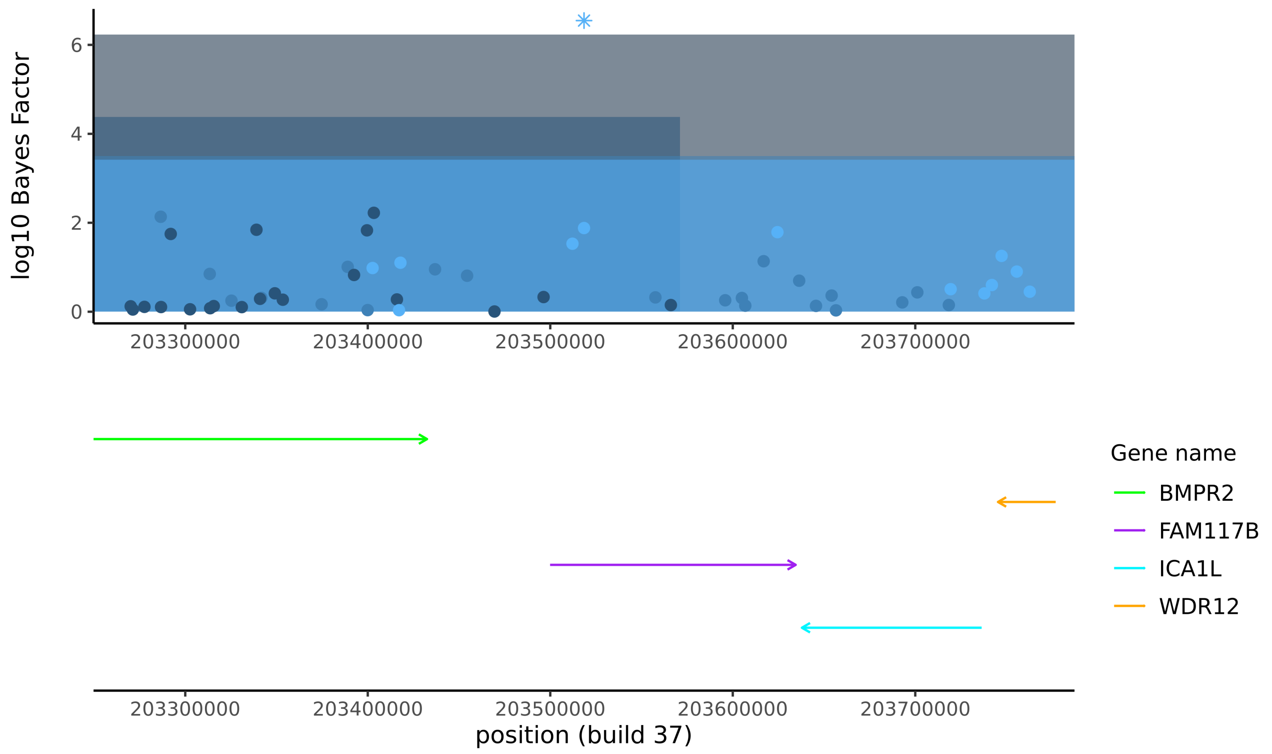


n) *CYP1B1-AS1* (ATD)

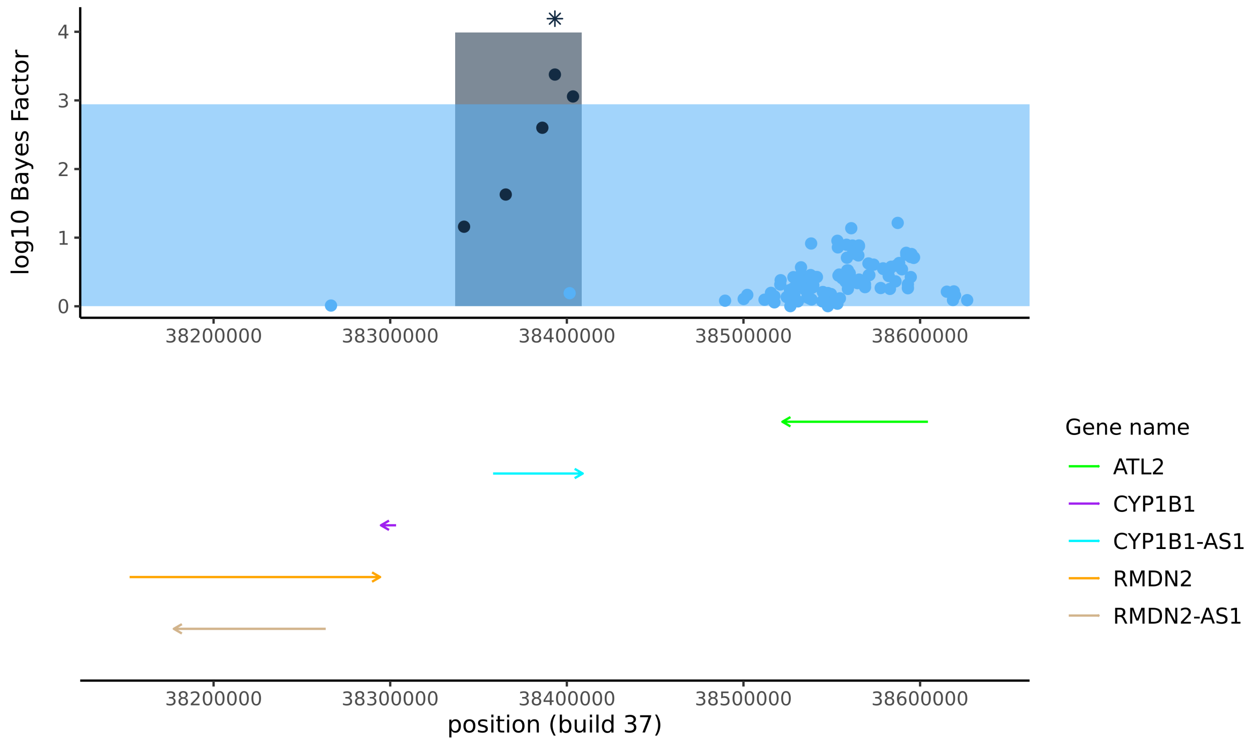

o) *MEOX2* (ATD)

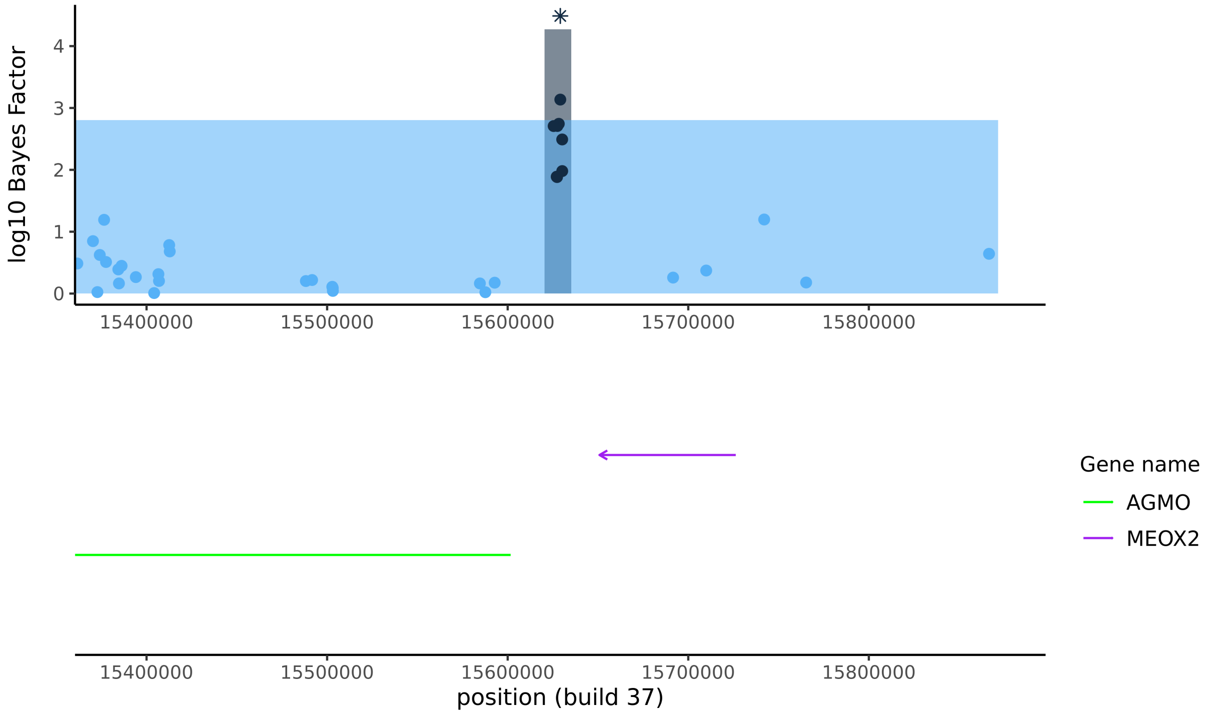


p) *SH3BP4* (ATD)

FINEMAP could not distinguish between SNPs in the credible set, but two deletion variants had log10 Bayes factors that were somewhat higher than other variants in the region (see also Table S9).

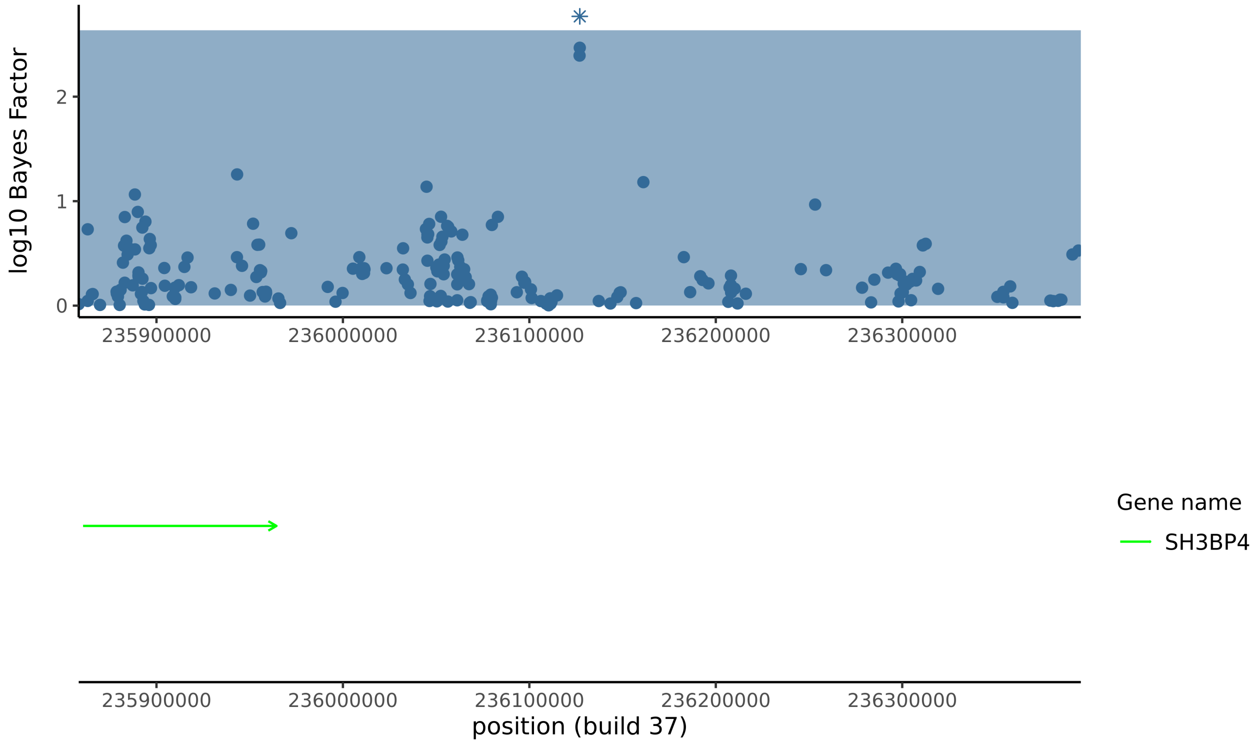

**Figure S6**

Colocalisation data for four previously unreported T1D BFDR<1% signals, *RAD51D* and *RAD51B*, and five ATD signals with both BFDR<1% and MAF≥3%. See Tables S5-S7 for colocalisation summary data from all signals. Diseases that each signal was found to associate with in the primary GWAS analysis are given in parenthesis.

a) *RPL7P10* (T1D)

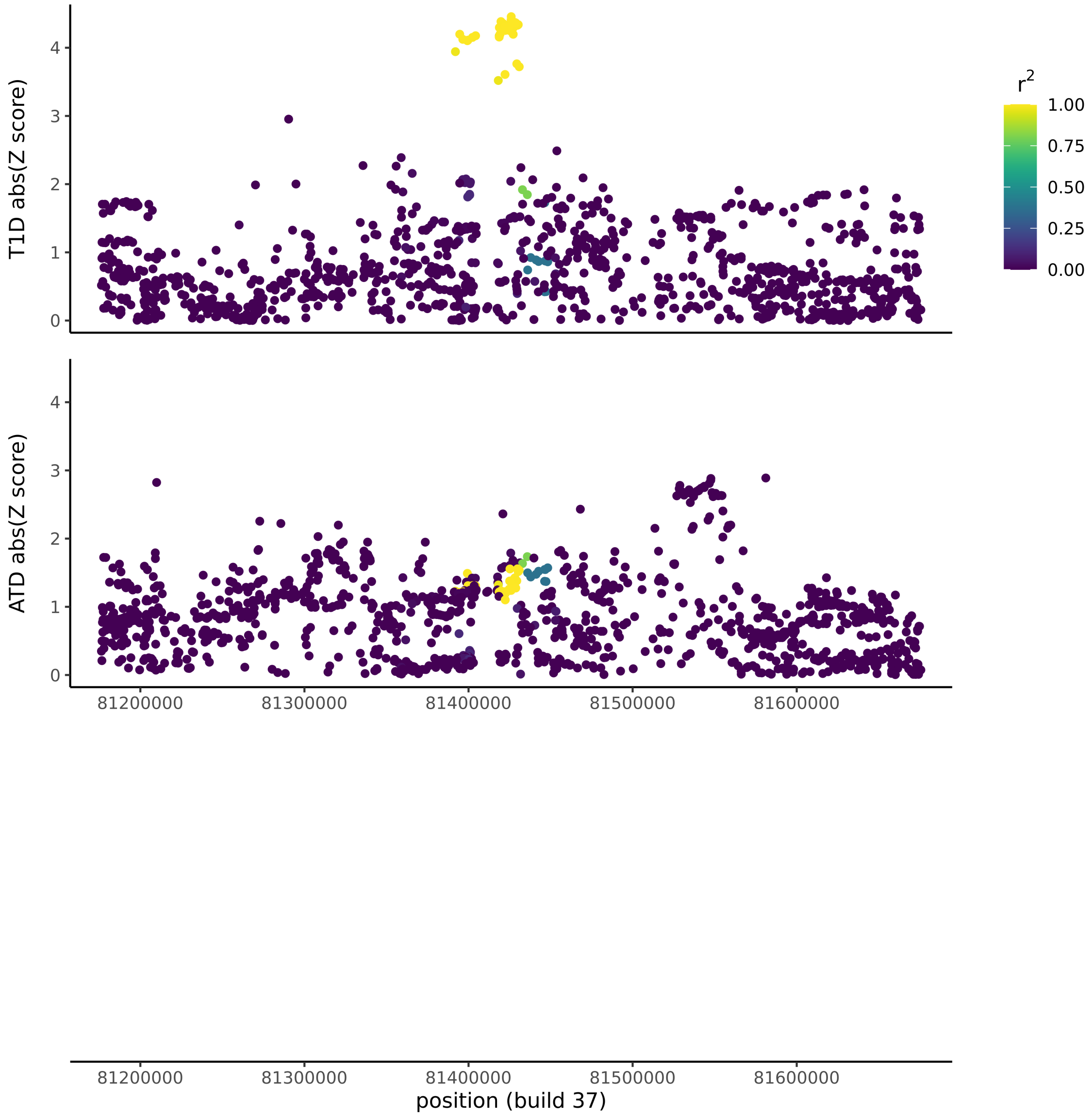

b) *ID4* (T1D)

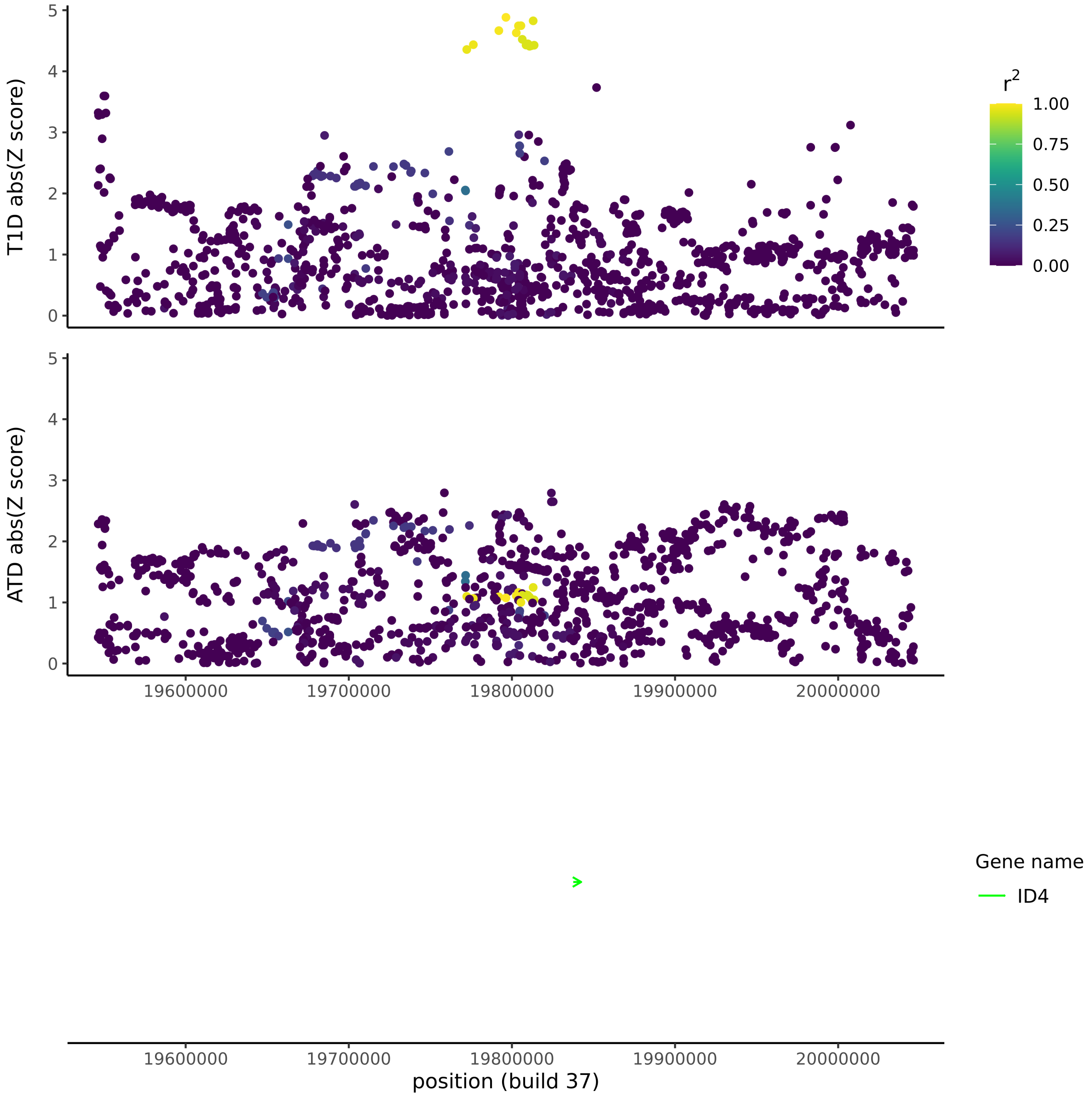
T1D abs(Z score)
ATD abs(Z score)
r2
1.00
0.75
0.50
0.25
0.00
19600000
19700000
19800000
19900000
20000000
Gene name
ID4
position (build 37)

c) *NRSN1* (T1D)

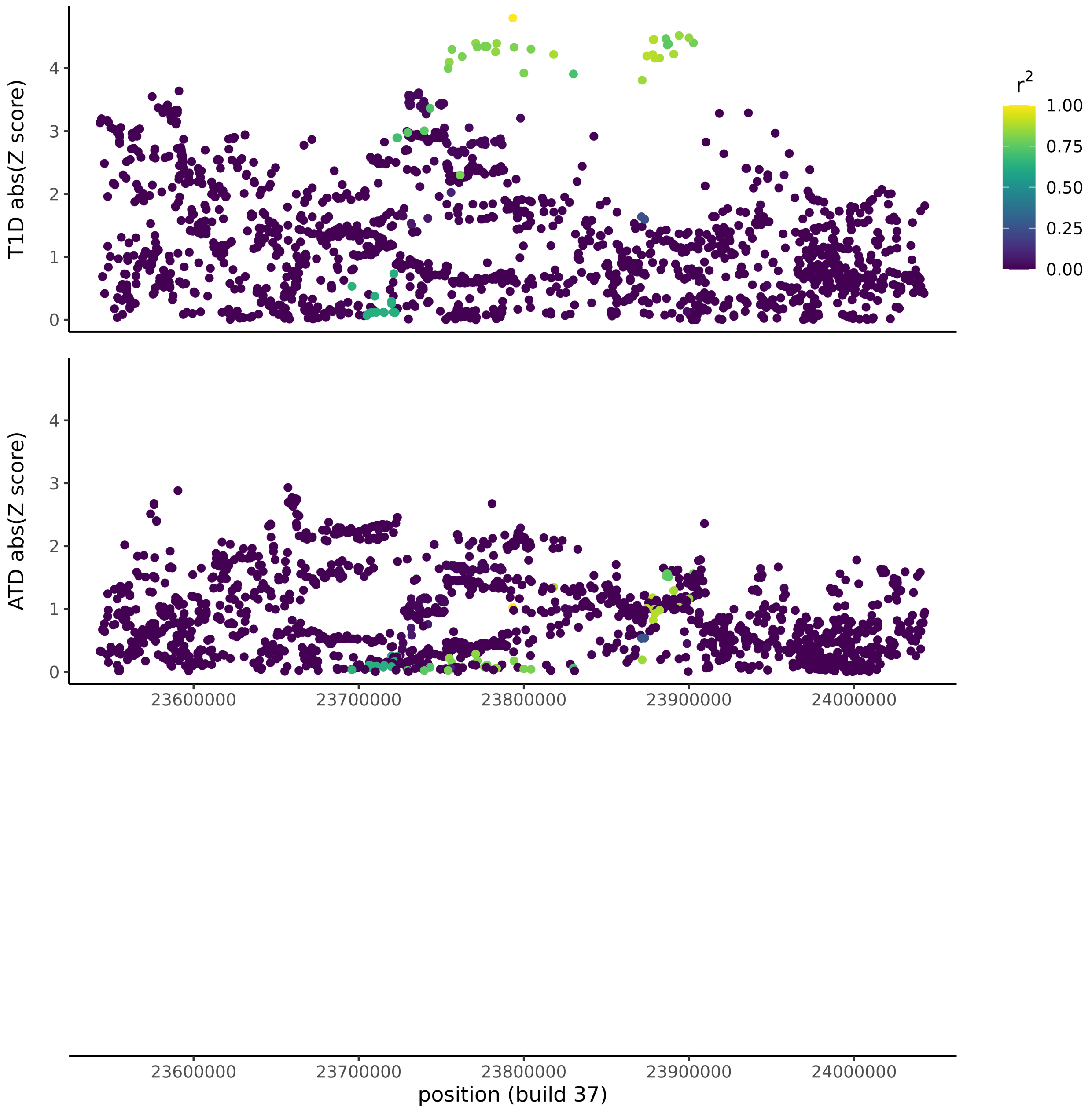
T1D abs(Z score)
ATD abs(Z score)
0
1
2
3
4
23600000
23700000
23800000
23900000
24000000
position (build 37)
r²
1.00
0.75
0.50
0.25
0.00

d) *RPL35AP21* (T1D)

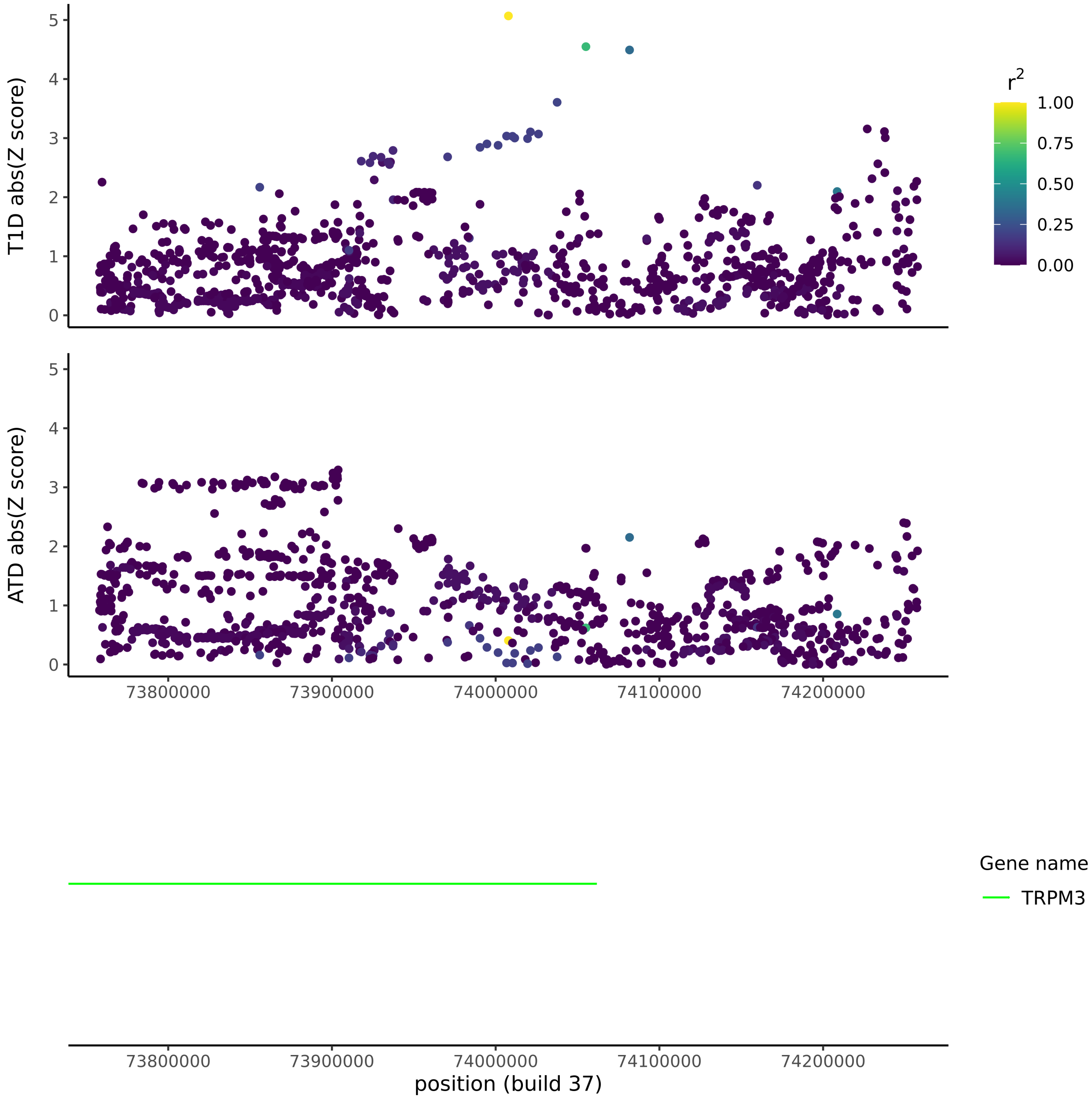

e) *RAD51D* (T1D)

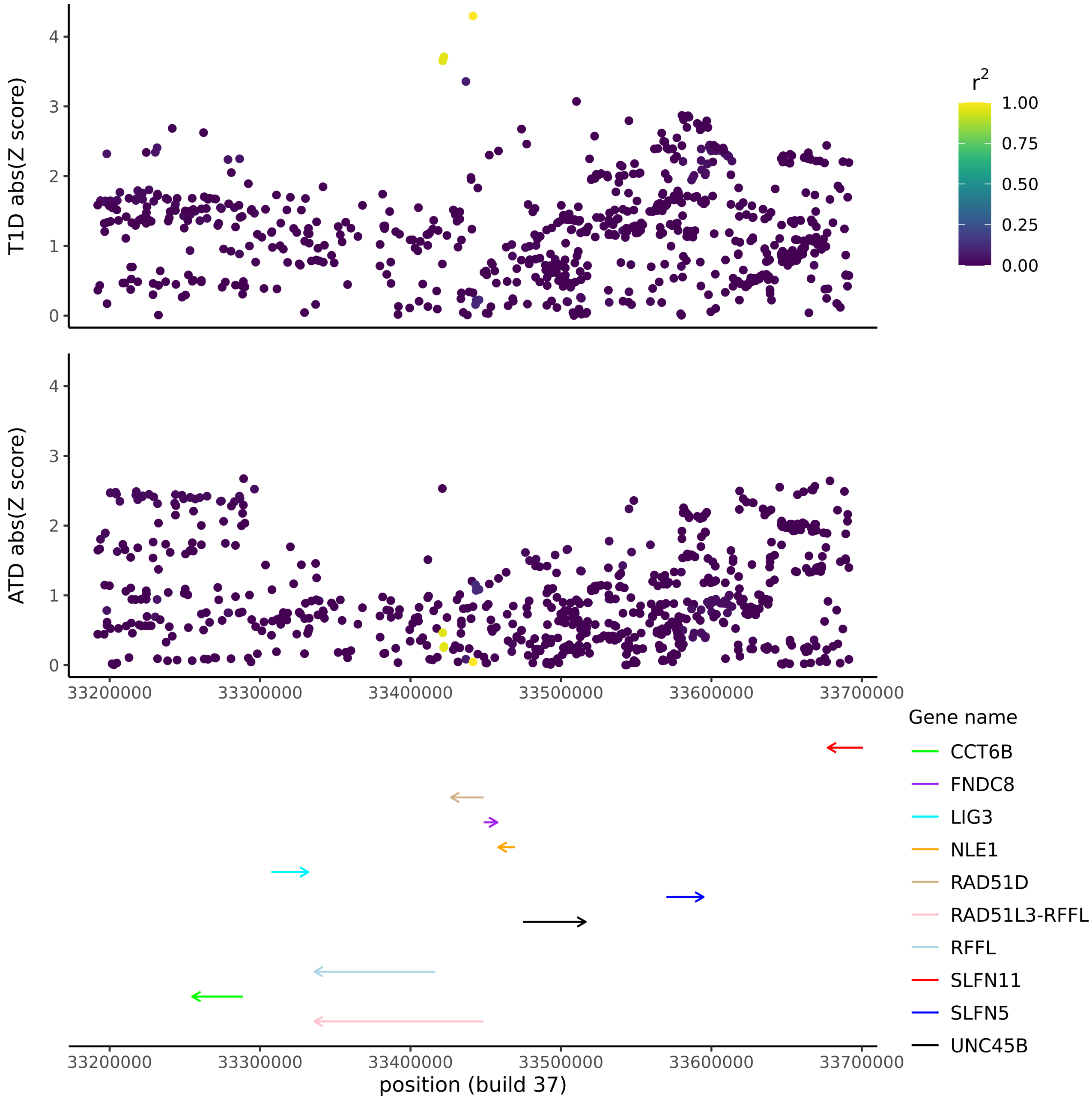
T1D abs(Z score)
ATD abs(Z score)
r2
1.00
0.75
0.50
0.25
0.00
Gene name
CCT6B
FNDC8
LIG3
NLE1
RAD51D
RAD51L3-RFFL
RFFL
SLFN11
SLFN5
UNC45B
33200000
33300000
33400000
33500000
33600000
33700000
position (build 37)

f) *RAD51B* (T1D)

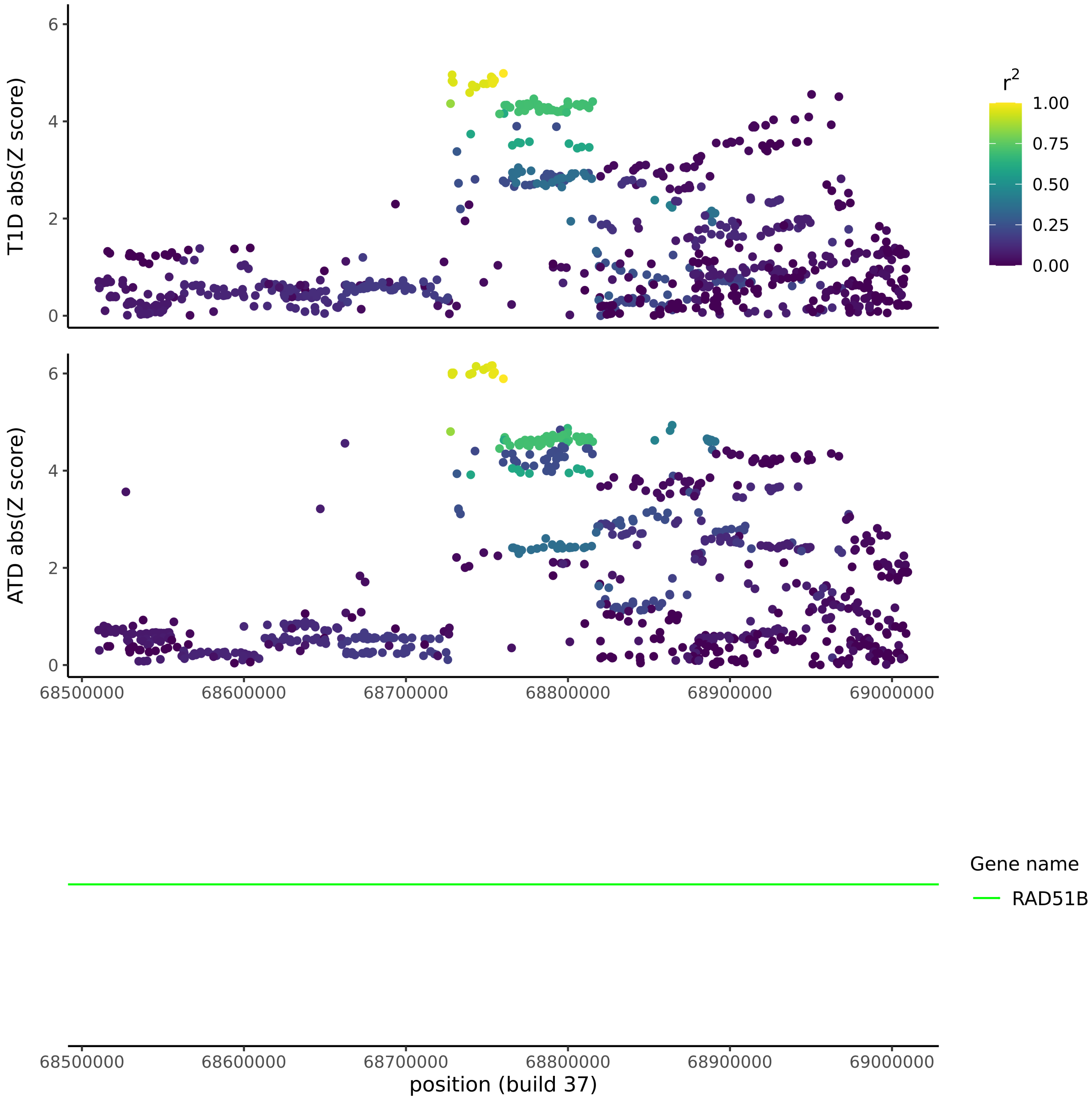
T1D abs(Z score)
ATD abs(Z score)
r2
1.00
0.75
0.50
0.25
0.00
Gene name
RAD51B
68500000
68600000
68700000
68800000
68900000
69000000
position (build 37)

g) *VAV3* (ATD)

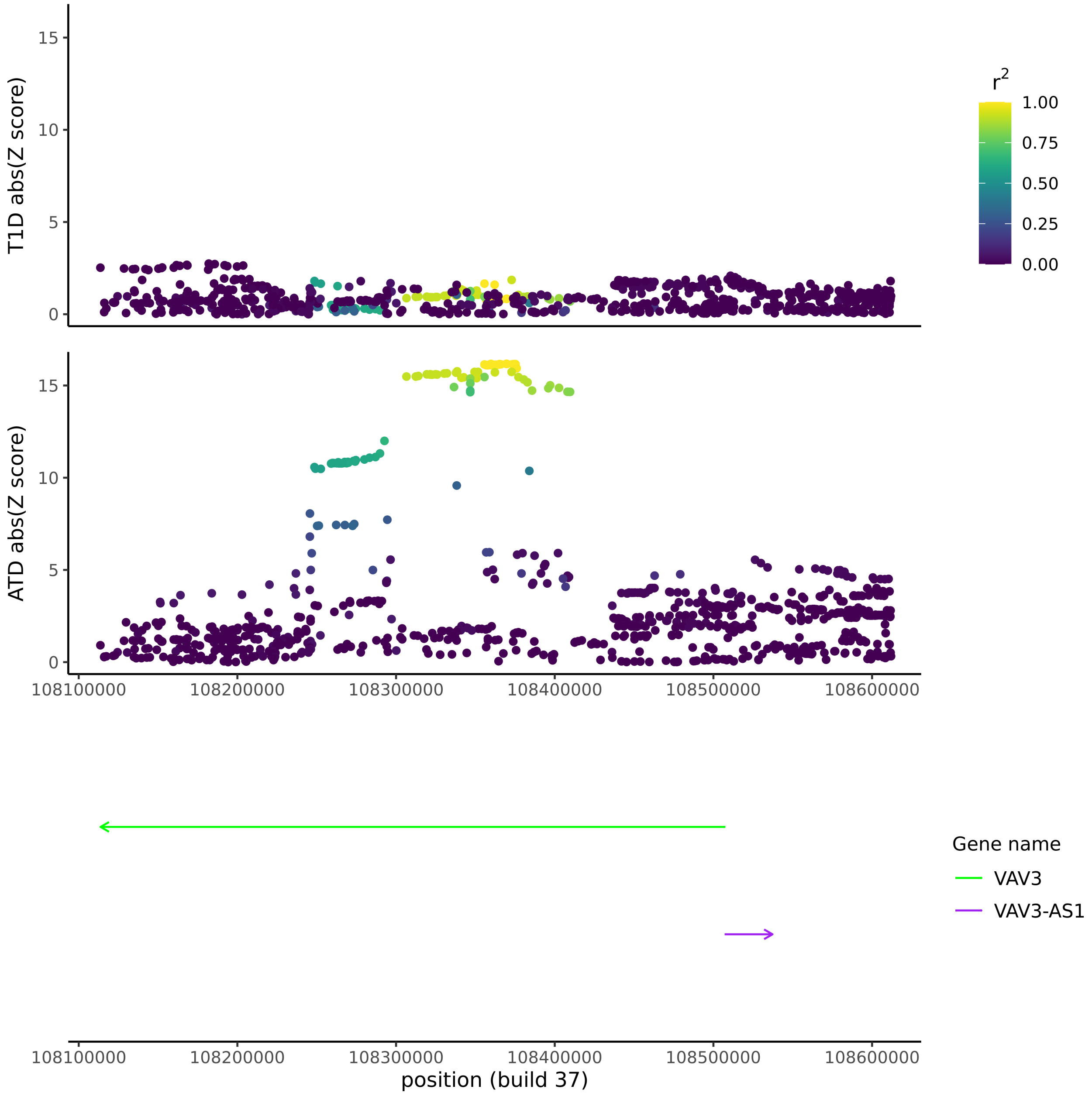
T1D abs(Z score)
ATD abs(Z score)
0
5
10
15
108100000
108200000
108300000
108400000
108500000
108600000
position (build 37)
r2
1.00
0.75
0.50
0.25
0.00
Gene name
VAV3
VAV3-AS1

h) *FOXE1* (ATD)

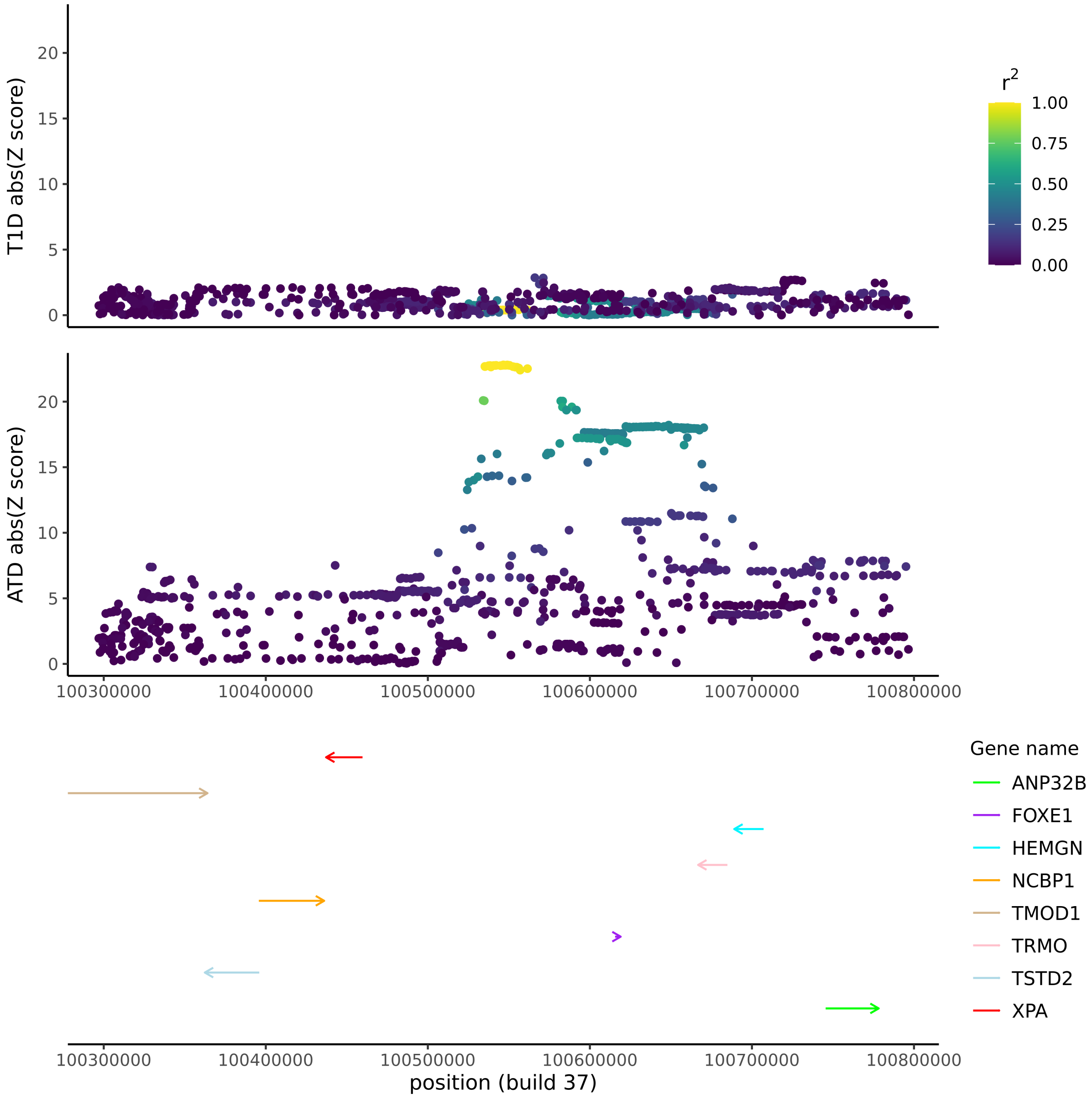

T1D abs(Z score)
ATD abs(Z score)
0
5
10
15
20
r2
1.00
0.75
0.50
0.25
0.00
100300000
100400000
100500000
100600000
100700000
100800000
Gene name
ANP32B
FOXE1
HEMGN
NCBP1
TMOD1
TRMO
TSTD2
XPA
position (build 37)

i) *PTPN22* (ATD)

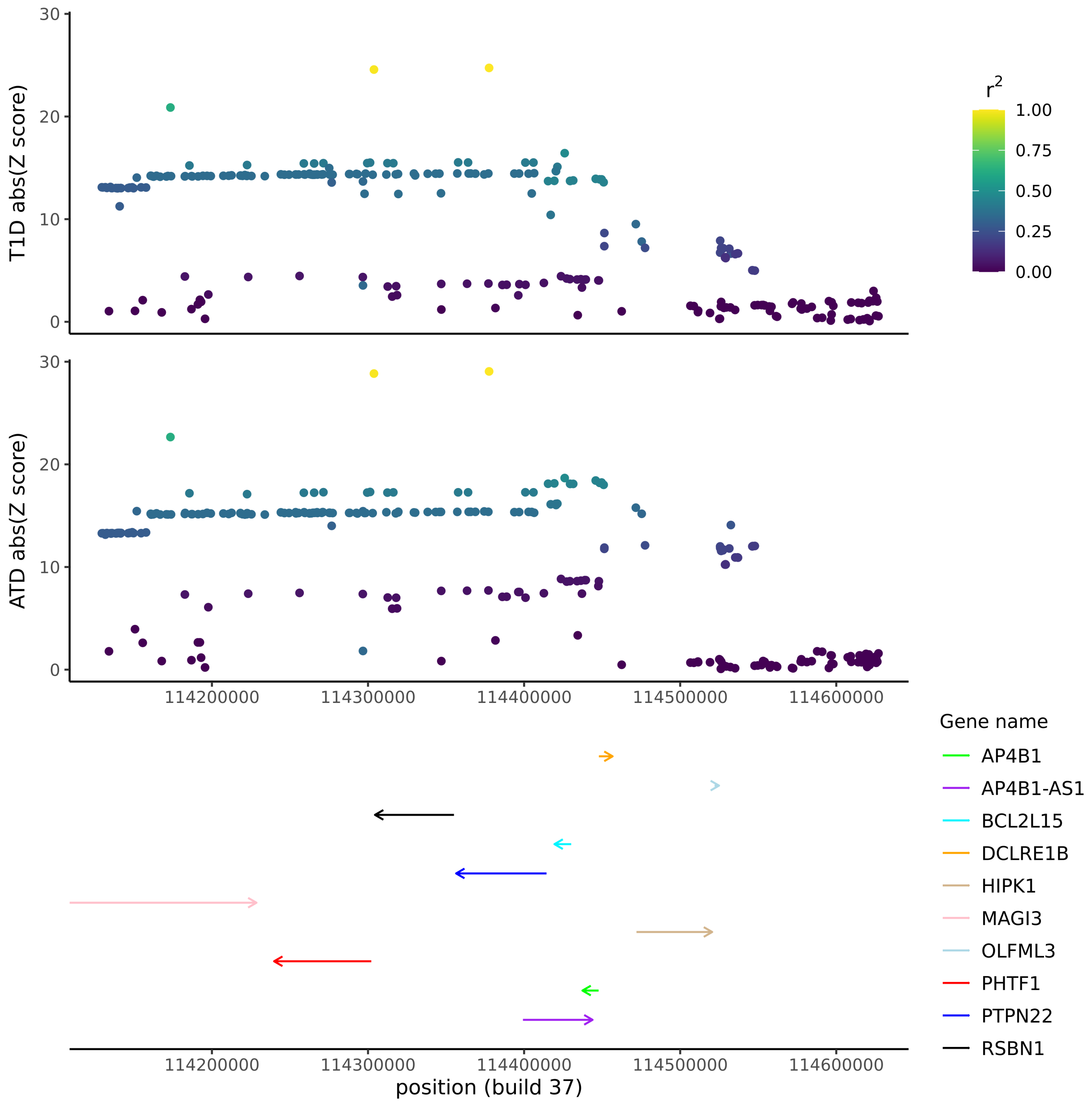
T1D abs(Z score)
ATD abs(Z score)
30
20
10
0
r2
1.00
0.75
0.50
0.25
0.00
114200000
114300000
114400000
114500000
114600000
position (build 37)
Gene name
AP4B1
AP4B1-AS1
BCL2L15
DCLRE1B
HIPK1
MAGI3
OLFML3
PHTF1
PTPN22
RSBN1

j) *SH2B3* (ATD)

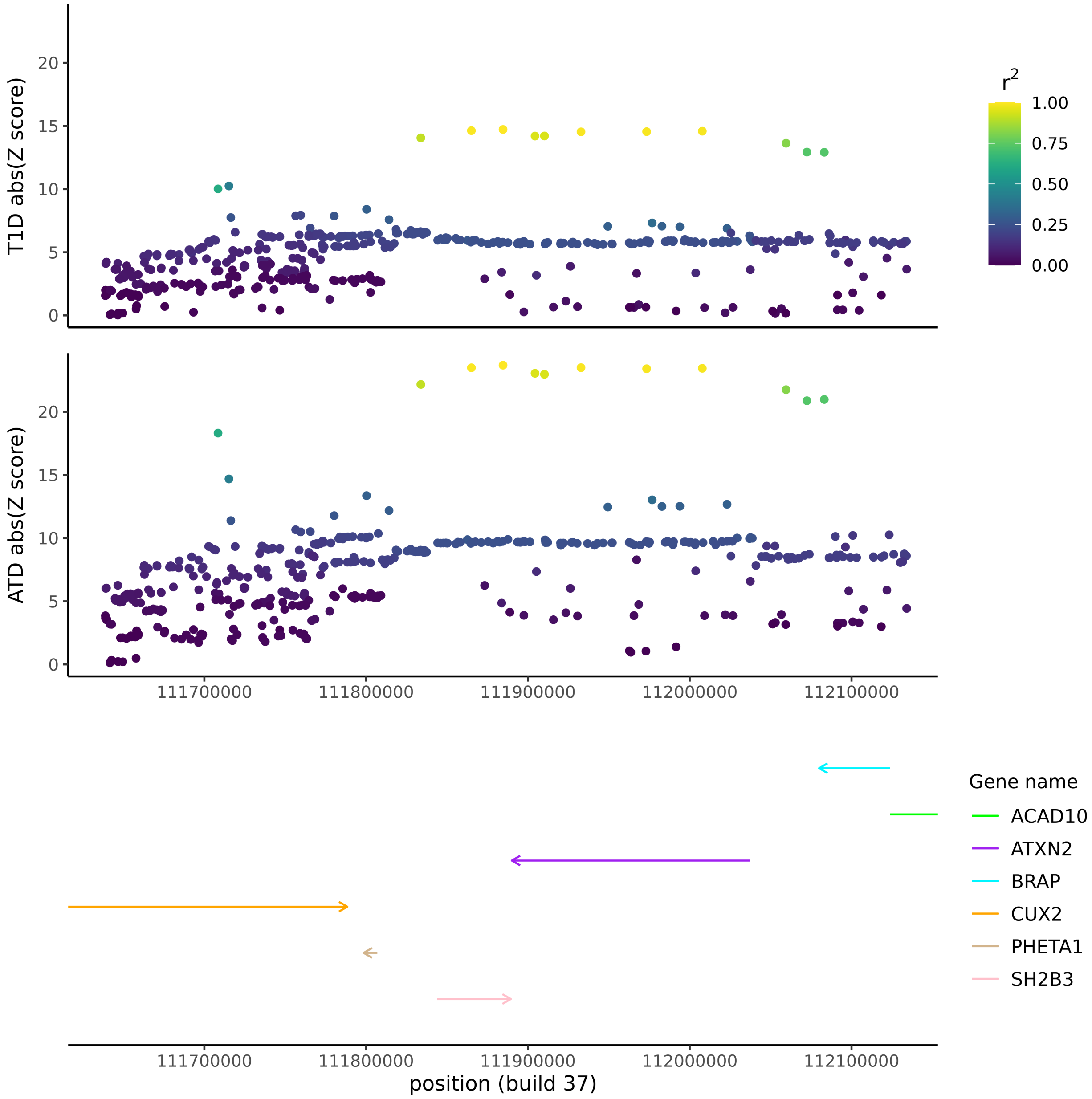
T1D abs(Z score)
ATD abs(Z score)
0
5
10
15
20
r2
1.00
0.75
0.50
0.25
0.00
111700000
111800000
111900000
112000000
112100000
Gene name
ACAD10
ATXN2
BRAP
CUX2
PHETA1
SH2B3
position (build 37)

k) *CTLA4* (ATD)

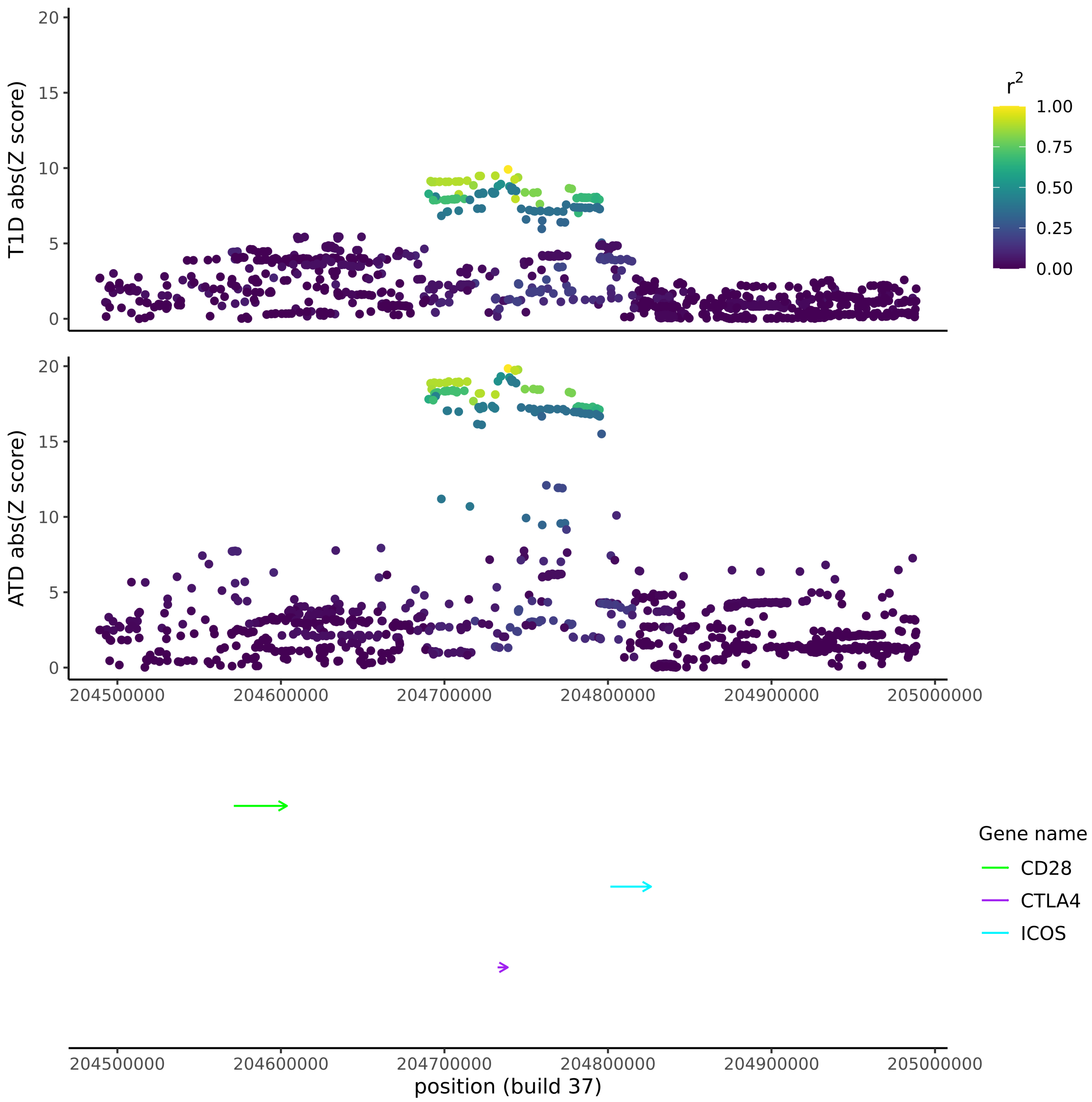
T1D abs(Z score)
ATD abs(Z score)
0
5
10
15
20
r2
1.00
0.75
0.50
0.25
0.00
204500000
204600000
204700000
204800000
204900000
205000000
Gene name
CD28
CTLA4
ICOS
position (build 37)

**Figure S7**

Type 1 diabetes test statistics from FinnGen (data release 6). For each SNP, the GWAS Z-score using strictly defined T1D cases (y-axis) is plotted against its Z-sore using non-strictly defined T1D cases (x-axis). Non-strictly defined cases (code E4_DM1, n=7608) have an ICD-10 code of insulin-dependent diabetes mellitus, while strictly defined cases (code E4_DM1_STRICT, n=3392) comprise the same samples after removal of those with ICD-10 codes of non-insulin-dependent diabetes mellitus. In the main text, we use non-strictly defined cases from FinnGen data release 4 (n=4933).

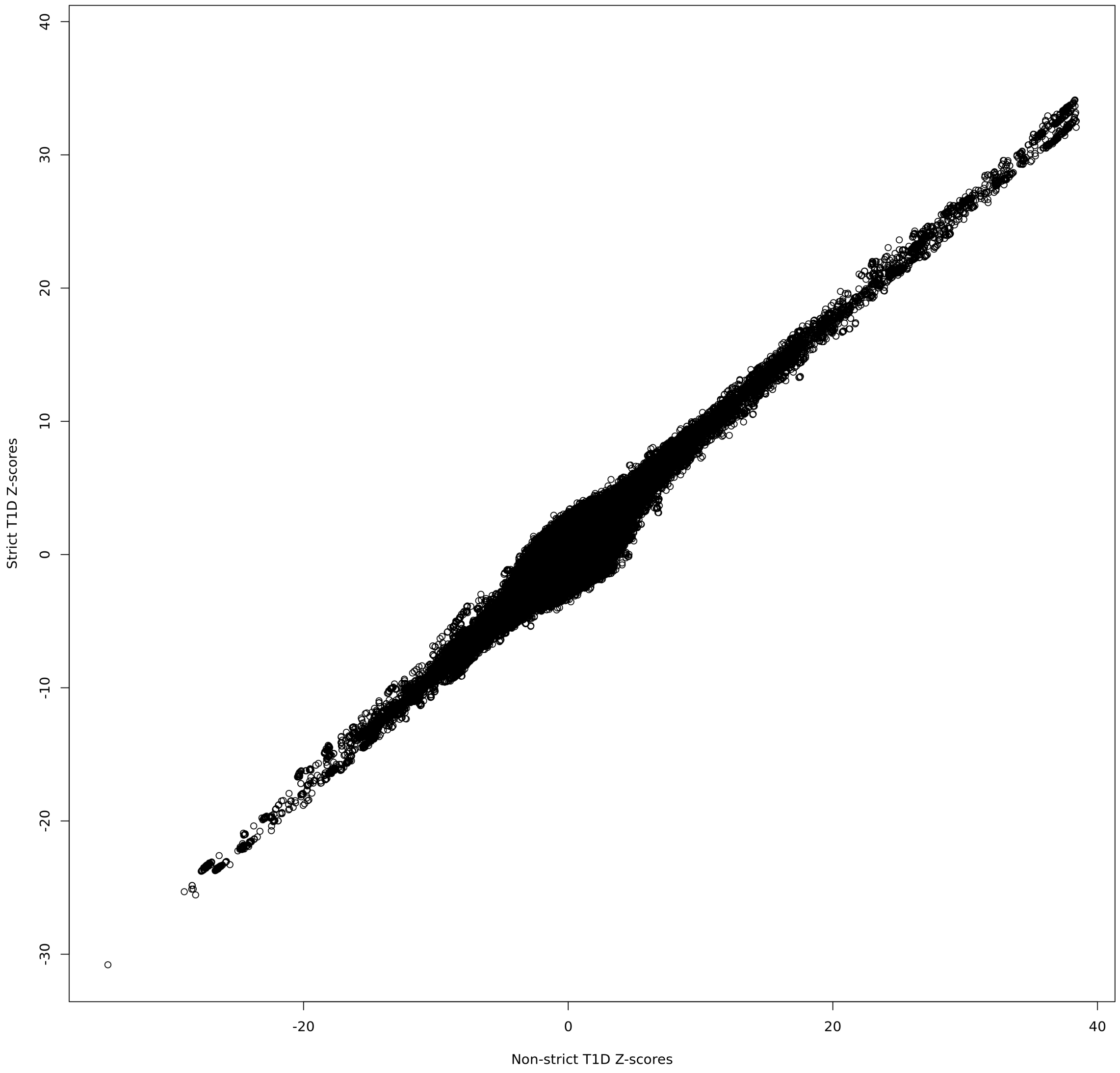
