## Supplementary Information Appendix for "Enhanced genetic analysis of type 1 diabetes by selecting variants on both effect size and significance, and by integration with autoimmune thyroid disease"

### BFDR method for selection of variants based on effect size and significance

#### 1 Overview

The BFDR (Bigger or False Discovery Rate) is a metric for controlling the overall cost of following up variables based on association data from a family of tests, based on a) the probability that the variables are a false positives, and b) the probability that they are true positives, but with effects that are insubstantial compared to other variables from the same family of tests. Our motivation for developing the BFDR came from observing the many SNPs that appear to be statistical outliers on the x-axis of a volcano plot (e.g. main text Figure 2), representing effect size, despite having modest levels of significance, shown on the y-axis. Such variants may be interesting despite not passing standard significance criteria, or relatively uninteresting despite having high significance. Throughout the following we describe the BFDR as it would be applied to GWAS SNP data, but it could equally be applied to other types of multivariate data.

See main text Equations 1-2 for an informal definition of the BFDR. When capitalised, we use $\text{BFDR}_i$ to refer to a tail-area quantity, like a P-value, pertaining to SNPs with results more extreme than SNP $i$:

$$\text{BFDR}_i = \text{FDR}_i + \text{BDR}_i \times (1 - \text{FDR}_i), \tag{1}$$

$$\text{FDR}_i = \Pr(H_0 = 1 \big| \text{fdr} \leqslant \text{fdr}_i, \text{bdr} \leqslant \text{bdr}_i), \tag{2}$$

$$\text{BDR}_i = \Pr(|\beta_{\text{alt}}| \geqslant |\beta| \big| \text{fdr} \leqslant \text{fdr}_i, \text{bdr} \leqslant \text{bdr}_i, H_0 = 0), \tag{3}$$

where $\beta$ is a SNP risk effect, $\beta_{\text{alt}}$ is a second SNP effect drawn from the prior distribution of alternative (alt, i.e. non-null) effect sizes, and $H_0$ is a binary indicator variable taking the value 1 when the null hypothesis is true and 0 when false. Following a general convention, we use the lower-case fdr and bdr to refer to 'local' versions of the FDR and BDR. These are defined as:

$$\text{fdr}_j = \Pr(H_{0(j)} = 1 \big| z_j), \tag{4}$$
$$\text{bdr}_j = \Pr(|\beta_{\text{alt}}| \geqslant |\beta_j| \big| \hat{\beta}_j, \hat{\sigma}_j, H_{0(j)} = 0), \tag{5}$$

where $z_j$, $\hat{\beta}_j$ and $\hat{\sigma}_j$ are the observed Z-score, estimated effect size and estimated standard error for SNP $j$, and $H_{0(j)}$ is the null hypothesis indicator variable for SNP $j$. Our method for estimating BFDR, described in the following sections, is to first estimate $\text{bdr}_j$ and $\text{fdr}_j$ for $j = 1, 2, ...N$, where $N$ is the total number of SNPs analysed, then to empirically integrate them across the tail area to obtain $\text{BDR}_i$ and $\text{FDR}_i$. As the method relies on empirical Bayesian reasoning in which it is assumed that all SNPs share the same prior distribution of risk effects, $N$ should be large, probably at least 1000, in order to obtain good BFDR estimates. If there is reason to believe that there is subset of SNPs, e.g. those in the the HLA region, which is believed not to share a prior effect distribution with the majority, we advise excluding this.

The first term in Equation 1 is the posterior probability that SNPs passing a bivariate threshold defined by SNP $i$ are false positives, while the second term, which can also be rewritten as

$$\mathrm{BDR}_i \times (1 - \mathrm{FDR}_i) = \Pr(|\beta_{\mathrm{alt}}| \geqslant |\beta| \cap H_0 = 0 \big| \mathrm{fdr} \leqslant \mathrm{fdr}_i, \mathrm{bdr} \leqslant \mathrm{bdr}_i), \tag{6}$$

is the probability that they are non-null, but have effect sizes that are exceeded by a randomly chosen non-null SNPs. Using the fact that the events

$$\{H_0 = 1\},\ \{|\beta_{\mathrm{alt}}| \geqslant |\beta| \cap H_0 = 0\} \text{ and } \{|\beta_{\mathrm{alt}}| < |\beta| \cap H_0 = 0\}, \tag{7}$$

have a total probability of 1, the usefulness of the BFDR can be further appreciated by subtracting it from 1:

$$1 - \mathrm{BFDR}_i = \Pr(|\beta_{\mathrm{alt}}| < |\beta| \cap H_0 = 0 \big| \mathrm{fdr} \leqslant \mathrm{fdr}_i, \mathrm{bdr} \leqslant \mathrm{bdr}_i), \tag{8}$$

which is the posterior probability that a SNP in the tail-area is a true positive ($H_0 = 0$) and has an effect size greater than a randomly chosen alt (non-null) SNP ($|\beta_{\mathrm{alt}}| < |\beta|$). We regard this as a good measure of the overall expected utility in following up SNPs in the tail area beyond SNP $i$. Alternatively, the utility for the selected SNPs can be interpreted as equalling zero when they are false positives, and equalling the *proportion* of non-null variants that they exceed in effect size when they are true positives (see main text Equation 2).

When computing the tail-area BFDR in Equation 1, we assume for simplicity that $\mathrm{FDR}_i$ only depends on the SNPs with $\mathrm{fdr} \leqslant \mathrm{fdr}_i$, and $\mathrm{BDR}_i$ only depends on SNPs with $\mathrm{bdr} \leqslant \mathrm{bdr}_i$:

$$\Pr(H_0 = 1 \big| \mathrm{fdr} \geqslant \mathrm{fdr}_i) \equiv \Pr(H_0 = 1 \big| \mathrm{fdr} \leqslant \mathrm{fdr}_i, \mathrm{bdr} \leqslant \mathrm{bdr}_i), \tag{9}$$

$$\Pr(|\beta_{\mathrm{alt}}| \geqslant |\beta| \big| \mathrm{bdr} \geqslant \mathrm{bdr}_i, H_0 = 0) \equiv \Pr(|\beta_{\mathrm{alt}}| \leqslant |\beta| \big| \mathrm{fdr} \leqslant \mathrm{fdr}_i, \mathrm{bdr} \leqslant \mathrm{bdr}_i, H_0 = 0). \tag{10}$$

#### 2 Estimating BFDRs

##### Summary

In essence, the method we present uses polynomial models to fit the distribution of SNP effects as a two-mixture model of null and alternative effects, and also to fit a three-mixture model for a SNPs effect signs, consisting of a 'negative' mixture (true effects below zero), a 'positive' mixture (true effects above zero) and an 'unknown' mixture (effects for which the sign cannot be resolved). We abbreviate these two mixture models as ANE (alt/null effects) and NUPE (negative/unknown/positive effects). To simplify computation, we opt to fit and then fix the parameters of the ANE model before fitting the NUPE model, despite NUPE depending on the parameters of ANE. Fitting the NUPE model involves a novel technique we call prior splitting, described in Section 3, which is the major statistical innovation in our approach to BFDR estimation.

We use the fitted ANE distribution together with Bayes' theorem to find $\text{fdr}_j$ for $j = 1, 2, ...N$. Using the fitted NUPE distribution, Bayes' theorem is applied again to find the posterior probabilities that SNPs $j = 1, 2, ...N$ have effects larger or equal to those for large random selections of SNPs $k_j \in \{1, 2, ...N\}$, so that averaging over $k_j$ estimates $\text{bdr}_j$ for each $j$ in a computationally tractable way, avoiding comparisons between all pairwise SNP combinations. Tail-area estimates are then obtained from $\text{fdr}_j$ and $\text{bdr}_j$ $\{\forall j \in 1, 2, ...N\}$ as overviewed in the previous section. The method, which uses summary statistics rather than individual-level genotype data, is implemented in R and is available in the package `priorsplitteR` (https://github.com/djmcrouch/priorsplitteR).

The ANE and NUPE models are estimated using polynomial likelihood models for the observed SNP effect estimate data, so our method falls within the f-modelling category of empirical Bayesian approaches, in contrast with g-modelling in which the prior is modelled explicitly. Although a discussion of the merits and drawbacks of the two approaches lies outside the scope of this work, we note that there is a strong intuitive appeal in fitting a model which describes data that can be directly observed, so that model and data can be easily compared, e.g. visually, to assess the quality of the fit. It is often the case that a variety of different prior distribution models can fit a dataset almost as well as each other [Efron and Hastie, 2016], so we see it as an advantage not to rely too heavily on any particular choice.

The stages of BFDR estimation are:

1) Estimate the parameters of ANE.

2) Simulate effect estimates from the estimated alt distribution using the `distr` R package. Take the difference between the two sets to give a simulated set of effect estimates under the alt hypothesis. Estimate NUPE parameters using the simulated effect estimate dataset.

3) Simulate two sets of effect estimates, one from the NUPE mixture corresponding to SNPs with positive effects, and one from the mixture corresponding to negative effects. Take the difference between each of these and the alt simulation from Stage 2, and fit models of to both sets of differences using the NUPE model ('NUPE-diff') as a means of estimating $\Pr(\beta_k > \beta_j | \beta_j > 0, \hat{\beta}_j, \hat{\sigma}_j, \hat{\beta}_k, \hat{\sigma}_k, H_{0(j)} = 0, H_{0(k)} = 1)$ and $\Pr(\beta_k < \beta_j | \beta_j < 0, \hat{\beta}_j, \hat{\sigma}_j, \hat{\beta}_k, \hat{\sigma}_k, H_{0(j)} = 0, H_{0(k)} = 1)$.

4) Compute $\text{fdr}_j$ for each SNP $j = 1, 2, ...N$ using the fitted ANE model

5) Compute $\text{bdr}_j$ for each SNP $j = 1, 2, ...N$ using the fitted NUPE-diff models, by averaging $\Pr(\beta_k > \beta_j | \beta_j > 0, \hat{\beta}_j, \hat{\sigma}_j, \hat{\beta}_k, \hat{\sigma}_k, H_{0(j)} = 0, H_{0(k)} = 1)$ and $\Pr(\beta_k < \beta_j | \beta_j < 0, \hat{\beta}_j, \hat{\sigma}_j, \hat{\beta}_k, \hat{\sigma}_k, H_{0(j)} = 0, H_{0(k)} = 1)$ and $\Pr(\beta_k < \beta_j | \beta_j < 0, \hat{\beta}_j, \hat{\sigma}_j, \hat{\beta}_k, \hat{\sigma}_k, H_{0(j)} = 0, H_{0(k)} = 1)$ over a set of alt-distribution SNPs indexed by $k$.

6) Estimate tail-area $\text{FDR}_i$ and $\text{BDR}_i$ using estimates of $\text{fdr}_j$ and $\text{bdr}_j$, and the approach outlined in the previous section. Calculate $\text{BFDR}_i$ from $\text{FDR}_i$ and $\text{BDR}_i$ using Equation 1, for each SNP $i = 1, 2, ...N$.

#### Stage 1: Estimate the parameters of ANE

For the present purpose, we assume that optimal ANE model fitting can be achieved using z-scores ($z_j = \hat{\beta}_j / \hat{\sigma}_j$ for SNP $j$), without considering the effect sizes and standard errors separately, i.e. that SNPs with different standard errors have similar z-score distributions[1]. Simulation experiments demonstrated that our method is robust to violations of this assumption (see main text and online methods).

We estimate distributions using an implementation of Lindsey's Poisson regression method ([Lindsey, 1974a, Lindsey, 1974b]). As if producing a histogram, imagine discretising the z-scores into $B$ bins $b = 1, 2...B$, with $B \equiv 100$ by default, treating the number of SNPs falling into each bin as a random Poisson variable. Thus, the likelihood for the number of SNPs

[1] Before fitting the ANE model in Stage 1, we remove outlying SNPs defined by default as those with $|z_j| > 15$, as these can cause numerical model-fitting problems. The remaining SNPs are sorted into a specified number (default 100) of z-score bins each of equal width, spanning the range of z-scores.

(the count, $c_b$) falling into bin $b$ would be

$$\frac{(N\lambda(z_b)^{c_b})e^{-N\lambda(z_b)}}{!c_b}, \tag{11}$$

where $\lambda(z_b)$ approximates the probability density function of $z$ within the region spanned by bin $b$ as a function of its central point $z_b$, and $N\lambda(z_b)$ is the Poisson parameter for bin $b$. As described in Efron and Hastie ([Efron and Hastie, 2016]), one could use polynomial model for $\lambda(z_b)$:

$$\lambda(z_b) = e^{\sum_{n=0}^{D} \gamma_n z_b^n}, \tag{12}$$

where $D$ is the degree of the polynomial[2]. Extending Lindsey's method, we use two $\lambda_b$ variables for each bin, representing the alternative and null mixture components:

$$\lambda_1(z_b) = e^{\sum_{n=0}^{D} \gamma_n z_b^n}, \tag{13}$$

$$\lambda_0(z_b) = w\phi(z_b), \tag{14}$$

where $\phi(\cdot)$ is the standard normal distribution function, $\gamma_n$ is a model parameter and $w$ the width of the bins[3]. We set $D \equiv 6$ by default, providing a balance between model flexibility and not overfitting the data. The likelihood for the counts in bin $b$ is

$$L_b(c_b) = \pi\frac{(N\lambda_0(z_b))^{c_b}e^{-N\lambda_0(z_b)}}{!c_b} + (1-\pi)\frac{(N\lambda_1(z_b))^{c_b}e^{-N\lambda_1(z_b)}}{!c_b}, \tag{15}$$

where $\pi$ is the mixing proportion, also the prior probability of a SNP belonging to the null distribution of zero effect. To obtain maximum-likelihood estimates of the parameters $\pi$ and $\boldsymbol{\gamma} = [\gamma_1, \gamma_2, ...\gamma_D]$, we maximise the overall log-likelihood $\sum_{b=1}^{B} \ln L_b(c_b)$. Rather than maximising directly, which is challenging for mixture models, we employ an EM-algorithm [Dempster et al., 1977], treating the expected proportion of counts belonging to the null distribution within each bin, conditioned on the current parameter choices $\theta = \{\pi, \boldsymbol{\gamma}\}$, as an unknown latent variable:

$$m_b = \frac{\lambda_0(z_b)\pi}{\lambda_0(z_b)\pi + \lambda_1(z_b)(1-\pi)}, \tag{16}$$

[2]Scaling is applied to $z_b^n$ so that SD=1 across $b$ for each $n$, equivalent to scaling each $\gamma_n$, to simplify numerical optimisation.

[3]In practice, we use the cumulative normal distribution to improve the approximation $w\phi(z_b)$ (and for the equivalent null distribution in the NUPE model), by computing the probability mass lying within the bin, but this is not essential.

where the expected proportion belonging to the alt distribution is $1 - m$. Using this equation to compute $m_b$ based on $\theta$ (the E-step), we maximise (in the M-step) $Q(\theta') = \sum_{b=1}^{B}(f_{b0}(c_b) + f_{b1}(c_b))$, where

$$f_{b1}(c_b) = c_b(1 - m_b)(\ln(1 - \pi') + \ln\lambda_1'(z_b)) - (1 - \pi')\lambda_1'(z_b)N, \tag{17}$$

$$f_{b0}(c_b) = c_b m_b \ln\pi' - \pi'\lambda_0(z_b)N, \tag{18}$$

with respect to $\theta' = \{\pi', \gamma'\}$ and $\lambda_1'(z_b)$ is Equation 13 parameterised by $\gamma'$ [4]. Maximisation is performed with the Broyden-Fletcher-Goldfarb-Shanno (BFGS) algorithm [Broyden, 1970, Fletcher, 1970, Goldfarb, 1970, Shanno, 1970] implemented in the `constrOptim` R function, using the first partial derivatives w.r.t. each parameter, which we derived analytical expressions for. The BFGS algorithm requires an initialisation point (described below), from which it explores the parameter space using the $Q(\theta')$ values and derivatives until a convergence threshold is exceeded, or until a maximum number of iterations is reached (default $10^8$). The $\pi'$ parameter is constrained by `constrOptim` to be within $[0, 1]$[5].

Equations 17 and 18 are derived from the expected value of the log-likelihood $\sum_{b=1}^{B}(h_{b0}(c_b) + h_{b1}(c_b))$, where

$$h_{b1} = c_b(1 - M_b)(\ln(1 - \pi') + \ln\lambda_1'(z_b) + \ln N) - (1 - \pi')\lambda_1'(z_b)N - (c_b(1 - M_b))!, \tag{19}$$

$$h_{b0} = c_b M_b(\ln\pi' + \ln\lambda_0(z_b) + \ln N) - \pi'\lambda_0(z_b)N - (c_b M_b)!, \tag{20}$$

and $M_b$ is a random variable, conditioned on current parameter values $\pi$ and $\lambda_1(z_b)$, giving the proportion of counts in bin $b$ belonging to the null distribution. The expected value of $M_b$ is $m_b$. In Equations 17 and 18 we ignore terms for which the equivalents in Equations 19 and 20 contain no parameters, as they remain constant under maximisation. This is particularly helpful as there are no closed-form expressions for expectations of the final terms $(c_b(1 - M_b))!$ and $(c_b M_b)!$.

The M- and E-steps are iterated until convergence. In order to increase the log-likelihood using a generalised EM-algorithm, it is sufficient to find values of $\pi'$ and $\lambda_1'(z_b)$ that increase (i.e. improve) the expected value of $Q(\theta')$, relative to $Q(\theta)$ [Hastie et al., 2009]. We therefore

[4] $\lambda_0(z_b)$ contains no parameters.

[5] We also aid fitting of ANE by constraining $\lambda_1(z_b) < \lambda_0(z_b)$ at $z_b = 0$, so as to avoid impossible distributions, by constraining $\gamma_0 < \ln w\phi(0)$.

iterate until $Q(\theta') - Q(\theta)$ becomes stable and close to zero, with a default of less than $10^{-6}$ for five consecutive iterations[6]. The first M-step is initialised at an arbitrary point in the parameter space, but subsequent M-steps are initialised at the maximised parameter values obtained by the previous M-step ($\theta' \equiv \theta$) as, being generally close to the optimum, we found this provided the most reliable convergence.

Estimation of ANE using our type 1 diabetes (T1D) and autoimmune thyroid disease (ATD) additive GWAS meta-analysis data produce the fitted models shown in Figure A1. Estimates of $\pi$ were 0.80 for ATD and 0.87 for T1D, indicating that approximately 80%-90% of variants are estimated to have no effect on disease. When performing T1D GWAS analysis within the subset of significantly ATD-associated variants, the estimate for $\pi$ was 0.16, showing that a large majority of ATD-associated SNPs also have risk effects on T1D. It is important to recognise that most SNP effects estimated to be non-null are likely to be non-causal LD-tagging associations, but the BFDR should be used to establish interesting correlative effects, before using stepwise regression and fine-mapping to identify which of these are likely to be causal.

##### Stage 2: Simulate alt effect estimates and estimate the NUPE parameters

Once the ANE model has been estimated, we fix its parameters at the maximum-likelihood estimates $\hat{\pi}$ and $\hat{\lambda}_1(z_b)$ (hat symbols throughout indicate maximum-likelihood estimates of the corresponding parameter) and proceed to estimate the parameters of NUPE[7]. The NUPE model decomposes the distribution of alt SNP estimates into separate mixtures for those with true effects above zero, those with true effects below zero, and those for which the true effect cannot be distinguished from zero due to insufficient power. As NUPE is only concerned with non-null SNPs, we draw a large sample of z-scores, $z_s$ for $s \equiv 1, 2, ...S$, from the fitted alt distribution defined by $\lambda_1(z_b)$ using the R package `distr`[8]. Sufficiently many samples should be drawn to avoid introducing significant levels of noise, so by default we set $S = 5 \times 10^6$.

The purpose of the NUPE model is to provide a basis for estimating the probability that a true effect $\beta_s$ is positive, negative or has an unknown sign, given its corresponding z-score:

[6] We also impose a default maximum of $10^5$ iterations before accepting the parameter estimates.

[7] We take this approach for computational tractability, though in reality the best NUPE estimates will probably depend on the parameters of ANE to some extent.

[8] As $\hat{\lambda}_1$ is the estimated probability mass within the bin, with width $w$, for sampling we use the density function $\hat{\lambda}_1(z)/w$.

**Figure A1:** Fitted ANE model for a) type 1 diabetes (T1D) and b) autoimmune thyroiditis (ATD) GWAS meta-analyses data (under additive models). Red, green and blue curves show the marginal mixture distribution, the alt mixture and null mixture respectively.

(a) T1D (b) ATD

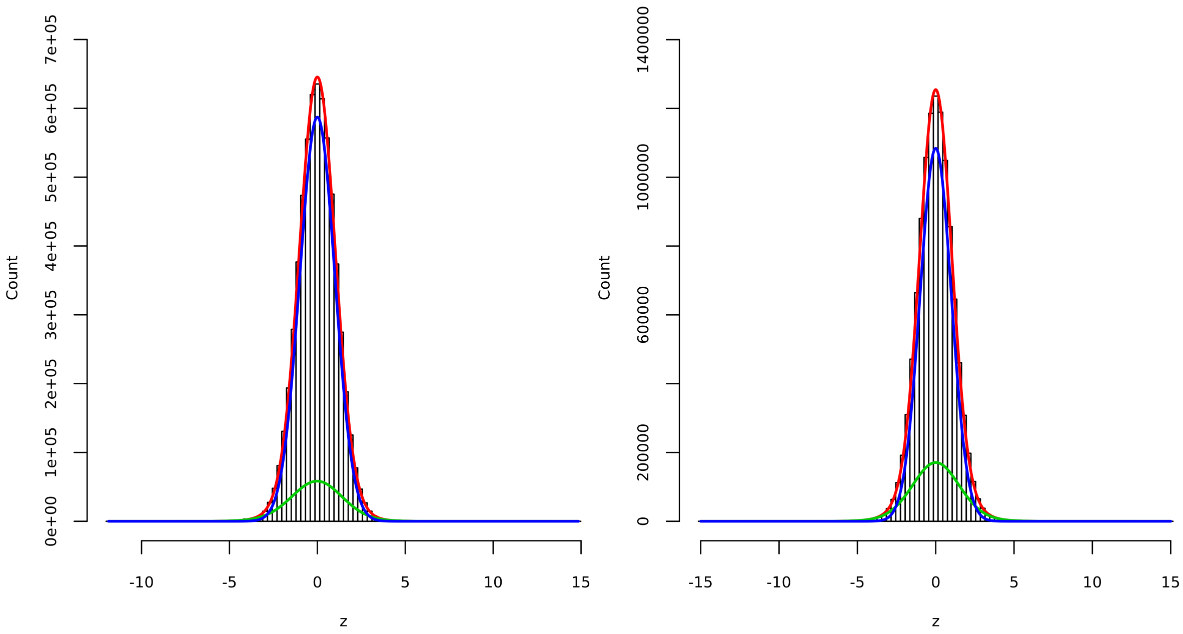

$$z_s = \frac{\beta_s}{\hat{\sigma}_s} + \epsilon_s, \tag{21}$$

where $\epsilon_s$ is a normal random variable with mean zero and unit variance. Though $\beta_s/\hat{\sigma}_s$ has an unknown distribution, we assume that it is drawn from the same prior distribution $g(\cdot)$ for all $s$. Rather than attempting to estimate $g(\cdot)$ explicitly, we use a novel technique to split the distribution of $z_s$ into a mixture of three distributions with non-overlapping priors, corresponding to the positive, negative and unknown-sign effects. Following Lindsey's method as applied in Stage 1[9], the distributions of z-scores within each mixture respectively are:

$$\Lambda_{(+)}(z_b) = W\phi(z_b)e^{z_b \sum_{n=0}^{D}\sum_{p=0}^{D} \frac{\Gamma_n \Gamma_p z_b^n z_b^p}{n+p+1} + a}, \tag{22}$$

$$\Lambda_{(-)}(z_b) = W\phi(z_b)e^{-z_b \sum_{n=0}^{D}\sum_{p=0}^{D} \frac{\Gamma_n^* \Gamma_p^* z_b^n z_b^p}{n+p+1} + a^*}, \tag{23}$$

$$\Lambda_{(u)}(z_b) = W\phi(z_b), \tag{24}$$

[9]By default we use the same choices of $D \equiv 6$ and the same specification of bins as for ANE.

where $z_b$ is the central point of bin $b$, $W$ is the width of the bins, and $\Gamma_n$, $\Gamma_p$, $a$, $\Gamma^*_n$, $\Gamma^*_p$ and $a^*$ are model parameters, with $a$ and $a^*$ acting as intercept terms. In this model, $\Lambda_{(u)}(z_b)$ is equivalent to a null distribution, but we avoid describing it in this way as the NUPE model is fitted using simulated alt (non-null) SNPs, so it is best interpreted as the distribution of alt SNPs that have effects too small to be able to assign a sign to. Summation expressions within the exponents are similar to squared polynomials[10], and as such have a general flexibility that becomes greater as $D$ is increased, similar to the polynomial used for ANE in Equation 13. However, derivatives of the ratios $\Lambda_{(+)}(z_b)/\Lambda_{(u)}(z_b)$ and $\Lambda_{(-)}(z_b)/\Lambda_{(u)}(z_b)$ are always non-negative and non-positive respectively. If $g_{(+)}(\cdot)$ and $g_{(-)}(\cdot)$ are the priors corresponding to distributions $\Lambda_{(+)}(z_b)$ and $\Lambda_{(-)}(z_b)$, we show in Section 3 that these constraints are equivalent to $g_{(-)}(x) = 0$ for $x \geqslant 0$ and $g_{(+)}(x) = 0$ for $x \leqslant 0$.

Estimation is performed using an EM-algorithm similar to Stage 1, where the expected proportions in each bin deriving from each mixture, conditioned on the current parameter estimates $\theta = \{\rho_{(+)}, \rho_{(-)}, \Lambda_{(+)}(z_b), \Lambda_{(-)}(z_b)\}$ are computed in the E-step as:

$$m_{b(+)} = \frac{\Lambda_{(+)}(z_b)\rho_{(+)}}{\Lambda_{(+)}(z_b)\rho_{(+)} + \Lambda_{(-)}(z_b)\rho_{(-)} + \Lambda_{(u)}(z_b)(1 - \rho_{(+)} - \rho_{(-)})}, \tag{25}$$

$$m_{b(-)} = \frac{\Lambda_{(-)}(z_b)\rho_{(-)}}{\Lambda_{(+)}(z_b)\rho_{(+)} + \Lambda_{(-)}(z_b)\rho_{(-)} + \Lambda_{(u)}(z_b)(1 - \rho_{(+)} - \rho_{(-)})}, \tag{26}$$

$$m_{b(u)} = 1 - m_{b(+)} - m_{b(-)}, \tag{27}$$

where $\rho_{(+)}$ and $\rho_{(-)}$ are the mixing proportions for $\Lambda_{(+)}(z_b)$ and $\Lambda_{(-)}(z_b)$. In the M-step, similar to ANE, we maximise $Q(\theta') = \sum_{b=1}^{B}(f_{b0}(c_b) + f_{b(+)}(c_b) + f_{b(-)}(c_b))$, with respect to new parameters $\theta' = \{\rho'_{(+)}, \rho'_{(-)}, \mathbf{\Gamma} = \{\Gamma_1, \Gamma_2, ...\Gamma_D\}, \mathbf{\Gamma^*} = \{\Gamma^*_1, \Gamma^*_2, ...\Gamma^*_D\}, a, a^*\}$ where

$$f_{b(+)}(c_b) = c_b m_{b(+)}\big(\ln\rho'_{(+)} + \ln\Lambda'_{(+)}(z_b)\big) - \rho'_{(+)}\Lambda'_{(+)}(z_b)S, \tag{28}$$

$$f_{b(-)}(c_b) = c_b m_{b(-)}\big(\ln\rho'_{(-)} + \ln\Lambda'_{(-)}(z_b)\big) - \rho'_{(-)}\Lambda'_{(-)}(z_b)S, \tag{29}$$

$$f_{b(u)}(c_b) = c_b m_{b(u)}\ln\big(1 - \rho'_{(+)} - \rho'_{(-)}\big) - \big(1 - \rho'_{(+)} - \rho'_{(-)}\big)\Lambda_{(u)}(z_b)S, \tag{30}$$

and where $c_b$ now represents the number of $z_s$ falling into each bin $b$, with $\sum_{b=1}^{B} c_b = S$. As for ANE estimation, maximisation of $Q(\theta')$ is made easier using functions we wrote to compute its first partial derivatives, and `constrOptim` is used to search the parameter space for the optimum. Each M-step is initialised at $\theta' \equiv \theta$, as in ANE estimation, and we iterate

[10]These summations are squared polynomials after dividing each term by $n + p + 1$.

E- and M-steps until the same convergence criteria are met. Both $\rho'_{(+)}$ and $\rho'_{(-)}$ and their sum are constrained to be within $[0, 1]$[11]. Equations 28-30 are derived from the expected log-likelihoods for each mixture distribution, ignoring terms containing no parameters, as in Stage 1.

Estimates of NUPE are shown in Figure A2. Effect signs were harder to infer for T1D, for which we had approximately half the number of cases as ATD, indicated by the larger mixing proportion of the blue unknown-sign curve.

**Figure A2:** Fitted NUPE model for a) type 1 diabetes (T1D) and b) autoimmune thyroiditis (ATD) GWAS meta-analyses data (under additive models). Green and turquoise curves are the fitted distributions for negative and positive true effects respectively. SNPs with ambiguous effect signs, which there is insufficient power to distinguish from zero, are distributed according to the blue curve, and the fitted marginal mixture distribution is red.

(a) T1D

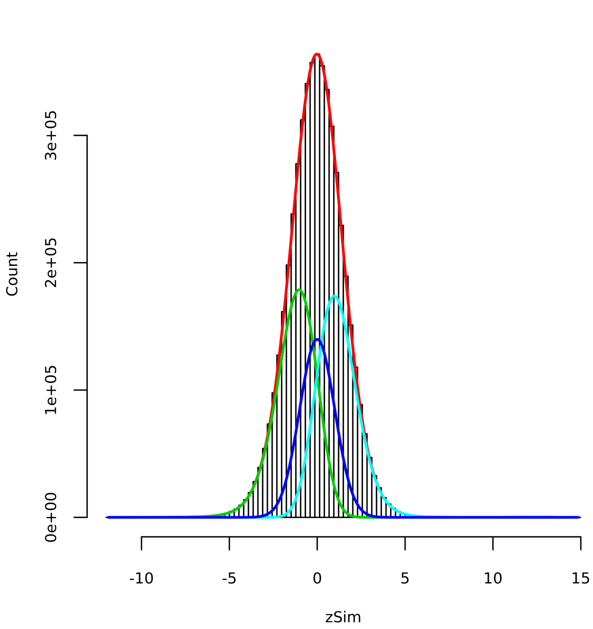

(b) ATD

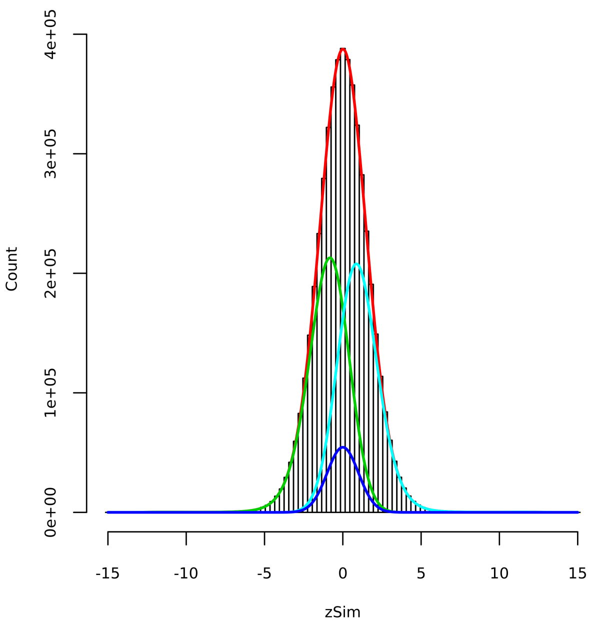

[11] Similar to ANE, we constrain $a$ and $a'$ to be negative with `constrOptim`, thereby avoiding impossible distributions by ensuring $\Lambda_{(+)}(z_b)$ and $\Lambda_{(-)}(z_b)$ are less than $\Lambda_{(u)}(z_b)$ at $z_b = 0$. To prevent potential maximisation problems arising from asymmetry in the model, we also constrain $\Gamma_0$ and $\Gamma_0^* > 0$.

##### Stage 3: Estimate the parameters of NUPE-diff models

As a route towards modelling $\Pr(\beta_k > \beta_j | \beta_j > 0, \hat{\beta}_j, \hat{\sigma}_j, \hat{\beta}_k, \hat{\sigma}_k, H_{0(j)} = 0, H_{0(k)} = 1)$ and $\Pr(\beta_k < \beta_j | \beta_j < 0, \hat{\beta}_j, \hat{\sigma}_j, \hat{\beta}_k, \hat{\sigma}_k, H_{0(j)} = 0, H_{0(k)} = 1)$, we use simulated z-scores to define two sets of scaled effect-estimate differences:

$$\delta_{s(+)} = \frac{z_{s(+)}\hat{\sigma}_{s(+)} - z_s\hat{\sigma}_s}{\sqrt{\hat{\sigma}^2_{s(+)} + \hat{\sigma}^2_s}}, \tag{31}$$

$$\delta_{s(-)} = \frac{z_{s(-)}\hat{\sigma}_{s(-)} - z_s\hat{\sigma}_s}{\sqrt{\hat{\sigma}^2_{s(-)} + \hat{\sigma}^2_s}}, \tag{32}$$

where $z_{s(+)}$ and $z_{s(-)}$ are simulated in the same way as $z_s$ but from the upper and lower sign distributions (taking $S$ samples), rather than from the alt distribution, and the SEs $\hat{\sigma}_{s(+)}$, $\hat{\sigma}_{s(-)}$ and $\hat{\sigma}_s$ are sampled from the empirical distribution of SEs according to their probabilities of belonging to the positive-sign, negative-sign or alt distributions respectively[12]. As $\delta_{s(+)}$ and $\delta_{s(-)}$ are scaled differences between z-scores, we model them as

$$\delta_{s(+)} = \Delta_{s(+)} + \epsilon_{s(+)}, \tag{33}$$

$$\delta_{s(-)} = \Delta_{s(-)} + \epsilon_{s(-)}, \tag{34}$$

where $\epsilon_{s(+)}$ and $\epsilon_{s(-)}$ have normal distributions with mean zero and unit variance, and $\Delta_{s(+)}$ and $\Delta_{s(-)}$ have unknown prior distributions similar to $g(\cdot)$ from the previous subsection. Applying the NUPE estimation method again[13] to the scaled differences $\delta_{s(+)}$ (NUPE-diff) provides estimates of three mixture distributions for $\delta_{s(+)}$ conditioned on $\Delta_{s(+)} > 0$, $\Delta_{s(+)} < 0$ or $\Delta_{s(+)}$ having unknown sign. All three of these mixtures are conditioned on $z_{s(+)}$ having a positive true effect, so are equivalent to the likelihoods:

[12]For $\hat{\sigma}_s$, $S$ random samples are drawn from a multinomial distribution with $N$ categories, where the probability of drawing each SNP $i = 1, 2, ...N$ is $\frac{1}{N}\frac{\hat{\lambda}_1(z_i)(1-\hat{\pi})}{\hat{\lambda}(z_i)}$, where hat symbols indicate estimated ANE parameters and $N^{-1}$ is the empirical marginal distribution, which estimates an empirical version of the marginal distribution $\lambda(z_i)$. For $\hat{\sigma}_{s(+)}$ and $\hat{\sigma}_{s(-)}$ this is multiplied by equivalent probabilities obtained using the estimates of the relevant distributions from NUPE, to reflect that the sign distributions are both conditional on the alt distribution.

[13]Bins are reset for analysis of $\delta_{s(+)}$ and $\delta_{s(-)}$, using $B \equiv 100$ by default, and we discard simulated differences with absolute values greater than $22.5 = 15 \times 1.5$, where 15 is the default threshold for discarding z-scores.

$$p(\delta_{jk}^{(\text{obs})}|\beta_k > \beta_j, \beta_j > 0, H_{0(j)} = 0) \equiv \hat{\Lambda}_{(-)}(\delta_{jk}^{(\text{obs})})|\beta_j > 0, \tag{35}$$
$$p(\delta_{jk}^{(\text{obs})}|\beta_k < \beta_j, \beta_j > 0, H_{0(j)} = 0), \equiv \hat{\Lambda}_{(+)}(\delta_{jk}^{(\text{obs})})|\beta_j > 0, \tag{36}$$
$$p(\delta_{jk}^{(\text{obs})}|\beta_k = \beta_j, \beta_j > 0, H_{0(j)} = 0), \equiv \hat{\Lambda}_{(u)}(\delta_{jk}^{(\text{obs})})|\beta_j > 0, \tag{37}$$

where $|\beta > 0$ indicates that estimates were obtained using the $\delta_{s(+)}$ simulated data set, and $\delta_{jk}^{(\text{obs})}$ is an observed scaled effect estimate difference:

$$\delta_{jk}^{(\text{obs})} = \frac{\hat{\beta}_j - \hat{\beta}_k}{\sqrt{\hat{\sigma}_j^2 + \hat{\sigma}_k^2}}. \tag{38}$$

The marginal likelihood conditioned on $\beta > 0$ is

$$p(\delta_{jk}^{(\text{obs})}|\beta_j > 0, H_{0(j)} = 0) \equiv \hat{\Lambda}(\delta_{jk}^{(\text{obs})})|\beta_j > 0 = (\hat{\Lambda}_{(-)}(\delta_{jk}^{(\text{obs})})|\beta_j > 0)(\hat{\rho}_{(-)}|\beta_j > 0) \tag{39}$$
$$+ (\hat{\Lambda}_{(+)}(\delta_{jk}^{(\text{obs})})|\beta_j > 0)(\hat{\rho}_{(+)}|\beta_j > 0) \tag{40}$$
$$+ (\hat{\Lambda}_{(u)}(\delta_{jk}^{(\text{obs})})|\beta_j > 0)(1 - \hat{\rho}_{(-)}|\beta_j > 0 - \hat{\rho}_{(+)}|\beta_j > 0). \tag{41}$$

Similarly, applying NUPE-diff a second time to $\delta_{s(-)}$ yields estimates of

$$p(\delta_{jk}^{(\text{obs})}|\beta_k > \beta_j, \beta_j < 0, H_{0(j)} = 0) \equiv \hat{\Lambda}_{(-)}(\delta_{jk}^{(\text{obs})})|\beta_j < 0, \tag{42}$$
$$p(\delta_{jk}^{(\text{obs})}|\beta_k < \beta_j, \beta_j < 0, H_{0(j)} = 0), \equiv \hat{\Lambda}_{(+)}(\delta_{jk}^{(\text{obs})})|\beta_j < 0, \tag{43}$$
$$p(\delta_{jk}^{(\text{obs})}|\beta_k = \beta_j, \beta_j < 0, H_{0(j)} = 0), \equiv \hat{\Lambda}_{(u)}(\delta_{jk}^{(\text{obs})})|\beta_j < 0, \tag{44}$$

with marginal $\hat{\Lambda}(\delta_{jk}^{(\text{obs})})|\beta_j < 0$. In order to quantify the probability that $\beta_k$ is larger than $\beta_j$ but with opposite sign, we re-run both NUPE-diff procedures using a set of reversed alt-distribution effect estimates:

$$\frac{z_{s(+)}\hat{\sigma}_{s(+)} - (-z_s\hat{\sigma}_s)}{\sqrt{\hat{\sigma}_{s(+)}^2 + \hat{\sigma}_s^2}}, \tag{45}$$
$$\frac{z_{s(-)}\hat{\sigma}_{s(-)} - (-z_s\hat{\sigma}_s)}{\sqrt{\hat{\sigma}_{s(-)}^2 + \hat{\sigma}_s^2}}, \tag{46}$$

producing an equivalent set of estimated likelihoods denoted by tilde symbols[14], e.g. $\tilde{\Lambda}(\delta_{jk}^{(\text{rev})})|\beta_j > 0$, in order to apply to reverse-orientation versions of $\delta_{jk}^{(\text{obs})}$:

[14]If the alt distribution is roughly symmetrical the original estimates, denoted by hat symbols, could be used in their place if desired.

$$\delta_{jk}^{(\text{rev})} = \frac{\hat{\beta}_j - (-\hat{\beta}_k)}{\sqrt{\hat{\sigma}_j^2 + \hat{\sigma}_k^2}}. \tag{47}$$

Bayes' theorem will be applied to these 12 likelihoods to obtain estimates of $\text{bdr}_j$ in Stage 5. Estimated NUPE-diff models are displayed in Figure A3.

#### Stage 4: Compute fdrs

With a view to estimating FDRs (Equation 9), we first compute the estimate of $\text{fdr}_j$ (see Equation 4) for each SNP[15] $j$ using Bayes' theorem and the ANE maximum-likelihood parameter estimates:

$$\widehat{\text{fdr}}_j = \frac{\hat{\lambda}_0(z_j)\hat{\pi}}{\hat{\lambda}_0(z_j)\hat{\pi} + \hat{\lambda}_1(z_j)(1-\hat{\pi})}. \tag{48}$$

#### Stage 5: Compute bdrs

To estimate $\text{bdr}_j$ (Equation 5), a random selection of $S$ SNPs is first drawn as if sampling from the empirical alt-distribution[16], before computing a set of effect differences[17] $\delta_{jk}^{(\text{obs})}$ and reverse orientation differences $\delta_{jk}^{(\text{rev})}$ (Equations 38 and 47) where $k$ indexes, for computational reasons, a smaller random subset $t_j \in \{1, 2, 3...S\}$ of the $S$ selected SNPs, with size $T$. For each $k$ we use the likelihoods in Expressions 35-44 to compute estimated posteriors

[15] Z-scores lying outside the range of those used for fitting the distributions (due to removal of outliers) are set to the value of either the most negative or most positive z-score used, depending on their sign, as fits are likely to be unreliable outside of this range.

[16] Using the same method as for randomly selecting $\hat{\sigma}_s$ in Stage 3.

[17] Suppressing outliers as in Stage 4.

$$\widehat{\Pr}(\beta_k \geqslant \beta_j | \delta_{jk}^{(\text{obs})}, \beta_j > 0, H_{0(j)} = 0) = \frac{(\hat{\Lambda}_{(-)}(\delta_{jk}^{(\text{obs})})|\beta_j > 0)(\hat{\rho}_{(+)}|\beta_j > 0)}{\hat{\Lambda}(\delta_{jk}^{(\text{obs})})|\beta_j > 0} + \frac{(\hat{\Lambda}_{(u)}(\delta_{jk}^{(\text{obs})})|\beta_j > 0)(1 - (\hat{\rho}_{(-)}|\beta_j > 0) - (\hat{\rho}_{(+)}|\beta_j > 0))}{\hat{\Lambda}(\delta_{jk}^{(\text{obs})})|\beta_j > 0},$$

$$\widetilde{\Pr}((-\beta_k) > \beta_j | \delta_{jk}^{(\text{rev})}, \beta_j > 0, H_{0(j)} = 0) = \frac{(\tilde{\Lambda}_{(-)}(\delta_{jk}^{(\text{rev})})|\beta_j > 0)(\tilde{\rho}_{(+)}|\beta_j > 0)}{\tilde{\Lambda}(\delta_{jk}^{(\text{rev})})|\beta_j > 0},$$

$$\widehat{\Pr}(\beta_k \leqslant \beta_j | \delta_{jk}^{(\text{obs})}, \beta_j < 0, H_{0(j)} = 0) = \frac{(\hat{\Lambda}_{(+)}(\delta_{jk}^{(\text{obs})})|\beta_j < 0)(\hat{\rho}_{(+)}|\beta_j < 0)}{\hat{\Lambda}(\delta_{jk}^{(\text{obs})})|\beta_j < 0}, + \frac{(\hat{\Lambda}_{(u)}(\delta_{jk}^{(\text{obs})})|\beta_j < 0)(1 - (\hat{\rho}_{(-)}|\beta_j < 0) - (\hat{\rho}_{(+)}|\beta_j < 0))}{\hat{\Lambda}(\delta_{jk}^{(\text{obs})})|\beta_j < 0},$$

$$\widetilde{\Pr}((-\beta_k) < \beta_j | \delta_{jk}^{(\text{rev})}, \beta_j < 0, H_{0(j)} = 0) = \frac{(\tilde{\Lambda}_{(+)}(\delta_{jk}^{(\text{rev})})|\beta_j < 0)(\tilde{\rho}_{(+)}|\beta_j < 0)}{\tilde{\Lambda}(\delta_{jk}^{(\text{rev})})|\beta_j < 0}. \tag{49}$$

Averaging each over the selection indexed by $k$ gives one-tailed bdr estimates conditioned on either $\beta_j > 0$ or $\beta_j < 0$:

$$\widehat{\text{bdr}}_j|\beta_j > 0 = \frac{1}{T}\sum_{k \in t_j} \widehat{\Pr}(\beta_k \geqslant \beta_j | \delta_{jk}^{(\text{obs})}, \beta_j > 0, H_{0(j)} = 0), \tag{50}$$

$$\widetilde{\text{bdr}}_j|\beta_j > 0 = \frac{1}{T}\sum_{k \in t_j} \widetilde{\Pr}((-\beta_k) > \beta_j | \delta_{jk}^{(\text{rev})}, \beta_j > 0, H_{0(j)} = 0), \tag{51}$$

$$\widehat{\text{bdr}}_j|\beta_j < 0 = \frac{1}{T}\sum_{k \in t_j} \widehat{\Pr}(\beta_k \leqslant \beta_j | \delta_{jk}^{(\text{obs})}, \beta_j < 0, H_{0(j)} = 0), \tag{52}$$

$$\widetilde{\text{bdr}}_j|\beta_j < 0 = \frac{1}{T}\sum_{k \in t_j} \widetilde{\Pr}((-\beta_k) < \beta_j | \delta_{jk}^{(\text{rev})}, \beta_j < 0, H_{0(j)} = 0). \tag{53}$$

This stage may be parallelised across several thousand SNPs simultaneously[18]. By default we take a relatively small sample of SNPs ($T \equiv 1000$), to save computation time. However, low conditional bdrs (default below 0.05) in Equations 50-53 are resampled with more SNPs (default $T \equiv 10000$) to ensure that estimates are accurate for potentially interesting SNPs.

[18] In practice the four conditional bdrs are estimated separately using new random subsets $t_j$ each time, but the procedure is simpler to explain in terms of a single subset.

The overall two-tailed bdr estimate is produced by integrating over the posterior probabilities that $\beta_j$ is positive, negative, or has unknown sign, obtained from the NUPE model estimates (denoted by $\acute{\Lambda}_{(+)}$, $\acute{\rho}_{(+)}$, etc):

$$\widehat{\mathrm{bdr}}_j = (\widehat{\mathrm{bdr}}_j|\beta_j > 0) + \widetilde{\mathrm{bdr}}_j|\beta_j > 0))\frac{\acute{\Lambda}_{(+)}(z_j)\acute{\rho}_{(+)}}{\acute{\Lambda}(z_j)} \tag{54}$$

$$+ (\widehat{\mathrm{bdr}}_j|\beta_j < 0) + \widetilde{\mathrm{bdr}}_j|\beta_j < 0))\frac{\acute{\Lambda}_{(-)}(z_j)\acute{\rho}_{(-)}}{\acute{\Lambda}(z_j)} \tag{55}$$

$$+ \frac{\acute{\Lambda}_u(z_j)(1 - \acute{\rho}_{(+)} - \acute{\rho}_{(-)})}{\acute{\Lambda}(z_j)}, \tag{56}$$

in which it is assumed that bdrs conditioned on SNP $j$ having an unknown sign (belonging to the $\acute{\Lambda}_u(z_j)$ distribution) are equal to one (i.e. all alt-SNPs have true effects equal or larger than SNP $j$).

##### Stage 6: Estimate tail area FDRs, BDRs and BFDRs

To obtain tail area FDR estimates for each SNP $i$ we use a standard approach of taking the mean fdr over SNPs with fdrs more extreme than SNP $i$:

$$\widehat{\mathrm{FDR}}_i = \frac{\sum_{\widehat{\mathrm{fdr}}_j \leqslant \widehat{\mathrm{fdr}}_i} \widehat{\mathrm{fdr}}_j}{\sum_j I(\widehat{\mathrm{fdr}}_j \leqslant \widehat{\mathrm{fdr}}_i)}, \tag{57}$$

and likewise for BDR:

$$\widehat{\mathrm{BDR}}_i = \frac{\sum_{\widehat{\mathrm{bdr}}_j \leqslant \widehat{\mathrm{bdr}}_i} \widehat{\mathrm{bdr}}_j}{\sum_j I(\widehat{\mathrm{bdr}}_j \leqslant \widehat{\mathrm{bdr}}_i)}. \tag{58}$$

BFDR estimates are then obtained using Equation 1:

$$\widehat{\mathrm{BFDR}}_i = \widehat{\mathrm{FDR}}_i + \widehat{\mathrm{BDR}}_i \times (1 - \widehat{\mathrm{FDR}}_i). \tag{59}$$

Simulation studies demonstrating the effectiveness of these estimates are described in the main text.

### 3 Prior splitting theory

##### Splitting priors into non-negative and non-positive mixtures

During NUPE/NUPE-diff estimation, we model the ratios[19] $\Lambda_{(+)}(z)/\Lambda_{(u)}(z)$ and $\Lambda_{(-)}(z)/\Lambda_{(u)}(z)$ as curves similar to squared-polynomials, but for which the derivates are always non-negative and non-positive respectively (Section 2, Stage 2). In Stage 3 the same model is applied to scaled effect estimate differences, $\delta$. Here we show why the resulting distribution models[20] $\Lambda_{(+)}(z)/W$ and $\Lambda_{(-)}(z)/W$ are conditional on $\beta > 0$ and $\beta < 0$ respectively, allowing the partitioning of the overall likelihood into mixtures with non-overlapping priors on $\beta$, which we term 'prior splitting'. Focussing first on the positive-effect distribution, based on the modelling assumptions provided in Section 2, $\Lambda_{(+)}(z)/W$ can be written as

$$\frac{\Lambda_{(+)}(z)}{W} = \int_{-\infty}^{\infty} f_x(z) g_{(+)}(x) dx, \tag{60}$$

where $x$ is expected value of $z$, the observed z-score, also its scaled true effect size $\beta/\hat{s}$, $g_{(+)}(x)$ is its unknown prior distribution function, and $f_x(z)$ is a normal distribution with mean $x$ and unit variance: $\frac{1}{\sqrt{2\pi}} e^{-(z-x)^2/2}$. As $\Lambda_{(u)}(z)/W = f_0(z)$, the ratio $\Lambda_{(+)}(z)/\Lambda_{(u)}(z)$ is

$$\frac{\Lambda_{(+)}(z)}{\Lambda_{(u)}(z)} \equiv R_{(+)}(z) = \int_{-\infty}^{\infty} e^{zx - x^2/2} g_{(+)}(x) dx, \tag{61}$$

which has the derivative

$$R'_{(+)}(z) = \int_{-\infty}^{\infty} x e^{zx - x^2/2} g_{(+)}(x) dx. \tag{62}$$

It can now be seen that whenever $g_{(+)}(x) > 0$ for any $x < 0$, one can find a $z < 0$ that is sufficiently negatively large that for any $y > 0$,

$$e^{zx - x^2/2} g_{(+)}(x) \gg e^{zy - y^2/2} g_{(+)}(y) \geqslant 0, \tag{63}$$

so that

$$x e^{zx - x^2/2} g_{(+)}(x) + y e^{zy - y^2/2} g_{(+)}(y) \ll 0 \tag{64}$$

[19]We illustrate the argument in this section with generic variables e.g. $z$, $\beta$ and $\hat{s}$, dropping the subscripts.

[20]Recall that $W$ is used to scale densities to give probability masses falling into z-bins of width $W$ (see Equation 22).

As such $R'_{(+)}(z)$ being negative anywhere is a necessary condition for $g_{(+)}(x) > 0$ when any $x < 0$. It follows that non-negativity of $R'_{(+)}(z)$ is a sufficient condition for $g(x) = 0$ when any $x < 0$, and as we have constrained $R'_{(+)}(z)$ to be non-negative everywhere this implies that $g_{(+)}(x) = 0$ for any $x < 0$.

The reverse implication, that $g_{(+)}(x) = 0$ for all $x < 0$ implies $R'_{(+)}(z) \geqslant 0$, is apparent in Equation 62, as the right hand side can only be negative when there is at least one $x < 0$ for which $g_{(+)}(x) > 0$. Therefore, $R'_{(+)}(z) \geqslant 0$ if and only if $g_{(+)}(x) = 0$ for all $x < 0$.

The same essential reasoning (with signs reversed) can be applied to $R_{(-)}(z)$. As its derivative is non-positive everywhere, neither a positive $z$ or positive $x$ can be found for which

$$xe^{zx-x^2/2}g_{(-)}(x) + ye^{zy-y^2/2}g_{(-)}(y) \gg 0, \tag{65}$$

for any $y$ where $y < 0$, therefore $g_{(-)}(x) = 0$ for any $x > 0$. Conversely, replacing $g_{(+)}(x)$ with $g_{(-)}(x)$ in Equation 62, one sees that the right hand side can only be positive when there is at least one $x > 0$ for which $g_{(-)}(x) > 0$. Therefore, $g_{(-)}(x) = 0$ for all $x > 0$ implies that $R'_{(-)}(z) \leqslant 0$, so $R'_{(-)}(z) \leqslant 0$ if and only if $g_{(-)}(x) = 0$ for all $x > 0$.

The theory we have presented is analogous to Tweedie's formula [Efron and Hastie, 2016, Robbins, 1956], which uses the derivative of a log-likelihood function to estimate the posterior expectation of $x$ conditioned on $z$. Our innovation is to constrain the derivative of the likelihood:null-likelihood ratio, allowing estimation of likelihoods with non-overlapping prior distributions of effects across either side of a given threshold, in this case zero.

#### Splitting from the zero-effect distribution

While the reasoning above demonstrates that the prior distributions $g_{(+)}(x)$ and $g_{(-)}(x)$ both equal zero for negative and positive $x$ respectively, they should also ideally be constrained to equal zero when $x = 0$. Assuming that $g_{(+)}(x) = 0$ for $x < 0$ as shown in the previous subsection, $R_{(+)}(z)$ decreases as $z \to (-\infty)$, due to $(zx - x^2/2)$ decreasing inside the exponent whenever $x$ is positive (see Equation 61). If it is also the case that $g_{(+)}(0) = 0$, $R_{(+)}(z) \to 0$ as $z \to (-\infty)$, because the exponent approaches zero for any $g_{(+)}(x) > 0$. This is not the case if $g_{(+)}(0) > 0$, as there will be a 'sill' at $g_{(+)}(0)$ in the function $R_{(+)}(z)$ as $z \to (-\infty)$, due to $e^{z0-0^2/2} = 1$. Therefore, $R_{(+)}(z) \to 0$ as $z \to (-\infty)$ if and only if $g_{(+)}(0) = 0$.

The condition is met as the expression inside the exponent on the right hand side of Equation 22 becomes dominated by the largest power of $z$ (when $n = D$ and $p = D$), so as $z \to (-\infty)$ the ratio approaches[21]

$$R_{(+)}(z) \to e^{z^{\frac{\Gamma_D \Gamma_D z^D z^D}{2D+1}}} \to 0. \tag{66}$$

This ensures that the prior $g_{(+)}(x)$ has zero probability at $x = 0$. Together with the reasoning in the previous subsection we are thus able to state that $g_{(+)}(x) = 0$ for all $x \leqslant 0$.

Reversing the logic shows that the right hand side of Equation 23 approaches zero as $z \to \infty$. Because $R_{(-)}(z) \to 0$ as $z \to \infty$ if and only if $g_{(-)}(0) = 0$, this ensures that $g_{(-)}(x) = 0$ for all $x \geqslant 0$.

[21]This assumes that $|\Gamma_D| > 0$ which should be the case, but if not the same reasoning can be applied to $D - 1$, or to whichever is the largest term in the polynomial which for which $|\Gamma_n| > 0$.

**Figure A3:** Fitted NUPE-diff model. Histograms show effect estimate differences between SNPs with positive true effects and randomly selected alt SNPs (a-b), and between SNPs with negative true effects and randomly selected alt SNPs (c-d). Green and turquoise curves are the fitted distributions for negative and positive true effect differences respectively. Null differences, for which there is insufficient power to distinguish from zero, are distributed according to the blue curve, and the fitted marginal mixture distribution is red.

(a) T1D - Differences between estimates for positive effect SNPs and random alt SNPs

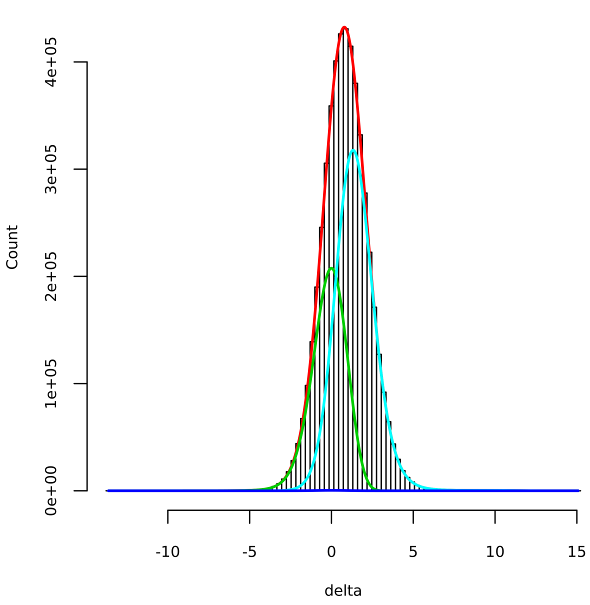

(b) ATD - Differences between estimates for positive effect SNPs and random alt SNPs

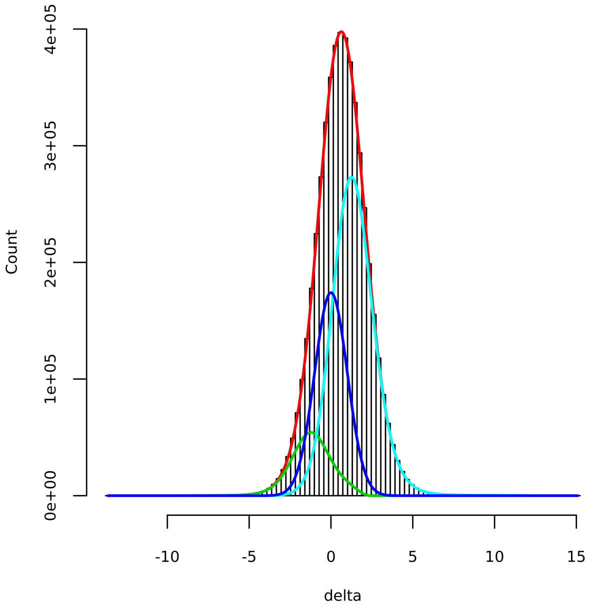

(c) T1D - Differences between estimates for negative effect SNPs and random alt SNPs

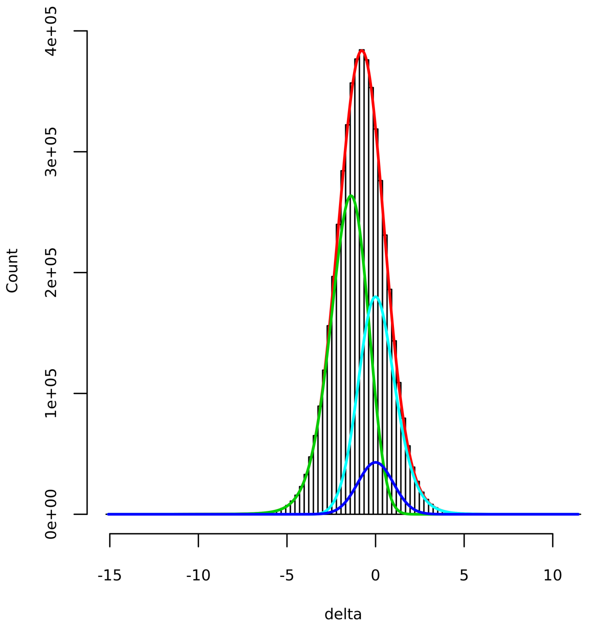

(d) ATD - Differences between estimates for negative effect SNPs and random alt SNPs

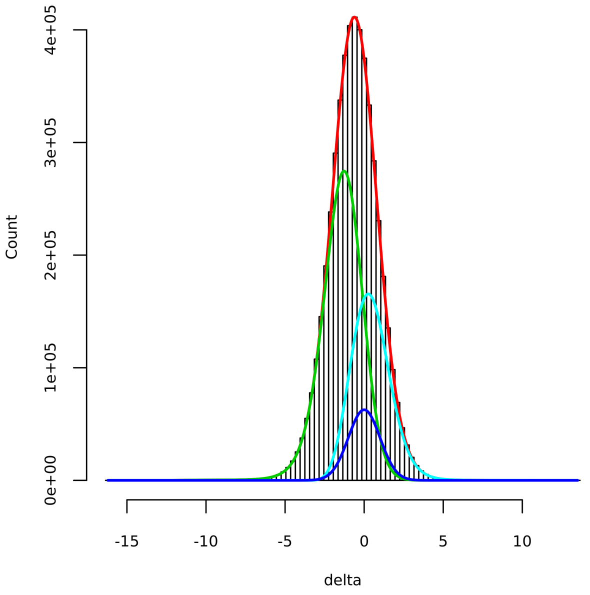
