## Supplementary Note: T1DGC Membership for "Enhanced genetic analysis of type 1 diabetes by selecting variants on both effect size and significance, and by integration with autoimmune thyroid disease"

**Asia-Pacific Network:** Tracey Baskerville (Mater Children's Hospital, Australia); Nines Bautista (Institute for Study on Diabetes Foundation, Philippines); Eesh Bhatia (Sanjay Gandhi Postgraduate Institute, India); Vijayalakshmi Bhatia (Sanjay Gandhi Postgraduate Institute, India); Kamaruzaman Bin Hasan (National University of Malaysia Hospital, Malaysia); Francois Bonnici (University of Cape Town, South Africa); Thomas Brodnicki (Walter & Eliza Hall Institute of Medical Research, Australia); Brian Browning (The University of Auckland, New Zealand); Fergus Cameron (Royal Children's Hospital, Australia); Katharee Chaichanwatanakul (Mahidol University, Thailand); Pik To Cheung (Queen Mary Hospital, Hong Kong); Peter Colman[2,5,11,15] (Walter & Eliza Hall Institute of Medical Research, Australia); Andrew Cotterill (Mater Children's Hospital, Australia); Jenny Couper (Women's and Children's Hospital, Australia); Patricia Crock (John Hunter Children's Hospital, Australia); Ric Cutfield (North Shore Hospital, New Zealand); Tim Davis (Fremantle Hospital, Australia); Paul Dixon (Diabetes Lifestyle Centre, New Zealand); Kim Donaghue (Children's Hospital at Westmead, Australia); Katrina Dowling[4] (Australian Red Cross Blood Service, Australia); Paul Drury (Auckland Diabetes Centre, New Zealand); Sarah Dye (Western Australia Institute for Medical Research, Australia); Shane Gellert[2] (The Royal Melbourne Hospital, Australia); Rohana Abdul Ghani (National University of Malaysia Hospital, Malaysia); Ristan Greer (University of Queensland, Australia); Xueyao Han (Peking University People's Hospital, China); Len Harrison (Walter & Eliza Hall Institute of Medical Research, Australia); Nick Homatopoulos[4] (Australian Red Cross Blood Service, Australia); Linong Ji (Peking University People's Hospital, China); Tim

Jones (Princess Margaret Hospital for Children, Australia); Loke Kah Yin (Children's Medical Institute, Singapore); Nor Azmi Kamaruddin (National University of Malaysia Hospital, Malaysia); Uma Kanga (All India Institute of Medical Sciences, India); Alok Kanungo (Cuttack Diabetes Research Foundation, India); Gurvinder Kaur (All India Institute of Medical Sciences, India); Betty Kek (Children's Medical Institute, Singapore); Simon Knowles[4] (Australian Red Cross Blood Service, Australia); Jeremy Krebs (The Diabetes Centre, New Zealand); Neeraj Kumar (All India Institute of Medical Sciences, India); Yann-Jinn Lee[7] (Mackay Memorial Hospital, Taiwan); Xiaoying Li (Shanghai Jiao-Tong University, China); Supawadee Liktimaskul (Mahidol University, Thailand); Margaret Lloyd (Children's Hospital at Westmead, Australia); Amanda Loth[5,10] (Walter & Eliza Hall Institute of Medical Research, Australia); Anthony Louey[3,4] (Australian Red Cross Blood Service, Australia); Narinder Mehra[8] (All India Institute of Medical Sciences, India); Tony Merriman (University of Otago, New Zealand); Liu Min (Beijing Children's Hospital, China); Grant Morahan[1,9,12,14,15] (Western Australia Institute for Medical Research, Australia); Robert Moses (Illawarra Diabetes Services, Australia); Grant Mraz[4] (Australian Red Cross Blood Service, Australia); Rinki Murphy (Auckland Diabetes Centre, New Zealand); Ian Nicholson[4] (Australian Red Cross Blood Service, Australia); Araceli Panelo (Institute for Studies on Diabetes Foundation, Philippines); Perlita Poh[2] (Royal Melbourne Hospital, Australia); Gareth Price (Mater Medical Research Institute, Australia); Nirubasini Ratnam (Princess Margaret Hospital for Children, Australia); Carani Sanjeevi[6] (Karolinska Hospital, Sweden); Saikiran Sedimbi (Karolinska Hospital, Sweden); Shuixian Shen (Fudan University, China); Goh Siok Ying (The Children's Medical Institute, Singapore); Brian Tait[3,4,5,6] (Australian Red

Cross Blood Service, Australia); Nikhil Tandon (All India Institute of Medical Sciences, India); Allison Thomas (Walter & Eliza Hall Institute of Medical Research, Australia); Mike Varney[3,4] (Australian Red Cross Blood Service, Australia); Praewvarin Weerakulwattana (Mahidol University, Thailand); Jinny Willis (Christchurch Hospital, New Zealand)

**European Network:** Elvis Abang Akwo (Yaounde Central Hospital, Cameroon); Lotte Albret[5,10] (Hagedorn Research Institute and Steno Diabetes Center, Denmark); Francisco Ampudia-Blasco (Clinic University Hospital Valencia, Spain); Jesus Argente (Hospital Infantil Universitario Nino Jesus, Spain); Magdalena Avbelj (University Children's Hospital, Slovenia); Gulja Babadjanova (Moscow State Medical University, Russia); Klaus Badenhoop[14] (University Clinic Frankfurt/Main, Germany); Tadej Battelino (University Children's Hospital, Slovenia); Georg Beilhack[14] (University of Ulm, Germany); Regine Bergholdt (Hagedorn Research Institute and Steno Diabetes Center, Denmark); Polly Bingley[2,5] (University of Bristol, United Kingdom); Bernhard Boehm[4,5,14] (Ulm University, Germany); Jo Bolidson[2] (University of Bristol, United Kingdom); Kerstin Brismar (Karolinska Hospital, Sweden); Caroline Brorsson[14] (Hagedorn Research Institute and Steno Diabetes Center, Denmark); Joyce Carlson[3,5] (University Hospital MAS, Sweden); Luis Castano (Hospital de Cruces, Spain); Kyla Chandler[2] (University of Bristol, United Kingdom); Valentino Cherubini (Salesi Hospital, Italy); Ondrej Cinek (Motol University Hospital, Czech Republic); Elisa Cipponeri (University Campus Bio-Medico, Italy); Raquel Corripio Collado (Consorci Sanitari Parc Tauli, Spain); Alberto de Leiva (Hospital Sant Pau, Spain); Iveta Dzivite (University

Children's Hospital, Latvia); Ana Fagulha (University Hospital, Portugal); Merce Fernandez Balcells (Hospital Trueta, Spain); Beatriz Garcia Cuartero (Hospital Severo Ochoa, Spain); Concepcion Garcia Lacalle (Hospital Severo Ochoa, Spain); Cristian Guja (Institute of Diabetes, Nutrition & Metabolic Diseases, Romania); Pilar Gutiérrez (Hospital Universitario de Getafe, Spain); Alona Hamou (Schneider Children's Medical Center of Israel, Israel); Erifili Hatziagelaki (University of Athens, Greece); Simon Heath[7] (Centre National de Genotypage, France); Kaire Heilman (Tartu University Children's Hospital, Estonia); Wolfgang Helmberg[5,7] (Medical University Graz, Austria); Orna Hermon (Schneider Children's Medical Center of Israel, Israel); Marta Hernandez (Hospital Universitari Arnau de Vilanova and Hospital Universitario de Canarias, Spain); Iris Holzheu[4] (Ulm University, Germany); Nora Hosszufalusi (Semmelweis University, Hungary); Jorma Ilonen (University of Turku, Finland); Constantin Ionescu-Tirgoviste (Institute of Diabetes, Romania); Jesper Johannesen (Steno Diabetes Center, Denmark); Cecile Julier[1,9,12,14] (Centre National de Genotypage, France); Heinrich Kahles[14] (Klinikum der J.W. Goethe-Universität, Germany); Ida Kinalska (Medical University of Bialystok, Poland); Mikael Knip (University of Helsinki, Finland); Ingrid Kockum[7,14] (Karolinska Hospital, Sweden); Eija Kojo (University of Helsinki, Finland); Olga Kordonouri (Children's Hospital auf der Bult, Germany); Adam Kretowski (Medical University of Bialystok, Poland); Dora Krikovszky (Semmelweis University, Hungary); Angelika Kurkhaus[4] (Ulm University, Germany); Madiusz Kuzmicki (Medical University of Bialystok, Poland); Eva Lavant[3] (University Hospital MAS, Sweden); Anna Long[2] (University of Bristol, United Kingdom); Johnny Ludvigsson (University Hospital, Sweden); Laszlo Madacsy (Semmelweis University, Hungary); Katarzyna Maliszewska

(Medical University of Bialystok, Poland); Mara Marga (P. Stradins University Hospital, Latvia); Marissa Penna Martinez (University Clinic Frankfurt/Main, Germany); Didac Mauricio[6] (Hospital Universitari Arnau de Vilanova and Hospital Sant Pau, Spain); Gertrud Mazurkievicz[4] (Ulm University, Germany); Jorn Nerup[1,15] (Steno Diabetes Center, Denmark); Antanas Norkus (Institute of Endocrinology of Kaunas University of Medicine, Lithuania); Francisco Javier Novoa Mogollon (Hospital Universitario Insular, Spain); Anna Okruszko (Medical University of Bialystok, Poland); Chiara Pettinari (Salesi Hospital, Italy); Moshe Phillip (Schneider Children's Medical Center of Israel, Israel); Valdis Pirags (P. Stradins University Hospital, Latvia); Flemming Pociot[1,11,12,14,15] (Hagedorn Research Institute and Steno Diabetes Center, Denmark); Paolo Pozzilli (University Campus Bio-Medico, Italy); Radu Racasan (University Clinic Frankfurt/Main, Germany); Klemens Raile (Vircow Clinic Charité Berlin, Germany); Rebecca Rappner[3] (University Hospital MAS, Sweden); Maria Jesus Rodriguez Troyano (University Hospital of Las Palmas de Gran Canaria, Spain); Bart O. Roep (Leiden University Medical Center, Netherlands); Saba Rokni[2] (Southmead Hospital, United Kingdom); Silke Rosinger[4] (Ulm University, Germany); Oscar Rubio-Cabezas (Hospital Infantil Universitario Nino Jesus, Spain); Christa Ruckgaber[4] (Ulm University, Germany); Ilhan Satman (Istanbul University, Turkey); Edith Schober (University Children's Hospital, Austria); Jochen Seufert (Medizinsche Poliklinik der Universität, Germany); Rosi Sing[4] (Ulm University, Germany); Jan Skrha (Faculty of Medicine 1, Czech Republic); Eugene Sobngwi (Central National Obesity Centre and Hospital of Diabetes Endocrine, Cameroon); Michelle Somerville[2] (University of Bristol, United Kingdom); Giatgen Spinas[8] (University Hospital, Switzerland); Zdenek Sumnik

(University Hospital Motol, Czech Republic); Vallo Tilmann (Tartu University Children's Hospital, Estonia); Dag Undlien[6] (University of Oslo, Norway); Vaidotas Urbanavicius (Vilnius University Hospital, Lithuania); Bart Van der Auwera[8] (Vrije Universiteit Brussel, Belgium); Federico Vasquez San Miguel (Hospital de Cruces, Spain); Andriani Vazeo-Gerasimidi (Diabetes Center P&A Kyriakou Children's Hospital, Greece); Dzilda Velickiene (Institute of Endocrinology of Kaunas University of Medicine, Lithuania); Ana Wägner[5,10] (University Hospital of Las Palmas de Gran Canaria, Spain, and Steno Diabetes Center, Denmark); Markus Walter (Diabetes Research Institute, Germany); Alistair Williams[2] (University of Bristol, United Kingdom); Anette Ziegler (Diabetes Research Institute, Germany)

**North American Network:** Matthew Agleham[3] (Roche Molecular Systems, United States); Alan Aldrich[5,10] (Benaroya Research Institute, United States); Ramin Alemzadeh (Medical College of Wisconsin, United States); Chester Alper (Immune Disease Institute, United States); Theresa Aly (Barbara Davis Center for Childhood Diabetes, United States); Dimitris Anastassiou (Columbia University, United States); Shaily Arora[3] (Children's Hospital Oakland Research Institute, United States); Audrey Austin (Children's National Medical Center, United States); Dorothy Becker (Rangos Research Center, United States); Christophe Benoist (Joslin Diabetes Center, United States); Noureddine Berka[6] (Calgary Laboratory Services, Canada); Suruchi Bhatia (Oakland Children's Hospital Research Center, United States); Persia Bonella[3] (Roche Molecular Systems, United States); Nunzio Bottini[14] (University of Southern California, United States); Sean Boyle[3] (Roche Molecular Systems, United States); Jeanah Braden

(Children's Hospital Oakland Research Institute, United States); Barry Brady (Arkansas Children's Hospital, United States); Wendy Brickman (Children's Memorial Hospital, United States); Richard Christensen (Humphreys Diabetes Center, United States); Patrick Concannon[1,9,12,14] (University of Virginia, United States); Robert Couch (University of Alberta, Canada); Debra Counts (University of Maryland, United States); Jill Crandall (Albert Einstein College of Medicine, United States); Mark Daniels (Children's Hospital of Orange County, United States; Larry Dolan (Cincinnati Children's Hospital Medical Center, United States); David Donaldson (Utah Diabetes Center, United States); Alessandro Doria[6] (Joslin Diabetes Center, United States); George Eisenbarth[2,5,13,14] (Barbara Davis Center for Childhood Diabetes, United States); James Elder (University of Michigan, United States); Rita El-Hajj (Main Line Health Heart Center, United States); Henry Erlich[1,3,5,13,14] (Roche Molecular Systems, United States); Pamela Fain (Barbara Davis Center for Childhood Diabetes, United States); Anna Lisa Fear[3] (Children's Hospital Oakland Research Institute, United States); Robert Ferry (The University of Texas Health Science Center at San Antonio, United States); Rosanna Fiallo-Scharer (Barbara Davis Center for Childhood Diabetes, United States); Daniel Geraghty (Fred Hutchinson Cancer Research Center, United States); Soumitra Ghosh[6] (Medical College of Wisconsin, United States); Steven Gitelman (University of California at San Francisco, United States); Michelle Godwin[4] (Fred Hutchinson Cancer Research Center, United States); Robin Goland (Naomi Berrie Diabetes Center, United States); Nathan Goodman[7] (Institute for Systems Biology, United States); Greg Goodwin (Joslin Diabetes Center, United States); Jenna Gravely[4] (Fred Hutchinson Cancer Research Center, United States); Carla Greenbaum[8,11,15] (Benaroya Research Institute,

United States); Chelsea Gudgeon[4] (Fred Hutchinson Cancer Research Center, United States); Fred Gunville (Billings Clinic, United States); William Hagopian[11] (University of Washington, United States); Hakon Hakonarson (Children's Hospital of Philadelphia, United States); John Hansen[4,5] (Fred Hutchinson Cancer Research Center, United States); Kimberly Harrington[4] (Fred Hutchinson Cancer Research Center, United States); Jeanne Hassing (Sacred Heart, United States); Wendy Hilliker[4] (Fred Hutchinson Cancer Research Center, United States); Robert Hoffman (Ohio State University, United States); Erin Hulbert (Institute for Systems Biology, United States); Roberto Izquierdo (SUNY Upstate Medical University, United States); Nicholas Jospe (University of Rochester, United States); Kevin Kaiserman (Children's Hospital Los Angeles, United States); Francine Kaufman (Children's Hospital Los Angeles, United States); Samuel Kim[3] (Roche Molecular Systems, United States); Erin Kloos[4] (Fred Hutchinson Cancer Research Center, United States); Roman Kosoy (Benaroya Research Institute, United States); James Lane (University of Nebraska, United States); Julie Lane[3] (Children's Hospital Oakland Research Institute, United States); Jean Lawrence (Kaiser Permanente, United States); Claresa Levetan (Main Line Health Heart Center, United States); Phil Levin (MODEL Clinical Research, United States); Rebecca Lipton (University of Chicago, United States); John Lonsdale (Human Biological Data Interchange, United States); Victoria Magnuson (Children's Hospital of Wisconsin, United States); Jennifer Marks (University of Miami, United States); Beth Mayer-Davis (University of South Carolina, United States); Robert McEvoy (Children's Hospital of Minnesota, United States); Richard McIndoe[7] (Medical College of Georgia, United States); Lesley Merkle[4] (Fred Hutchinson Cancer Research Center, United States); Daniel Metzger (BC

Children's Hospital, Canada); Dongmei Miao[2] (Barbara Davis Center for Childhood Diabetes, United States); Eric Mickelson[4] (Fred Hutchinson Cancer Research Center, United States); Priscilla Moonsamy[3] (Roche Molecular Systems, United States); Wayne Moore (Children's Mercy Hospital, United States); Antoinette Moran (University of Minnesota, United States); Janelle Noble[3,5,13,14] (Children's Hospital Oakland Research Institute, United States); Gary Olsem[4] (Fred Hutchinson Cancer Research Center, United States); Suna Onengut-Gumuscu[14] (University of Virginia, United States); Tihamer Orban (Joslin Diabetes Center, United States); Craig Orlowski (University of Rochester Medical Center, United States); Andrew Paterson (University of Toronto, Canada); Massimo Pietropaolo (University of Michigan Medical School, United States); Catherine Pihoker (Children's Hospital and Regional Medical Center, United States); Constantin Polychronakos[11,14] (McGill University Health Center, Canada); Jeff Post[3] (Roche Molecular Systems, United States); Daniel Postellon (Helen DeVos Children's Hospital, United States); Alberto Pugliese[9,14] (University of Miami, United States); HuiQi Qu[14] (Montreal Children's Hospital, Canada); Teresa Quattrin (Women and Children's Hospital of Buffalo, United States); Mark Rappaport (Pediatric Endocrine Associates, United States); Philip Raskin (University of Texas Southwestern Medical Center, United States); Heather Risbeck[4] (Fred Hutchinson Cancer Research Center, United States); Henry Rodriguez (Riley Hospital for Children, United States); Luisa Rodriguez (Baylor College of Medicine, United States); Michelle Rogers[4] (Fred Hutchinson Cancer Research Center, United States); Leticia Rubalcava (Children's Hospital Oakland Research Institute, United States); Bill Russell (Vanderbilt University, United States); Desmond Schatz (University of Florida, United States); Carla Scott (University of Texas

Health Science Center at San Antonio, United States); Jin-Xiong She[14] (Medical College of Georgia, United States); Heather Shilling (Benaroya Research Institute, United States); Dorothy Shulman (University of South Florida, United States); Leslie Soyka (University of Massachusetts Memorial Center, United States); Phyllis Speiser (Schneider Children's Hospital, United States); Harold Starkman (Atlantic Health System, United States); Andrea Steck[14] (Barbara Davis Center for Childhood Diabetes, United States); Sarah Stender (University of Tennessee, United States); Lorraine Stratton (University of Arizona, United States); Daniel Sur[3] (Roche Molecular Systems, United States); Shayne Taback (University of Manitoba, United States); Kathryn Thrailkill (Arkansas Children's Hospital, United States); Ellen Toth (University of Alberta, Canada); Patricia Trymbiski (Doylestown Hospital, United States); Eva Tsalikian (University of Iowa, United States); Katherine Vertachnik[4] (Fred Hutchinson Cancer Research Center, United States); Jack Wahlen (Endocrine Research Specialists, United States); Xujing Wang (Max McGee National Research Center of Juvenile Diabetes, United States); Sandra Weber (Greenville Hospital System, United States); Diane Wherrett (Hospital for Sick Children, Canada); Steven Willi (Children's Hospital of Philadelphia, United States); Darrell Wilson (Stanford University, United States); Jerry Youkey (Greenville Hospital System, United States); Neal Young (National Institutes of Health, United States); Liping Yu[2] (Barbara Davis Center for Childhood Diabetes, United States); Lue Ping Zhao (Fred Hutchinson Cancer Research Institute, United States); Donald Zimmerman (Children's Memorial Hospital, United States)

**United Kingdom Network:** Ellen Adlem[4] (University of Cambridge, United Kingdom); James Allen[4] (University of Cambridge, United Kingdom); Jeffrey Barrett (WT Sanger Institute, United Kingdom); Judy Brown[4] (University of Cambridge, United Kingdom); Oliver Burren[4] (University of Cambridge, United Kingdom); Pamela Clarke[4] (University of Cambridge, United Kingdom); David Clayton[4] (University of Cambridge, United Kingdom); Gillian Coleman[4] (University of Cambridge, United Kingdom); Jason Cooper[4] (University of Cambridge, United Kingdom); Francesco Cucca[6] (University of Sassari, United Kingdom); Lucy Davison (University of Cambridge, United Kingdom); Kate Downes (University of Cambridge, United Kingdom); Simon Duley[4] (University of Cambridge, United Kingdom); David Dunger[11] (University of Cambridge, United Kingdom); Laura Esposito (University of Cambridge, United Kingdom); Vin Everett[4] (University of Cambridge, United Kingdom); Sarah Field (University of Cambridge, United Kingdom); Jason Hafler (University of Cambridge, United Kingdom); Matthew Hardy[4] (University of Cambridge, United Kingdom); Deborah Harrison[4] (University of Cambridge, United Kingdom); Inge Harrison[4] (University of Cambridge, United Kingdom); Steve Hawkins[4] (University of Cambridge, United Kingdom); Barry Healy[4] (University of Cambridge, United Kingdom); Simon Hood[4] (University of Cambridge, United Kingdom); Simon Howell[8] (King's College, United Kingdom); Joanna Howson (University of Cambridge, United Kingdom); Meeta Maisuria[4] (University of Cambridge, United Kingdom); William Meadows[4] (University of Cambridge, United Kingdom); Trupti Mistry[4] (University of Cambridge, United Kingdom); Sergey Nezhenstsev (University of Cambridge, United Kingdom); Sarah Nutland[4,5] (University of Cambridge, United Kingdom); Nigel Ovington[4] (University of Cambridge, United Kingdom);

Vincent Plagnol (University of Cambridge, United Kingdom); Dan Rainbow (University of Cambridge, United Kingdom); Kara Rainbow (University of Cambridge, United Kingdom); Srilakshmi Raj (University of Cambridge, United Kingdom); Helen Schuilenburg[4] (University of Cambridge, United Kingdom); Anna Simpson[4] (University of Cambridge, United Kingdom); Luc Smink[7] (University of Cambridge, United Kingdom); Debbie Smyth (University of Cambridge, United Kingdom); Helen Stevens[4] (University of Cambridge, United Kingdom); Niall Taylor[4] (University of Cambridge, United Kingdom); John Todd[1,4,12,14,15] (University of Cambridge, United Kingdom); Jaakko Tuomilehto (National Public Health Institute, Finland); Neil Walker[4,5] (University of Cambridge, United Kingdom); Linda Wicker (University of Cambridge, United Kingdom); Barry Widmer[4] (University of Cambridge, United Kingdom); Mark Wilson[4] (University of Cambridge, United Kingdom); Heather Withers[5,10] (University of Cambridge, United Kingdom); Jennie Yang (University of Cambridge, United Kingdom)

**Coordinating Center:** Mark Brown (Wake Forest University Health Sciences, United States); Wei-Min Chen (University of Virginia, United States); Arnetta Crews (Wake Forest University Health Sciences, United States); Jason Griffin (Wake Forest University Health Sciences, United States); Mark Hall[8] (Wake Forest University Health Sciences, United States); Teresa Harnish (Wake Forest University Health Sciences, United States); John Hepler (Wake Forest University Health Sciences, United States); Joan Hilner[5,8,10] (Wake Forest University Health Sciences, United States); Nancy King[8] (Wake Forest University Health Sciences, United States); Kurt Lohman (Wake Forest University Health Sciences, United States); Lingyi Lu (Wake Forest University Health Sciences,

United States); Josyf Mychaleckyj[5,7] (University of Virginia, United States); Jay Nail (Wake Forest University Health Sciences, United States); Letitia Perdue[5,10] (Wake Forest University Health Sciences, United States); June Pierce (Wake Forest University Health Sciences, United States); David Reboussin[5,6] (Wake Forest University Health Sciences, United States); Stephen Rich[1,12,14] (University of Virginia, United States); Scott Rushing (Wake Forest University Health Sciences, United States); Michele Sale (University of Virginia, United States); Elizabeth Sides[5,10] (Wake Forest University Health Sciences, United States); Beverly Snively[11] (Wake Forest University Health Sciences, United States); Hoa Teuschler (Wake Forest University Health Sciences, United States); Goodrich Theil (Wake Forest University Health Sciences, United States); Lynne Wagenknecht (Wake Forest University Health Sciences, United States); Dustin Williams (Wake Forest University Health Sciences, United States)

**Project Office:** Beena Akolkar[1,5,6,9,12] (National Institute of Diabetes and Digestive and Kidney Diseases/National Institutes of Health, United States); Catherine McKeon[8] (National Institute of Diabetes and Digestive and Kidney Diseases/National Institutes of Health, United States); Concepcion Nierras[9] (Juvenile Diabetes Research Foundation International, United States); Elizabeth Thomson[8] (National Human Genome Research Institute/National Institutes of Health, United States)

**Other Contributors:** David Altshuler (Whitehead Institute for Biomed Research, United States); Kinman Au[14] (Medical College of Georgia, United States); Steve Bain[14] (University of Wales Swansea, United Kingdom); Lisa Barcellos[13] (University of

California at Berkeley, United States); Sandra Barral[13] (Rockefeller University, United States); Tim Becker[13] (Karolinska Institute, Sweden); Farren Briggs[13] (University of California at Berkeley, United States); Paola Bronson[13] (University of California at Berkeley, United States); Mark Daly[7,13] (Massachusetts General Hospital, United States); Paul de Bakker[13] (Massachusetts General Hospital, United States); Panos Deloukas[13] (Wellcome Trust Sanger Institute, United Kingdom); Bernie Devlin[13] (University of Pittsburgh, United States); Morten Chrisoph Eike[13,14] (Institute of Immunology, Norway); Leigh Field[14] (University of British Columbia, Canada); Stacey Gabriel (Broad Institute of MIT and Harvard, United States); Nikhil Garge[14] (Medical College of Georgia, United States); Silvana Gaudieri[13] (Murdoch University, Australia); Ben Goldstein[13] (University of California at Berkeley, United States); Clara Gorodezky (INDRE SSA, Mexico); Sara Hamon[13] (Rockefeller University, United States); Chungsheng He[13] (Rockefeller University, United States); Joanna Howson[4,13,14] (University of Cambridge, United Kingdom); Keith Humphreys[13] (Karolinska Institute, Sweden); Ian James[13] (Murdoch University, Australia); Mark Lathrop[13] (Centre National de Genotypage, France); Benedicte Alexandra Lie[13] (University of Oslo, Norway); Dawei Li[13] (Rockefeller University, United States); Steven Mack[13] (Roche Molecular Systems, United States); Ralph McGinnis[13] (Wellcome Trust Sanger Institute, United Kingdom); Elizabeth McKinnon[13] (Murdoch University, Australia); William McLaren[13] (Wellcome Trust Sanger Institute, United Kingdom); David Nolan[13] (Murdoch University, Australia); Marita Olsson[13] (Karolinska Institute, Sweden); Jurg Ott[13] (Rockefeller University, United States); David Owerbach (Baylor College of Medicine, United States); Chris Patterson[14] (Queen's University Belfast, United Kingdom); Robert Podolsky[14] (Medical

College of Georgia, United States); Patricia Ramsay[13] (University of California at Berkeley, United States); Venkatesh Rangantah[13] (Wellcome Trust Sanger Institute, United Kingdom); Neil Risch[13] (University of California at San Francisco, United States); Kjersti Skjold Ronningen[14] (Norwegian Institute of Public Health, Norway); Xiarong Shao[13] (University of California at Berkeley, United States); Richard Single[13] (University of Vermont, United States); Michael Steffes[5] (University of Minnesota, United States); Glenys Thomson[13] (University of California at Berkeley, United States); Ana Maria Valdes[5,13] (Lartech, Italy); Claire Vandiedonck[13] (Wellcome Trust Centre for Human Genetics, United Kingdom); Pam Whittaker (Wellcome Trust Sanger Institute, United Kingdom); Qingrun Zhang[13] (Beijing Institute of Genomics, China)

**Study Roles:** [1]Steering Committee, [2]Autoantibody Laboratory, [3]HLA Genotyping Laboratory, [4]Network DNA Repository, [5]Quality Control Committee, [6]Access Committee, [7]Bioinformatics Committee, [8]Ethical Legal and Social Issues Committee, [9]Molecular Technology Committee, [10]Network Coordinators Committee, [11]Phenotyping and Recruitment Committee, [12]Publications and Presentations Committee, [13]MHC Fine Mapping Working Group, [14]Rapid Response Working Group, [15]Network Principal Investigator
